# Size-dependent protein segregation creates a spatial switch for Notch signaling and function

**DOI:** 10.1101/2020.06.28.176560

**Authors:** Minsuk Kwak, Kaden M. Southard, Woon Ryoung Kim, Nam Hyeong Kim, Ramu Gopalappa, Minji An, Hyun Jung Lee, Min K. Kang, Seo Hyun Choi, Justin Farlow, Anastasios Georgakopoulos, Nikolaos K. Robakis, Matthew L. Kutys, Daeha Seo, Hyeong Bum Kim, Yong Ho Kim, Jinwoo Cheon, Zev J. Gartner, Young-wook Jun

## Abstract

Aberrant cleavage of Notch by γ-secretase is implicated in numerous diseases, but how cleavage is regulated in space and time is unclear. Here, we report that cadherin-based adherens junctions (cadAJs) are sites of high cell-surface γ-secretase activity, as well as sites of constrained physical space that excludes γ-secretase substrates having large extracellular domains (ECDs) like Notch. ECD shedding initiates drastic spatial relocalization of Notch to cadAJs, allowing enzyme-substrate interactions and downstream signaling. Spatial mutations by adjusting the ECD size or the physical constraint alter signaling. Dysregulation of this spatial switch promotes precocious differentiation of ventricular zone neural progenitor cells in vivo. We show the generality of this spatial switch for amyloid precursor protein proteolysis. Thus, cadAJs create spatially distinct biochemical compartments regulating cleavage events involving γ-secretase and preventing aberrant activation of receptors.

**One Sentence Summary:** Notch cleavage by γ-secretase is regulated through dynamic spatial control of receptors, adhesion molecules, and activating proteases

---

Notch is a highly conserved mediator of contact-dependent cell-cell communication, which orchestrates diverse functions in metazoans (*1, 2*). Tight control of Notch signal activation is essential for many developmental processes, while dysregulation of Notch activation can cause severe disease including developmental, neurological, and immunological disorders and cancer (*1–4*). Accordingly, to enable precise signal regulation, receptor activation occurs through multiple steps, independently gated by chemical (e.g., ligand-receptor interactions, posttranslational modifications) and mechanical cues (*1, 2, 5–9*). However, many signaling processes involving physical contact between two cells – so called juxtacrine signaling processes – are also regulated by spatial cues (*10–12*). As exemplified by the kinetic segregation model in immune cells (*10, 12, 13*), spatial rearrangements of these signaling molecules modify their physical and biochemical environments to facilitate receptor activation. Notch, which also mediates signal exchange by physical contact, is subjected to similar structural and spatial constraints (*14, 15*). To test whether Notch signaling is regulated by these spatial rearrangements, we investigated the spatial dynamics of Notch and its signaling partners at cellular interfaces. Specifically, we focused on five key signaling molecules – Notch, delta-like ligand 1 (Dll1), ADAM 10/17, and γ-secretase. We analyze their dynamics relative to cadherin-based adherens junctions (cadAJs): the cellcell adhesions that initiate many structural and spatial changes by bringing and holding opposing membranes together (*16–18*) (**Fig. 1A**).

**Figure 1.**
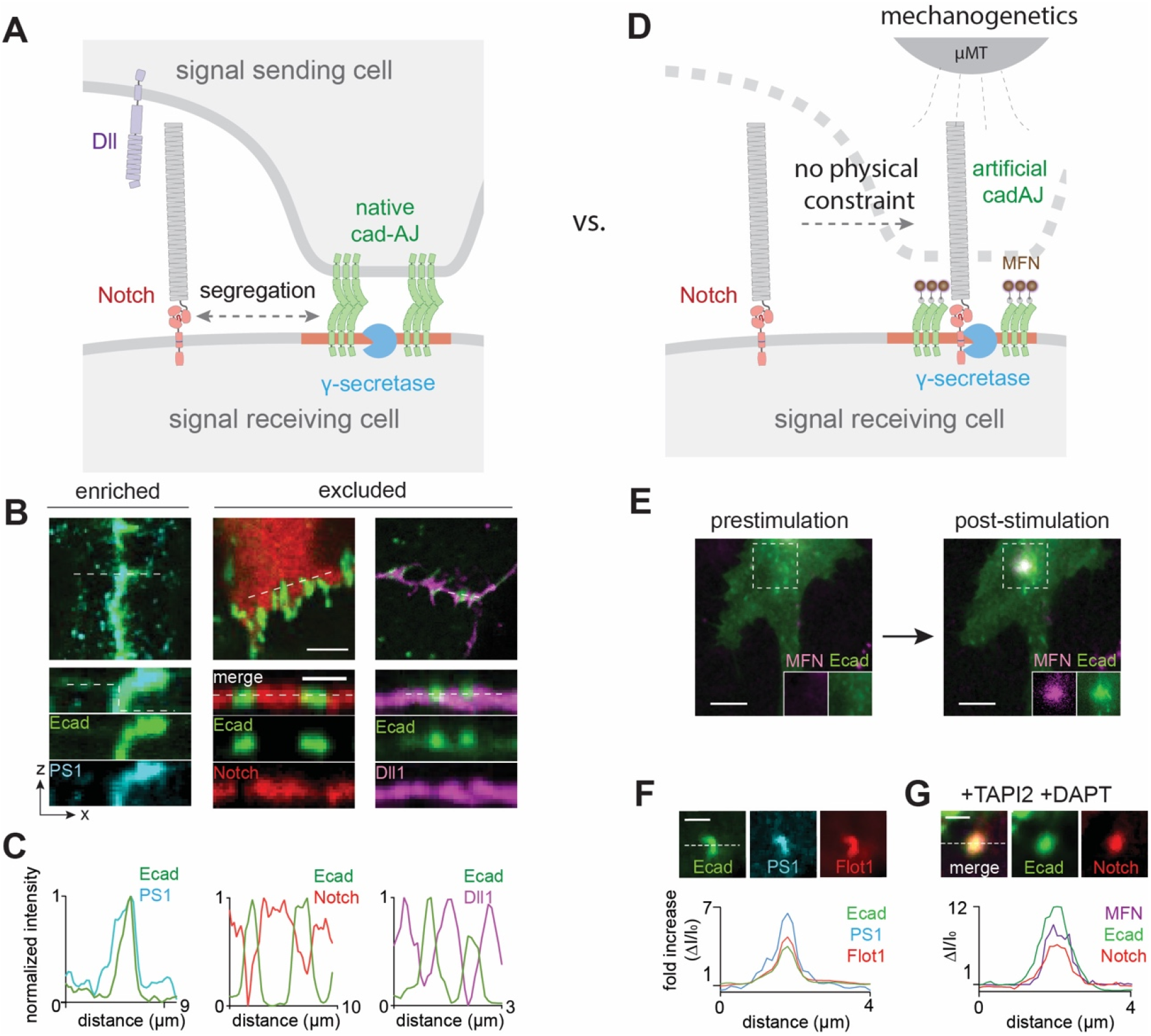
Spatial segregation of Notch receptors and ligands from cadAJs prevents their interactions with γ-secretase. (**A)** A schematic showing spatial distribution of Notch signaling components at the juxtaposed cell-cell interface. (**B)** Representative confocal fluorescence images showing presenilin-1 (PS1), Notch1, and Dll1 distributions relative to E-cadherin-based cadherin junction (cadAJ). (top) A maximum projection image. Scale bar, 5 μm. (bottom) z-resliced images. Scale bar, 3 μm. (**C)** Line profiles quantifying fluorescence from E-cadherin (green), PS1 (cyan), Notch (red), and Dll1 (magenta) along a representative section of the cadAJ (a white dashed line in z-resliced images of panel (B)). (**D**) A schematic showing mechanogenetic interrogation of γ-secretase and Notch distribution relative to the artificial cadAJs. Artificial cadAJs were formed by clustering Ecad-GFP labeled with magnetofluorescent nanoparticles (MFNs) by application of an external micromagnetic tweezer (μMT). DAPT was used to inhibit γ-secretase activity. **(E)** Epifluorescence images showing the formation of an artificial cadAJ by mechanogenetics. After stimulation by μMT, vivid MFN and E-cadherin signals at the magnetic focus were seen, indicating formation of cadAJs. Scale bar, 5 μm. **(F)** Confocal fluorescence images of E-cadherin, PS1, and Flotillin-1 (Flot1) at the artificial cadAJ. Scale bar, 2 μm. Line profiles of E-cadherin, PS1 and Flot1 signals along a white dashed line. ΔI/I_0_ represents a fold change relative to nonjunctional membrane signals. **(G)** Confocal fluorescence images showing E-cadherin and Notch distributions at the artificial cadAJs after μMT application. Strong accumulation of Notch signals at the artificial cadAJ was clearly seen. Line profiles of MFN, E-cadherin, and Notch signals along a white dashed line. ΔI/I_0_ represents a fold change relative to fluorescence intensity before stimulation. Scale bar, 2 μm.

## CadAJs segregate Notch from γ-secretase, preventing interactions

To map the distribution of Notch signaling components, we generated a series of U2OS cells expressing recombinant Notch1, Dll1, and/or epithelial cadherin (E-cadherin, Ecad) proteins. To facilitate imaging, we fused them with self-labeling tags (SNAP or Halo-tags) and/or fluorescent proteins (EGFP or mCherry) at the N- or C-termini, respectively (**Table S1**). The endogenous Notch processing enzymes (*i.e*., ADAM 10/17, and γ-secretase) were immunostained and then imaged by confocal microscopy. With the exception of ADAM 10/17 which exhibited no preferential distribution relative to cadAJs (**fig. S1, A and F**), all other proteins exhibited cadAJ-dependent localization (**Fig. 1, B and C**). γ-secretase, visualized by staining with an anti-presenilin-1 antibody, was strongly enriched at the cadAJs with negligible non-junctional membrane signal (**Fig. 1, B and C, and fig. S1, B to E**) (*19*). In contrast, both Notch and Dll1 were excluded from cadAJs (**Fig. 1, B and C, and fig. S2**) and consequentially γ-secretase (**fig. S2, A and B**). Notch exclusion from cadAJs was observed in multiple contexts, including endogenous *vs*. recombinant expression of Notch (**fig. S3**), different cadherin types, cell types, and cell polarization states (**fig. S4**). Quantitative analysis using the Manders’ overlap coefficient (MOC) to calculate fractional overlap with E-cadherin also confirmed the enrichment of γ-secretase (0.85 ± 0.21) at cadAJs and the exclusion of Notch (0.24 ± 0.19) and Dll1 (0.26 ± 0.18) from cadAJs (**fig. S2, C and D**) (*19*). These observations suggest two mechanisms by which cadAJs might influence Notch signaling: first, cadAJs recruit γ-secretase; second, cadAJs segregate Notch ligands and receptors from γ-secretase to prevent their interactions.

To interrogate how cadAJs drive the spatial segregation of the enzyme (*i.e*. γ-secretase) and substrate (*i.e*. Notch) pair, we used our recently developed single-cell perturbation tool, mechanogenetics, which allows quantitative control over the location and mechanical loading of targeted receptors hence cell signaling (*9, 20, 21*). Specifically, we clustered E-cadherin labeled with magnetofluorescent nanoparticles (MFNs) to generate artificial cadAJs that recapitulate the functional and signaling roles of native cadAJs (*9, 20, 21*) (**Fig. 1, D and E**; See Methods for details). We then monitored the consequence of cadAJ formation on the spatial distribution of γ-secretase, Notch, and associated proteins (e.g., Flotillin-1, Flot1) relative to the artificial cadAJ (**Fig. 1, F and G**). To image full-length Notch explicitly, we prevented proteolytic processing by treating cells with TAPI2 and DAPT, ADAM 10/17 and γ-secretase inhibitors, respectively. Similar to native cell-cell cadAJs, γ-secretase was localized at the artificial cadAJ (**Fig. 1F and fig. S5**), indicating that cadherin clustering is sufficient to recruit γ-secretase to the cell surface. We also observed colocalization of Flot1, a protein enriched in spatially discrete and ordered membrane microdomains, at the artificial cadAJs (*16–18*) (**Fig. 1F and fig. S5**). This observation, along with analysis of native cell-cell junctions (**fig. S6**), molecular dynamics (MD) simulation (**fig. S7, A to C**), and cholesterol depletion experiments (**fig. S7D**), suggests that the cadAJs recruit and stabilize γ-secretase through a common spatially discrete and ordered membrane microdomain (for more discussion, see **Supplementary Text**).

In stark contrast to Notch depletion at native cadAJs (**Fig. 1, A** to **C**), we observed an intense Notch localization at the artificial cadAJ (**Fig. 1G and fig. S8**). The extraordinarily large size (extended length = 136 nm) of the Notch extracellular domain (NECD) (*22*) suggested a potential explanation for these contradicting observations. Specifically, Notch could be excluded from native cadAJs due to the NECD size greatly exceeding the narrow intermembrane cleft created by native cadAJs (20 nm) (*17, 23*). In contrast, artificial cadAJs generated by MFNs are free of membrane juxtaposition and lack a narrow intermembrane cleft, thus allowing for Notch diffusion and accumulation, presumably through its association with γ-secretase or other components of the membrane microdomains (*24*). These observations fit a model wherein the size-based physical constraint induced by cadAJ formation at a cell-cell interface segregates Notch from γ-secretase, preventing interactions that would otherwise serve to colocalize and concentrate the enzyme-substrate pair (*25*).

## Size-dependent segregation choreographs the Notch proteolytic sequence

Upon activation by juxtacrine ligand-receptor interactions, Notch is processed by ADAM 10/17 (S2 cleavage) and then by γ-secretase (S3 cleavage), sequentially (*1, 2*). Since each proteolytic event leads to a dramatic reduction in the size of NECD, we investigated how each cleavage step during activation correlates with the spatial distribution of Notch. We plated U2OS cells co-expressing SNAP-N^FL^-mCherry and Ecad-GFP on a Dll4-coated substrate to trigger Notch activation. Cells were also treated with TAPI2 (S2 cleavage inhibition) or DAPT (S3 cleavage inhibition) to capture Notch intermediates (**Fig. 2A**). To quantify the spatial redistribution of each Notch intermediate relative to the cadAJ, we measured the average mCherry fluorescence signal inside (I_IN_) and outside (I_OUT_) of the cadAJ and estimated an enrichment ratio (I_IN_/I_OUT_, See Materials and Method section) (**Fig. 2C**). An intensity ratio of 1 represents no cadAJ-dependent spatial localization as validated with Dil membrane dyes (I_IN_/I_OUT_ = 1.07, **Fig. 2C and fig. S9A**), and values less than or greater than 1 represent exclusion from or enrichment within cadAJs, respectively. With TAPI2, we observed minimal mCherry fluorescence signal from cadAJs (I_IN_/I_OUT_ = 0.50, **Fig. 2, B**(ii) **and C**(ii)), indicating that the ligand-receptor interaction did not alter the spatial distribution of Notch before S2 cleavage. In contrast, when TAPI2 was removed to activate S2 cleavage and DAPT was added to inhibit S3 cleavage, we observed strong enrichment of the mCherry signal at cadAJs (I_IN_/I_OUT_ = 2.14, **Fig. 2, B**(iii) **and C**(iii)). Not only did the Notch with extracellular domain truncation (NEXT, the product of S2 cleavage) relocalize, but it was in fact concentrated into cadAJs. When γ-secretase activity was rescued by washing out DAPT, the initially intense mCherry signal at cadAJs gradually disappeared (**Fig. 2, B**(iv) **and C**(iv)**, and fig. S9, B and C**), presumably corresponding to release of Notch intracellular domain (NICD) from the membrane. These results suggest a role for the size-dependent protein segregation as a spatial switch that regulates the distribution of Notch intermediates, thereby choreographing the sequential steps in Notch proteolysis. According to this model, the large size of NECD poses a physical constraint preventing entry of Notch to the narrow space between membranes in the cadAJ cleft where γ-secretase is localized. Hence, γ-secretase cannot process the full-length Notch before S2 cleavage. Following S2 cleavage, removal of NECD relieves the physical constraint, allowing Notch to enter into the cadAJ cleft. This facilitates a productive Notch-γ-secretase interaction, S3 cleavage, and then downstream signaling.

**Figure 2.**
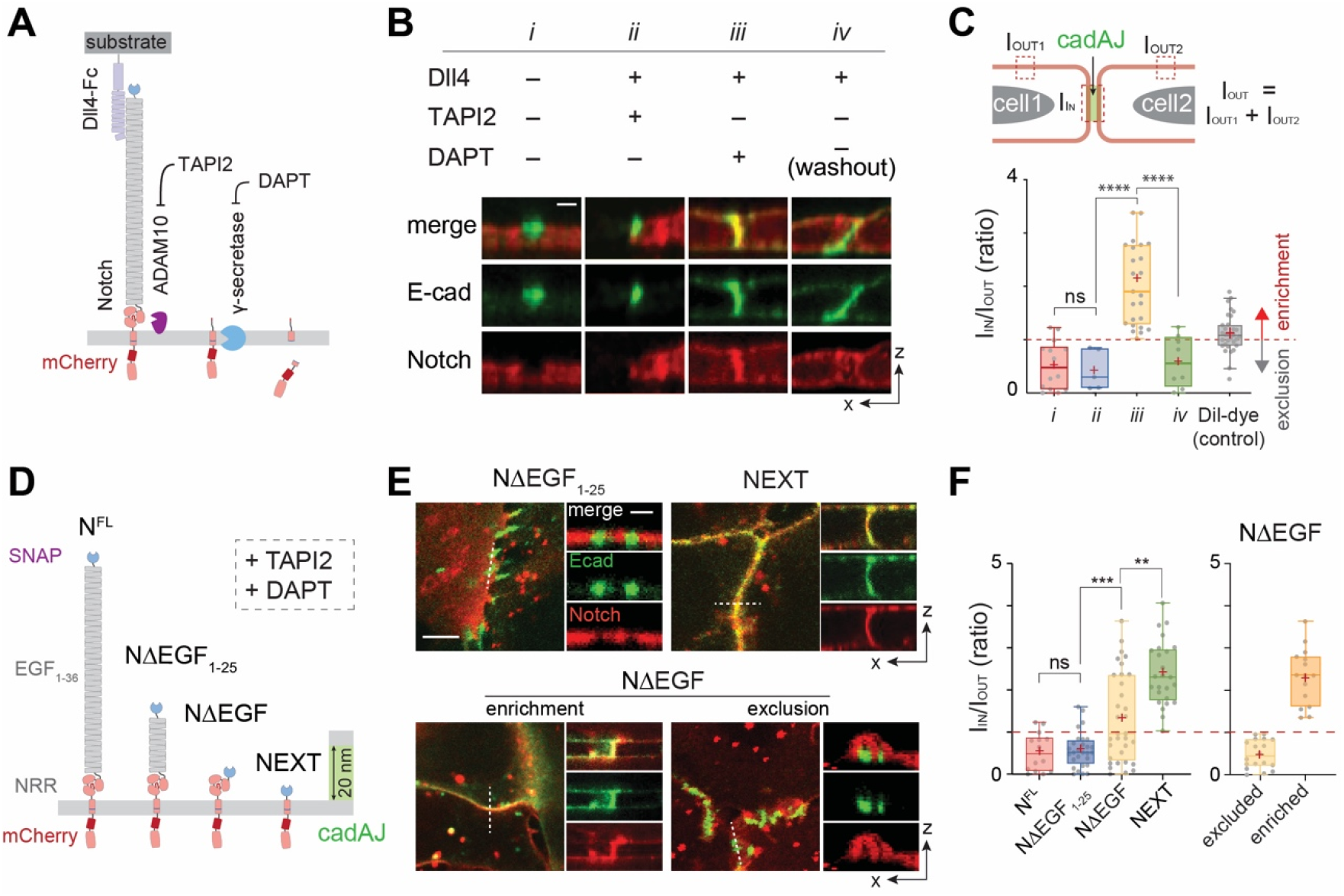
Size-dependent segregation of Notch at the cadAJs controls the proteolytic sequence. (**A**) A schematic to capture the spatial distribution of Notch intermediates during the cell-surface activation pathway. (**B**) Confocal z-resliced images showing Notch distribution (red) relative to cadAJ (green) from the cells without Dll4 activation (i), treated with Dll4 and TAPI2 (ii), treated with Dll4 and DAPT (iii), and washed out to remove DAPT inhibition (iv). Scale bar, 3 μm. (**C**) Quantification of Notch signal enrichment at the cadAJs. Notch enrichment (I_IN_/I_OUT_) is calculated as the ratio of average Notch fluorescence intensity within cadAJs (I_IN_) and outside cadAJ (I_OUT_). The enrichment factor of Dil is present as a control showing cadAJ-independent distribution. ****P < 0.0001, ns: non-significant, oneway ordinary ANOVA followed by Tukey’s multiple comparison testing. (**D**) Schematics of Notch variants with different truncation lengths, in comparison with the cadAJ intermembrane cleft. (**E**) Confocal fluorescence images showing spatial distribution of the Notch variants (red) relative to the cadAJs (green). To prevent any ligand-independent activation, cells were incubated with TAPI2 and DAPT. (left) Maximum projections of confocal z-stacks. Scale bar, 5 μm. (right) Confocal z resliced images along the white dashed lines in the maximum projection images. Scale bar, 2 μm. (**F**) Quantification of the enrichment factor (I_IN_/I_OUT_) of Notch variants relative to the cadAJs. A box-plot showing binary localization of NΔEGF which is defined as either excluded (yellow) or enriched (orange) is shown on the right. **P < 0.01, ***P < 0.001, ns: non-significant, one-way ordinary ANOVA followed by Tukey’s multiple comparison testing. Boxes and whiskers in (**C and F**) indicate the interquartile and the full ranges, respectively. Colored lines and (+) marks indicate median and mean, respectively.

To explore how size-dependent segregation controls Notch signaling, we first generated a series of U2OS cell lines stably expressing Notch variants with different truncation lengths: a partial deletion of the EGF repeats (NΔEGF_1-25_, approximate height: 48 nm), complete deletion of the EGF repeats but retention of negative regulatory region (NΔEGF, approximate height: 10 nm), and a complete removal of NECD (NEXT, approximate height: 4 nm) (**Fig. 2D**). We fused these Notch variants with SNAP- and mCherry-tags at their N- and C-termini, to differentially image the extracellular and intracellular domains. All cells were treated with TAPI2 and DAPT to prevent any potential proteolysis of the variants. Consistent with predictions based on size-dependent protein segregation, NΔEGF_1-25_, the Notch variant with an ECD taller than the height of the intermembrane cadAJ cleft, was excluded from cadAJs (I_IN_/I_OUT_ = 0.57) (**Fig. 2, E and F**). NEXT with an ECD smaller than the junctional height was enriched at cadAJs (I_IN_/I_OUT_ = 2.39) (**Fig. 2, E and F**). Interestingly, we observed a mixed binary localization pattern of NΔEGF (intermediate height) relative to cadAJs, with a mean I_IN_/I_OUT_ value of 1.32 (**Fig. 2, E and F**). Some cadAJs displayed NΔEGF enrichment (**Fig. 2, E**(bottom left) **and F**(right)), consistent with the size-based prediction. Meanwhile, other cadAJs excluded NΔEGF (**Fig. 2, E**(bottom right) **and F**(right)). Considering the fact that density of cadherin clusters within cadAJs vary with the size, type, and degree of junction maturation (*26, 27*) and that the glycosylated negative regulatory region domain (*6*) is susceptible to steric crowding, this unanticipated exclusion might result from a lateral crowding effect in high-density cadAJs.

We next investigated the functional consequences of the two populations of cadAJs at the cell surface – the pool that excludes partially truncated NΔEGF and the pool that enriches NΔEGF in the presence of TAPI2 and DAPT. Focusing on the subset of cadAJs showing strong enrichment for Notch signal, we inferred γ-secretase activity by measuring the changes in extracellular SNAP (labeled with SNAP surface dye) and intracellular mCherry fluorescence signals after DAPT removal but in the presence of TAPI2. This construct has different fluorescence markers on the extracellular and intracellular domains of the protein, allowing us to differentially map the S2 and S3 cleavage events of the construct. While SNAP fluorescence signal remained strong, the mCherry signal at the cadAJ was negligible, indicating selective release of NICD from cadAJs (**Fig. 3, A and B**). Consistently, we also observed a correspondingly modest but statistically significant increase of the nuclear mCherry signal, suggesting nuclear translocation of cleaved NICD (**fig. S10**).

**Figure 3.**
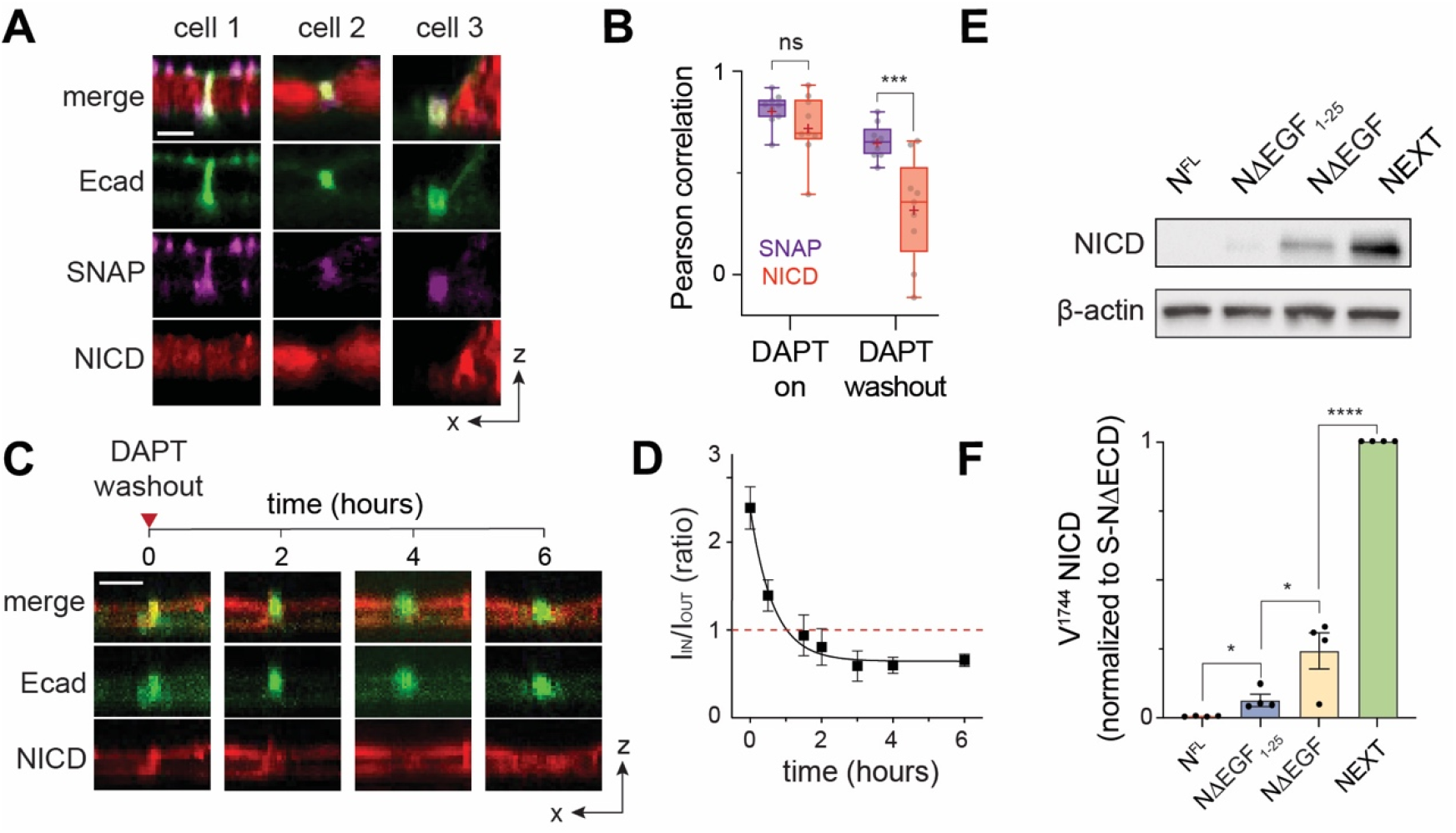
Colocalization of Notch with γ-secretase is sufficient to trigger its proteolysis and signaling, regardless of S2 cleavage or ligand presentation. (**A**) Confocal *z*-resliced images showing the distribution of extracellular SNAP (purple) and intracellular mCherry (red) tags of NΔEGF relative to cadAJs (green) after DAPT removal. Scale bar, 3 μm. (**B**) A box-whisker plot showing Pearson correlation coefficients of extracellular-SNAP (purple) and intracellular-mCherry (red) domains relative to the cadAJs before and after DAPT washout (***P < 0.001; ns, not significant; n = 9 biological replicates; two-tailed unpaired Student’s t-test). Boxes and whiskers indicate the interquartile and the full ranges, respectively. Colored lines and (+) marks indicate median and mean, respectively. (**C**) Time series of confocal z-resliced images showing the enrichment of NΔEGF (red) at the cadAJ (green) under DAPT treatment (t = 0), and the dissipation during DAPT washout (t ≥ 2). Scale bar, 3 μm. (**D**) Single-cell traces showing the time-course of the decline of NΔEGF enrichment factor at the cadAJs during DAPT washout (mean ± s.e.m.; n = 4 independent single-cell experiments). (**E and F**) Western blot analyses of cleaved NICD levels in the cells stably expressing N^FL^, NΔEGF_1-25_, NΔEGF, and NEXT. All cells were transfected with Ecad-GFP and incubated with TAPI2 for 24 hr. β-actin levels represent the loading control. A representative image of immunoblotting (**E**), and quantification (**F**) of cleaved NICD levels. The average intensity of each NICD band relative to respective β-actin band was quantified and then normalized to that of NEXT (mean ± s.d.; n = 4 biological replicates; *P < 0.05, ****P < 0.0001, one-way ANOVA followed by Tukey’s multiple comparison test).

We measured the kinetics of S3 processing by tracking mCherry intensity at the cadAJ in single cells. Initially intense mCherry signal (average I_IN_/I_OUT_ = 2.89 ± 1.15) rapidly dissipated within the first 2 hours following DAPT removal (average I_IN_/I_OUT_ = 0.81 ± 0.42), reaching a plateau at 4 hours (I_IN_/I_OUT_ = 0.65 ± 0.14) **(Fig. 3, C and D, and fig. S11, A and B**). Removing DAPT did not elicit a significant reduction in mCherry signal intensity from the non-cadAJ membrane, indicating that the S3 cleavage activity was strongly localized at the cadAJ (**fig. S11C**).

To confirm that the observed decrease in mCherry signal at the cadAJ corresponds specifically to the S3 cleavage, we performed western blot analysis of the cells expressing the Notch variants. We cultured cells with TAPI2 to decouple γ-secretase processing from S2 cleavage, and measured cleaved NICD levels by immunoblotting with Notch antibodies that detect N-terminal V1744. Cells expressing N^FL^ or NΔEGF_1-25_ resulted in no or minimal NICD, respectively (**Fig. 3, E and F** and **fig. S13A**). Whereas, cells expressing NΔEGF produced a significant amount of NICD (**Fig. 3, E and F** and **fig. S13A**). Cells expressing NEXT exhibited the highest NICD production, about a four-fold increase compared with that of NΔEGF (**Fig. 3, E and F** and **fig. S13A**). The observed NICD production was proportional to the enrichment ratio (I_IN_/I_OUT_) of the Notch variants at cadAJs, suggesting the essential role of size-dependent protein segregation as a spatial switch to direct Notch activation. The substantial NICD production from the cells expressing NΔEGF indicates that, when localized together, γ-secretase can process Notch, bypassing S2 cleavage. Size-dependent but ligand-independent activation of Notch receptors with an intact S2 site was observed previously in Notch variants and synNotch constructs (*28–32*), but the mechanism of this activation has been unclear so far. Our observations support the notion that colocalization of these Notch variants with γ-secretase is sufficient to trigger S3 proteolysis and signaling.

## Spatial mutations alter Notch signaling

Prevailing models of Notch proteolysis by γ-secretase are based on the notion that S2-cleavage of Notch serves to potentiate the cleavage by modifying the molecular interface at the enzyme-substrate pair (*31, 33*). For example, it has been suggested that γ-secretase selectively recognizes S2-cleaved Notch (*i.e*., NEXT) through hydrogen bonding between a glutamate residue in nicastrin and the new N-terminus of NEXT (*33*). Another model proposed that S2 cleavage serves to reduce steric repulsion between nicastrin and NECD, strengthening their interaction (*31*). However, another key feature of S2 cleavage is that it generates a smaller molecular intermediate that can uniquely access cadAJs, thereby colocalizing Notch with γ-secretase and significantly increasing its concentration near the enzyme active site. This suggests a third model, wherein γ-secretase activity on full length Notch and its intermediates is blocked by maintaining concentrations of Notch below the K_M_ for γ-secretase through spatial segregation.

To explicitly test the effect of Notch spatial localization relative to cadAJs and γ-secretase on the signaling, we designed three experiments that induce spatial mutations of Notch. First, we employed a DNA-mediated crosslinking strategy to enhance NΔEGF – a Notch variant that exhibited a binary localization relative to cadAJs and relatively low Notch activation – enrichment at the cadAJ. We generated cells co-expressing SNAP-NΔEGF-mCherry and Halo-Ecad-GFP and treated the cells with complementary benzylguanine (BG)-and chloroalkane-modified oligonucleotides in the presence of TAPI2 and DAPT (**Fig. 4A**). Notch-E-cadherin heterodimers were formed efficiently as evidenced by the appearance of a higher molecular weight band corresponding to the DNA-linked complex on western blots (**fig. S13, B and C**). Compared to untreated cells (I_IN_/I_OUT_ = 1.32 ± 1.06), we observed further enrichment of NΔEGF at cadAJs in the presence of DNA crosslinking (I_IN_/I_OUT_ = 1.89 ± 0.91) (**Fig. 4B**). We then maintained cells in TAPI2 but removed DAPT to allow S3 cleavage. We observed decreases in mCherry signal at cadAJs after DAPT removal, indicating efficient S3 cleavage without S2 cleavage (**fig. S13, D to F**). Accordingly, in western blots, we observed increased V1744-terminated NICD levels from the cells treated with DNA crosslinkers, compared with the untreated control (**Fig. 4C**). There results suggest that new molecular interfaces produced by mechanical activation and S2 cleavage are not necessary when γ-secretase is concentrated with its substrate. Considering that DNA crosslinking (molecular weight = 21.4 kD) increases the ECD size of NΔEGF, the observed increase in NICD production cannot be explained by the nicastrin-induced steric repulsion model. Rather, this result favors a model wherein the increased concentration of the NΔEGF at cadAJs facilitated its interaction with γ-secretase and thus promoted S3 cleavage.

**Figure 4.**
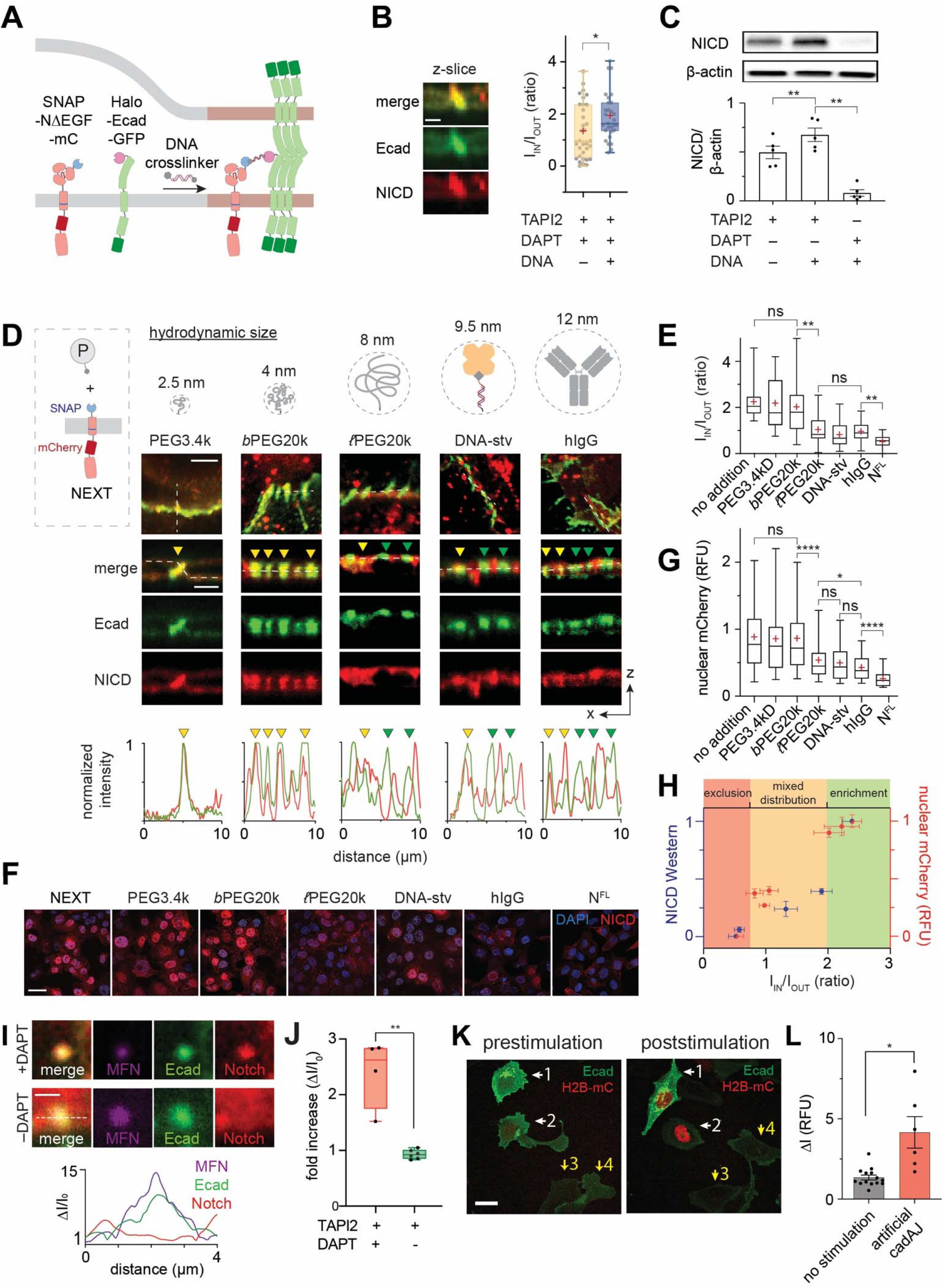
Spatial mutations alter Notch activation, regardless of ligand presentation or S2 cleavage. **(A)** A schematic describing DNA-mediated crosslinking strategy to enhance NΔEGF localization at the cadAJ. (**B**) Confocal z-resliced images showing intense NΔEGF fluorescence (red) enriched at the cadAJ (green) after the DNA crosslinking. Scale bar, 3 μm. Quantification of enrichment (I_IN_/I_OUT_) without (n = 33) and with (n = 29) DNA crosslinker treatment, indicating an increase in enrichment after the crosslinking (*P < 0.05; two-tailed Mann-Whitney-Wilcoxon test). (**C**) Western blot analyses showing increased S3-cleaved NICD levels in the NΔEGF cells treated with the DNA crosslinker. (top) A representative gel image showing immunoblotting for NICD and β-actin. (bottom) Quantification of cleaved NICD levels. The average intensity of NICD band was normalized to that of β-actin band in each sample. (mean ± s.d.; **P < 0.01; n = 5 biological replicates; ordinary one-way ANOVA). (**D**) Spatial mutation of NEXT via chemical ligation of macromolecular pendants (denoted as P). BG-modified polymers or proteins were conjugated to the extracellular SNAP tag of NEXT. Cartoons depicting shape and hydrodynamic size of different pendants are shown. Confocal images showing size-dependent spatial mutation of NEXT (red) at the cadAJs (green). The top row shows maximum projection images of the cells treated with the indicated pendants. Scale bar, 5 μm. The middle row shows confocal z resliced images along the white dashed lines in the maximum projection images. Yellow and green arrowheads indicate the cadAJs enriches with and those that excludes Notch, respectively. Scale bar, 3 μm. The bottom row shows line profiles quantifying fluorescence signals from NEXT (red) and E-cadherin (green) along the white lines in the z-resliced images. Images and line profiles are representative of n ≥ 15 biological replicates. (**E**) Quantification in I_IN_/I_OUT_ of NEXT with macromolecular pendants (n ≥ 15). (**F and G**) Confocal fluorescence images (**F**) and quantification (n ≥ 180) (**G**) of nuclear mCherry signals of the NEXT-expressing cells treated with macromolecular pendants. DAPI signals (blue) indicate cell nucleus. Scale bar, 5 μm. Cells expressing N^FL^ were used as a negative control. **(H)** A plot representing the NICD level of various Notch variants as a function of the enrichment factor (I_IN_/I_OUT_, mean ± s.e.m.; n ≥ 15 biological replicates). All Notch variants with different truncation length, DNA crosslinking, or pendant addition used in Figures 2 and 3 were included. median ± s.e.m.; n ≥ 4 for western blot for NICD levels; n ≥ 180 for nuclear mCherry fluorescence. **(I)** Representative confocal fluorescence images of cells with an artificial cadAJ in the presence of TAPI2 and DAPT (upper), and TAPI2 only (lower). Line profiles of MFNs, E-cadherin, and Notch signal along the white dashed line. ΔI/I_0_ represents a fold change relative to nonjunctional membrane signal. Scale bar, 2 μm. **(J)** Quantification of mCherry signal at artificial cadAJs after μMT application in the presence or absence of DAPT (with DAPT, n = 4; no DAPT, n = 6). (**E, G, J**) Boxes and whiskers indicate the interquartile and the full ranges, respectively. Black lines and (+) marks indicate median and mean, respectively. **P < 0.01; ****P < 0.001; ns, not significant; ordinary one-way ANOVA followed by Tukey’s multiple comparison. **(K)** Representative confocal fluorescence images of the reporter cells with artificial cadAJs. White arrows: cells with stimulation, Yellow arrows: control cells. Scale bar, 10 μm. **(L)** Statistical analysis of stimulated cells (n = 6) vs. control cells (n = 14). Error bars indicate SEM. *P < 0.05; ns, not significant; ordinary one-way ANOVA.

To further test the importance of size-dependent spatial segregation, we induced spatial mutation of NEXT (*i.e*., the Notch variant that showed the strongest enrichment at the cadAJ and activation) by chemically conjugating it with macromolecules. Specifically, we conjugated BG-modified polymers and proteins, to the extracellular SNAP tag (4.0 nm) of the variant via BG-SNAP chemistry (**Fig. 4D**). Grafting of these macromolecular pendants onto NEXT increases the size of the Notch construct but does not modify the N-terminal amine for hydrogen bonding with nicastrin. To interrogate the size-dependent spatial mutation of NEXT systematically, we used a series of pendants with different hydrodynamic sizes, that includes polyethylene glycol with an average molecular weight of 3.4 kD (PEG3.4k, 2.5 nm), branched PEG20k (*b*PEG20k, 4.0 nm), linear PEG20k (*ℓ*PEG20k, 8.0 nm), DNA-streptavidin conjugates (DNA-stv, 9.5 nm), and human immunoglobulin G (hIgG, 12 nm) (**Fig. 4D and fig. S14A**). In the presence of DAPT, we observed a size-dependent distribution of NEXT at the cadAJ, where the larger pendant resulted in a greater decrease in mCherry signal at the cadAJ. With pendants smaller than 5 nm (*i.e*., PEG3.4k and *b*PEG20*k*), NEXT remained enriched at the cadAJ with I_IN_/I_OUT_ of 2.21 and 2.01, respectively (**Fig. 4, D and E**). When *ℓ*PEG20k, DNA-stv, or hIgG were added, we observed a binary localization pattern of NEXT (*i.e*., enriched at or excluded from the cadAJs) with mean I_IN_/I_OUT_ values of 1.06, 0.82, or 0.98, respectively (**Fig. 4, D and E**). These observations were similar to the mixed spatial behavior of NΔEGF having a comparable ECD size, where only less dense cadAJs allowed Notch colocalization. We then examined the signaling consequences of each size-dependent spatial mutation of NEXT. Following S3 cleavage, NICD traffics to the nucleus, allowing us to measure nuclear mCherry signal as a proxy for Notch pathway activation. The PEG3.4k or *b*PEG20k addition did not significantly alter nuclear mCherry signal of NEXT, compared with cells with no pendant addition (**Fig. 4, F and G, and fig. S14B**). Conjugation of *ℓ*PEG20k and DNA-stv resulted in a substantial decrease in nuclear mCherry signal to 0.39 and 0.37, respectively (**Fig. 4, F and G, and fig. S14B**). hIgG addition suppressed nuclear mCherry signal further to 0.27 (**Fig. 4, F and G, and fig. S14B**). We summarized the NICD production of all Notch variants as a function of the Notch enrichment factor, I_IN_/I_OUT_, in **Fig. 4H**, clearly visualizing the spatial dependence of S3 cleavage.

Lastly, we investigated whether γ-secretase can process full-length Notch when specifically directed to cadAJs by artificial means. To do so, we generated artificial cadAJs on the cells expressing SNAP-N^FL^-mCherry using mechanogenetics, in the presence of TAPI2 (to prevent S2 cleavage) but without DAPT (to allow γ-secretase activity). Contrary to the experiment with DAPT (**Fig. 1G and fig. S8E)**, we observed no enrichment of mCherry signal at the artificial cadAJ, presumably due to NICD release (**Fig. 4, I and J, fig. S8F)**. To confirm that the loss of mCherry signal corresponded to bona fide signaling from Notch, we employed a UAS-Gal4 reporter system that detects Notch activation with the nuclear mCherry fluorescence (*8, 9, 34*). To a cell recombinantly expressing SNAP-Notch-Gal4 (SNAP-N^FL^-Gal4) and Halo-Ecad-GFP, we again generated artificial cadAJs via mechanogenetics and measured the nuclear mCherry fluorescence every two hours. Note that no source of S2 cleavage (*e.g*.,no ligand-immobilized substrate) was added. We observed strong nuclear mCherry signal from the cells with artificial cadAJs, but no signal from neighboring cells (**Fig. 4, K and L, fig. S12**). Altogether, these results suggest that the size-dependent spatial switch serves as a substrate-selection mechanism for γ-secretase.

## The cadAJ-mediated spatial switch is indispensable for Notch signaling

Given the significant role of cadAJs in coordinating the spatial dynamics of Notch and γ-secretase, we next interrogated Notch signal activation in cells lacking cadAJs. To minimize physical contact between cells and hence cadAJ formation, we sparsely plated cells expressing SNAP-N^FL^-Gal4 on a Dll4-coated substrate – effectively decoupling cell-cell contact from Notch-Dll4 interactions by allowing ligand presentation from the glass substrate rather than neighboring cells. After 16 hours from cell seeding, we measured the mCherry fluorescence in cells having no prior contact with other cells. For comparison, we also analyzed the mCherry signal in cells with robust cadAJs within high-density culture. While cells with physical contacts with adjacent cells exhibited a robust increase in nuclear mCherry fluorescence signal, those without cell-cell contact elicited no increase in signal (**Fig. 5, A and B, fig. S15A, and Movie S1**). Reestablishing cadAJs by plating cells on a substrate coated with Ecad-Fc and Dll4-Fc rescued the Notch signaling of the solitary cells (**Fig. 5, A and C, and Movie S2**). These results support that cadAJs (or some other means of enriching γ-secretase to the cell surface) are required for Notch processing at the cell surface and downstream signaling. Critically, E-cadherin seems to function in this capacity in a manner that is independent of its role in mediating cell-cell contact. To further test this notion, we knocked out the gene encoding E-cadherin (CDH1) in the reporter cell line via CRISPR-Cas9 (**fig. S16**), then plated the cells at high density on Dll4-Fc coated plates. Strikingly, E-cadherin knockout (Ecad-KO) resulted in abrogation of Notch activation even with robust cell-cell contact (**Fig. 5, D and E, and fig. S15B**). Reintroduction of plasmids encoding E-cadherin or N-cadherin into Ecad-KO cells recovered Notch activation with substantial nuclear mCherry signal (**Fig. 5, D and E, and fig. S15B, B to D**). Single cell analysis of the nuclear fluorescence signal exhibited a clear positive correlation with E-cadherin expression in the respective cells, confirming cadAJ-dependent Notch signaling (**fig. S15E)**.

**Figure 5.**
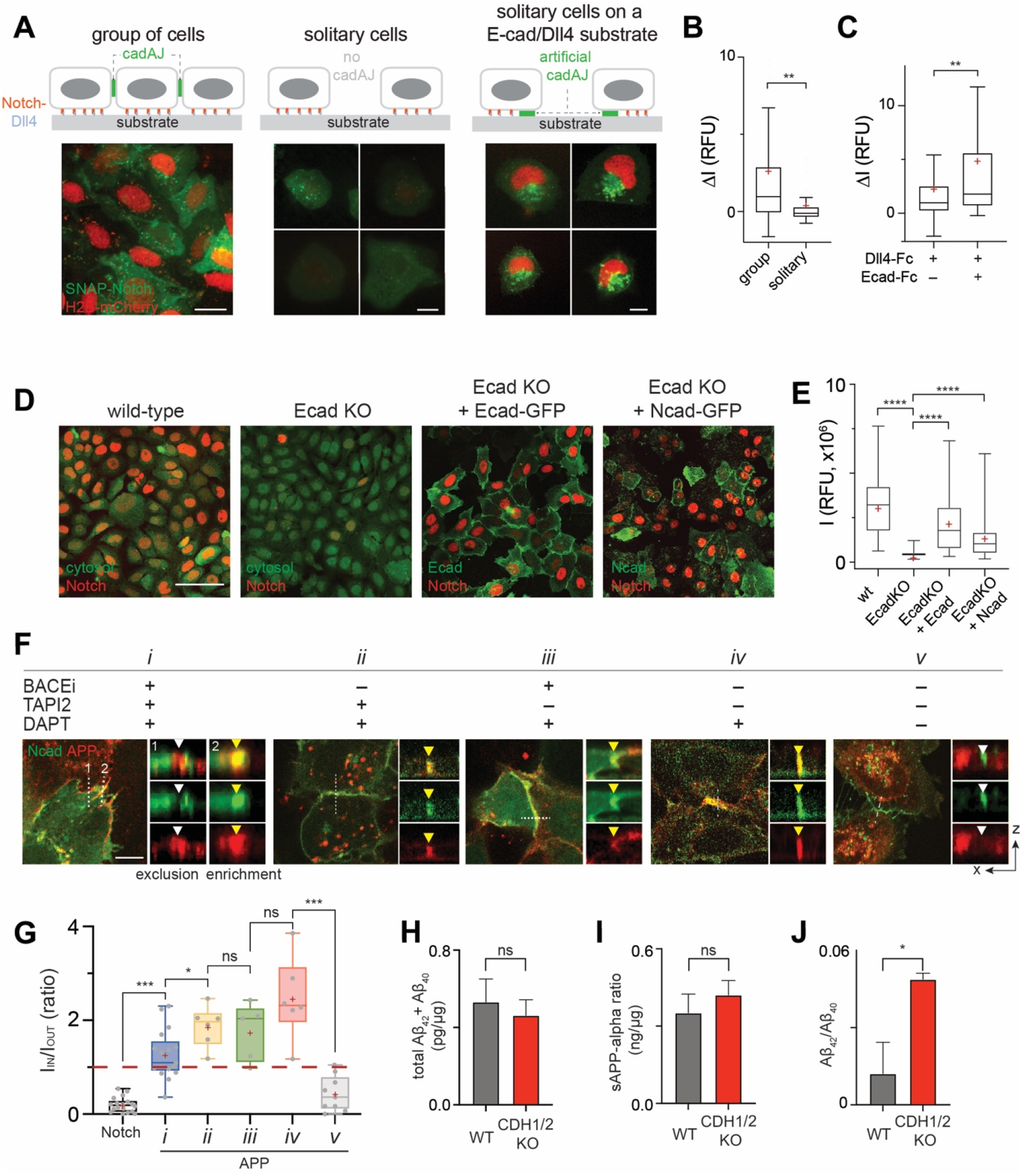
The cadAJ-mediated spatial switch regulates Notch and APP signaling. (**A**) Representative epi-fluorescence images showing Notch activation in U2OS SNAP-N^FL^-Gal4 reporter cell lines in different cellular environments: Group of cells on a Dll4-Fc coated substrate (left), solitary cells with no prior contact on a Dll4-Fc coated substrate (middle), and solitary cells plated on a Dll4-Fc and Ecad-Fc coated substrate (right). Scale bars, 20 μm (see Movie S1 and S2). (**B**) Quantification of Notch activation by measuring H2B-mCherry fluorescence changes in cells within a group (n = 152), solitary cells (n = 50). (**C**) Quantification of Notch activation in solitary cells cultured on a Dll4-Fc coated substrate (n = 27) and those cultured on a Dll4-Fc and Ecad-Fc coated substrate (n = 27). (**D**) Representative confocal images of H2B-mCherry fluorescence in U2OS SNAP-N^FL^-Gal4 reporter cells (wt), E-cadherin knockout cells (Ecad-KO), Ecad-KO cells with recombinant E-cadherin transfection (Ecad-KO + Ecad), and Ecad-KO cells with N-cadherin transfection (Ecad-KO + Ncad). Cytosol labeled with CMFDA dye was shown for wt and Ecad-KO cells. E-cadherin and N-cadherin were shown for Ecad-KO + Ecad and Ecad-KO + Ncad cells. Scale bar, 100 μm. (**E**) Quantification of Notch activation in the wt (n = 86), Ecad-KO (n = 100), Ecad-KO + Ecad (n = 52), and Ecad-KO + Ncad (n = 80) cells. (**F**) Confocal fluorescence maximum projection (left) and z-resliced images (right) of U2OS cells co-expressing Ncad-mCherry (green) and APP-EGFP (red) in different combinations of α-, β-, and γ-secretase inhibitors. Scale bars, 10 μm (maximum projection) and 3 μm (inset). (**G**) Quantification of the enrichment factor (I_IN_/I_OUT_) of APP signal relative to the NcadAJs. **(H-J)** Total sum of Aβ_42_ and Aβ_40_ **(H)**, soluble APPα **(I)**, and Aβ_42_/Aβ_40_ ratio (**J**) measured by ELISA in wild-type cells or CDH1/2 KO cells (mean ± SEM, n = 3, *P< 0.05, two-tailed paired Student’s t test). (**B**, **C**, **E, and G**) Boxes and whiskers indicate the interquartile and full ranges, respectively. Black lines and (+) marks indicate median and mean, respectively. *P<0.05; **P < 0.01; ****P < 0.001; ns, not significant; unpaired two-tailed t test in **(B)** and **(C)**; ordinary one-way ANOVA followed by Tukey’s multiple comparison in (**E**) and **(G)**.

## Size-dependent spatial dynamics and proteolysis of amyloid precursor proteins

The proteolysis processes of amyloid precursor protein (APP) plays a central role in amyloid beta (Aβ) pathology, which can cause failures in many organs such as brain, heart, kidney, and vasculature (*35–38*). Interestingly, APP has a strikingly similar topology and proteolytic cleavage sequence to that of Notch. APP possesses a large ECD (> 20 nm) and is processed by α-or β-secretase and then γ-secretase (*35–37*). These characteristics motivated us to investigate the generality of size-dependent protein segregation to APP proteolysis by γ-secretase. We generated U2OS cells co-expressing APP-GFP and SNAP-N-cadherin (SNAP-Ncad) and monitored the cell surface spatial dynamics of APP intermediates relative to N-cadherin-based AJs (NcadAJs) in the presence of protease inhibitors. Having an intermediate ECD height (80 kD in size), full-length APP showed binary localization (*i.e*.,excluded or enriched) relative to cadAJs in the presence of inhibitors (**Fig. 5, F and G)**, similar to the Notch variant with EGF repeat truncation (*i.e*., NΔEGF). APP diffused into the NcadAJs after ECD shedding by α- or β-secretase, and then was processed by γ-secretase within it (**Fig. 5, F and G, fig. S18**).

APP proteolysis by γ-secretase produces more soluble p3 and Aβ_40_ predominately, along with less soluble and pathogenic Aβ_42_ and longer isoforms (*35–37*). It has been previously shown that local acidic pH environment (e.g., pH 5.5) leads to a gain in the proportion of pathogenic Aβ species (*39*). Additionally, N-cadherin expression in cells stabilizes an open conformation of PS1 that favors Aβ_40_ production over Aβ_42_ (*40*). Given our previous observation that loss of cadAJs leads to a decrease in cell surface γ-secretase, we hypothesized that APP processing would be biased under these conditions towards Aβ_42_. We tested this hypothesis by constructing U2OS cell lines lacking both E- and N-cadherins (CDH1/2-KO cells) using CRISPR-Cas9 (**fig. S17**). We then transfected plain U2OS cells or CDH1/2-KO cells with a plasmid encoding APP sequence and measured APP fragment production by ELISA. While no significant changes were observed in total Aβ(40+42) and soluble APPα(sAPPα) (**Fig. 5, H and I**), CDH1/2-KO cells produced higher relative levels of Aβ_42_, the isoform prone to severe fibril aggregation, compared to cells with endogenous cadherin expression (**Fig. 5J**).

## The cadAJ-mediated spatial switch regulates neuronal progenitor cell differentiation *in vivo*

Notch signaling is essential for the maintenance of stemness, self-renewal, and differentiation of neural progenitor cells (NPCs) (*41*). In the mammalian cerebral cortex, Notch signaling orchestrates developmental neurogenesis, where it modulates a balance between tangential proliferative (*i.e*., symmetric division) and radial differentiative (*i.e*., asymmetric division) expansion of the apical ventricular-zone NPCs (VZ-NPCs) to establish a stratified neuronal organization (*42*). Several reports also emphasize the critical role of apical-endfoot cadAJs in Notch signaling and the decision-making process of VZ-NPC development (i.e., proliferation vs. differentiation) (*43–45*).

Given the essential role of the cadAJ-mediated spatial switch in Notch signaling of cell line models, we mapped the spatial distribution of Notch and γ-secretase relative to cadAJs in VZ-NPCs of the developing mouse brain (E13.5) (**Fig. 6**, **A** to **G**). Consistent with observations in cell lines, Notch and PS1 exhibited exclusion (MOC = 0.14 ± 0.05, n = 9) from and enrichment (MOC = 0.69 ± 0.07, n =9) within NcadAJs, respectively (**Fig. 6, C** to **G**). We also captured the spatial distribution of the Notch activation intermediate by intracerebroventricular injection of DAPT into postnatal mice (P3). The immunostaining showed inclusion of Notch signal within NcadAJs, presumably resulting from NEXT accumulation (**Fig. S19A**) as observed in cell lines (**Fig. 2B**). These results support the notion that cadAJs also serve as a spatial switch regulating Notch signaling *in vivo*.

**Figure 6.**
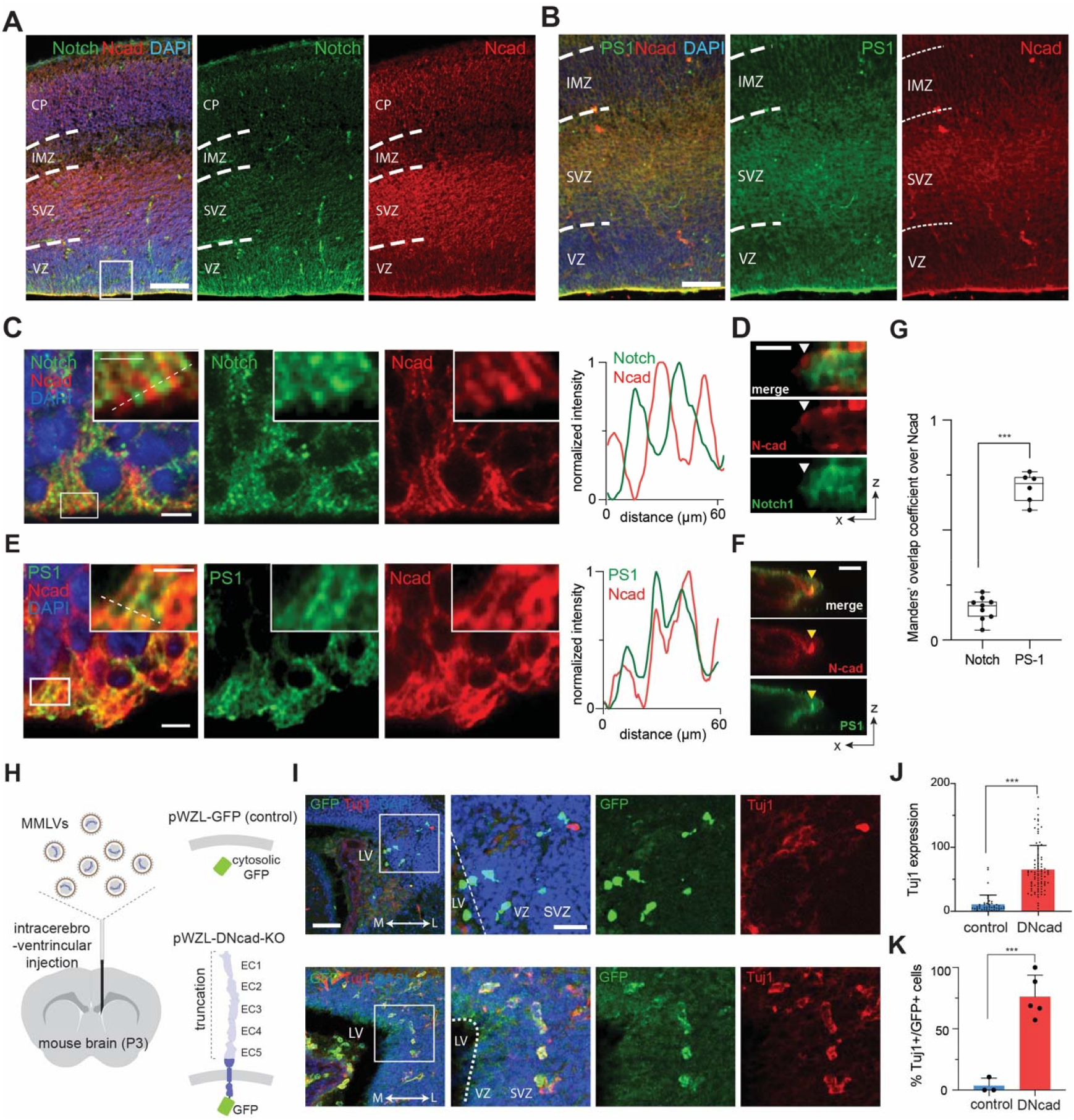
The cadAJ-mediated spatial switch regulates neuronal progenitor cell differentiation *in vivo*. (**A to F**) Immunostaining of the subventricular zone (SVZ) in the lateral ventricle (LV) of the E13.5 mouse brain. Notch (**A, C, D**) and PS1 **(B, E, F**) distributions relative to NcadAJs. **(A, B)** Representative low (**A and B**) and high (**C and E**) magnification images. Scale bars, 100 μm and 5 μm, respectively. The boxed area in panels (C) and (E) is further magnified in the inset. Scale bar, 2.5 μm. Line profile analysis shown in panels (C) and (E). **(D and F)** Representative confocal z-resliced image showing Notch exclusion (white arrowhead) and PS1 colocalization (yellow arrowhead) with the cadAJ. Scale bar, 3 μm. (**G**) Quantitative assessment of Notch and PS1 colocalization with N-cadherin *in vivo*. Each dot represents MOCs quantifying colocalized Notch (n = 9) or PS1 (n = 6) over selected cadAJs. **(H)** Retroviral infection of a plasmid encoding control vector (EGFP) or dominant negative form of E-cadherin vector (DN-cad-EGFP) to developing P3 mice via intracerebroventricular injection. **(I)** Retroviral infection of DN-cad-GFP increases the differentiation of NPCs compared with control. Cells differentiated into post-mitotic neurons can be identified as EGFP+/Tuj1+, while those which remained as NPCs with plasmid transfection are only EGFP+. Scale bar, 50 μm. **(J)** Quantification of the expression of Tuj1 per single cells (n = 43 cells across 3 mice and n = 86 cells across 5 mice per control and DN-cad, respectively). **(K)** Quantification of the percentage of Tuj1-expressing post-mitotic neurons among all transfected EGFP+ cells were quantified. Data are represented as mean ± SEM. ** p<0.01, *** p<0.001, two-tailed unpaired Student’s t-test.

To understand the function of the spatial switch on VZ-NPC development, we disrupted cadAJs via dominant-negative cadherin expression. We retrovirally transfected a plasmid encoding a dominant-negative form of E-cadherin with the extracellular domain truncation (DN-cad) (*45*) and a C-terminal GFP tag to VZ-NPCs of developing mice (P3) via intracerebroventricular injection (**Fig. 6H**). 48 hours after transfection, we analyzed NPC differentiation via TuJ1 immunostaining, a neuronal marker. While mice transfected with a control plasmid (n = 3) showed negligible TuJ1 signals, those with DN-cad plasmid transfection (n = 5) exhibited robust TuJ1 expression, presumably through downregulation of Notch signaling (**Fig 6, I** to **K** and **fig. S19, B and C**). These results support that cadAJ-mediated spatial switch modulates NPC maintenance and differentiation via Notch signaling.

## Discussion

Unlike most other juxtacrine signaling systems, the Notch ligand-receptor interaction (chemical switch) is converted into intracellular signals only following multiple additional regulatory steps gated by mechanical, enzymatic, and spatial events. These include unfolding of the negative regulatory region domain (mechanical switch), S2- and S3-cleavage (enzymatic switch), and finally translocation of the NICD from the cell membrane to the nucleus (spatial switch) (*1, 2*). Our study reveals that Notch integrates an additional spatial switch at the cell surface to tightly choreograph the enzymatic cleavage sequence prior to NICD release. Previously, it was thought that this enzymatic sequence was regulated by modification of the molecular interface between Notch and nicastrin after S2-cleavage (*33, 46*). Our model is not incompatible with a contribution of the nicastrin-Notch chemical interface on γ-secretase activity. However, it strongly suggests that the spatial switch is the major regulator of Notch-γ-secretase interaction and signaling, functioning by increasing the concentration of the γ-secretase substrate to the point that it exceeds the K_m_ and is efficiently processed by the enzyme. Particularly, our mechanogenetic experiment shown in **Fig. 1, D** to **G** and **Fig. 4, I** to **L**, respectively, supports the notion that Notch with an intact S2 site and when concentrated together with γ-secretase, is effectively engaged and then processed so long as the spatial constraints of juxtaposed cell membranes at cadAJs are removed.

The operating principle of the spatial switch is closely related to another unique feature of Notch receptor: its unusually tall NECD. The functional residues responsible for ligand binding are located near the N-terminus, which protrudes above the crowded cell surface, where they are poised to engage ligands on neighboring cells. Surprisingly, however, it has been also shown that replacing the EGF-like domain repeats with a relatively smaller ligand binding domain (*e.g*., synNotch) maintains the receptor function (*47*). Why then does Notch receptor bear such a massive ECD? Our study provides insight into this question, where the large ECD is crucial for its spatial segregation from γ-secretase thereby minimizing nonspecific ligand-independent activation. Low level NICD production was observed even for Notch variants with partial EGF truncation (NΔEGF_1-25_) and levels gradually increased upon successive truncations to the NECD size. NΔEGF_1-25_ has comparable size with smaller Notch family proteins, including C. elegans LIN-12/Notch and GLP-1/Notch (13 and 10 EGF repeats, respectively), suggesting the relevance of spatial switch across the Notch family and metazoans. Our model also explains previous observations where synNotch with a relatively small ECD exhibited significant ligand-independent activation (10-50% of ligand-induced activation) (*29*).

We also show that the size-dependent spatial segregation regulates APP cleavage and Aβ production. It has been previously shown that γ-secretase presenting in different subcellular compartments cleaves APP into diverse Aβ isoforms (*36, 37, 48*). Our study shows that, after the ECD cleavage, cadAJ potentiates cell surface processing of APPs within the junction, yielding Aβ_40_ predominantly, while removal of cadAJ produces more Aβ_42_. To establish the relevance of this observation to APP processing will require further investigation in a neuronal system, but our results in model cell lines are consistent with the predominant secretion of Aβ from the synapse, where N-cadherin junctions localize (*49*).

Our study also suggests a critical role of the cadAJ-mediated spatial switch in VZ-NPC maintenance and differentiation during development. It has been previously proposed that apical-endfoot cadAJs promote Notch signaling in NPCs (*43–45*), but the mechanism underlying precise Notch signal regulation was unclear. Our study suggests that Notch signaling is maintained by the size-dependent protein segregation mechanism, and disruption of the cadAJ-mediated spatial switch downregulates Notch signaling and hence promotes NPC differentiation.

The spatial switch described here is highly analogous to the kinetic segregation model of T cell activation, where the large CD45 phosphatase is excluded from T cell receptor (TCR) immunological synaptic clefts (*10–12*). However, there are several distinct features of the Notch spatial switch compared to the kinetic segregation model. First, unlike the immunological synapse where TCR and CD45 remain constant in size throughout activation, Notch undergoes a dramatic size change during the course of cell surface activation, enabling its dynamic spatial redistribution and sequential proteolysis. Second, the role of cadAJs in Notch signaling is not limited to creating a physical barrier, but also plays the critical role of recruiting γ-secretase to facilitate processing of S2-cleaved Notch at the cell surface. Third and finally, the consequences of size-dependent segregation on signaling are reversed in comparison to the immunological synapse. While spatial segregation of CD45 enables sustained TCR phosphorylation and downstream signaling, Notch segregation from cadAJs inhibits signal activation. Our result extends the relevance of size-dependent spatial segregation models beyond immune cells (*10–12, 50*), supporting the notion that size-dependent protein segregation can serve as a general mechanism for regulating a broad range of receptor signaling at the cell-cell interface, including Notch and APPs. It is also important to note that our model may not be limited to the cadAJs, but may be extended to other cell-cell junctions that provides an environment for size-dependent protein segregation while effectively concentrating proteases.

Overall, a spatial switching mechanism based on size-dependent protein segregation not only sheds light on the mechanism underlying the sequential proteolysis of Notch and APPs, but also may extend to other receptors processed by γ-secretase. Finally, we anticipate further implications of our work in other areas of research such as providing new design principles for synthetic receptors like synNotch, as well as new therapeutic approaches that target Notch and APP signaling by spatial mutation in cancer and neurodegenerative diseases.

## Acknowledgement

The authors thank Drs. S. Blacklow (Harvard U.), C. Miller (King’s College London), and K. Shimamura (Kumamoto U.) for the kind gifts of Notch, APP, and DN-cadherin plasmids, respectively. We also thank Drs. A. Balmain, M. Moasser, and E. Collison (UCSF) for sharing cell lines. Dr. Daniel Fletcher (UC Berkeley), Mr. Ari Joffe (UC Berkeley), and Dr. Duaa Al-Rawi (Stanford U.) provided insightful discussion. For reagents, technical support, and discussions we thank the Kim, Cheon, Gartner, and Jun laboratories, as well as the Nikon Imaging Center and Wynton at UCSF.

## Author contributions

M.K., K.M.S., Z.J.G., and Y.J. conceived the ideas and designed research; M.K. and K.M.S. constructed plasmids, generated cell lines, and performed confocal microscopy. M.K. performed mechanogenetics, truncation study, spatial mutation, immunoblot analysis, reporter cell assay, and APP experiment. K.M.S. performed Notch exclusion and activation experiments, designed truncation study, wrote custom python image analysis scripts. W.R.K. performed animal experiment, flotillin staining, and mechanogenetic experiment. N.K. performed coarse-grained MD simulation. R.G. generated cadherin-KO cells. M.A. synthesized magnetic nanoparticles. H.L. helped confocal imaging/western blot analysis, respectively. M.K.K. performed Elisa analysis of Aβ secretion from CDH KO cells. S.H.C. helped western blot analysis. J.F. and D.S. performed initial proof-of-concept experiments. A.G. and N.K.R. provided anti-PS1 antibodies. M.L.K. helped VE-cad experiment, H.K., Y.H.K., and J.C. oversaw CRISPR-Cas9 KO experiment, MD simulation, and magnetic nanoparticle synthesis, respectively. Z.J.G. oversaw and supervised all spatial mapping and Notch truncation experiments. Y.J. oversaw and supervised all aspects of the study. M.K. and K.M.S. analyzed data. M.K., K.M.S., Z.J.G., and Y.J. wrote the manuscript.

## Data and materials availability

All raw images, immunoblot gel images, and analyzed data are available on request. Additional data that support the findings of this study are available from the corresponding author upon reasonable request.

## Supplementary Materials

Materials and Methods

Table S1

Figure S1-S19

References (51-80)

Movie S1-S2

