## Supplementary Materials for "Size-dependent protein segregation creates a spatial switch for Notch signaling and function"

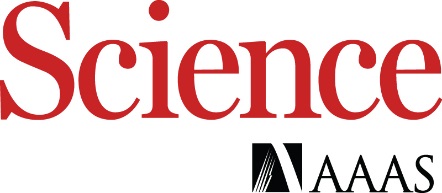

Supplementary Materials for

**Size-dependent protein segregation creates a spatial switch for Notch signaling and function**

Minsuk Kwak^1,2,3,4,5,6,^*, Kaden M. Southard^1,2,^*, Woon Ryoung Kim^1,2,3,^*, Nam Hyeong Kim^1,2,3,6,7^, Ramu Gopalappa^4,8^, Minji An^4,5^, Hyun Jung Lee^1,2,3^, Min K. Kang^9^, Seo Hyun Choi^4,5^, Justin Farlow^2^, Anastasios Georgakopoulos^10^, Nikolaos K. Robakis^10^, Matthew L. Kutys^11^, Daeha Seo^12^, Hyeong Bum Kim^4,5,8,13,14^, Yong Ho Kim^6,7^, Jinwoo Cheon^4,5,15^, Zev J. Gartner^2,16,†^ , Young-wook Jun^1,2,3,4,5†^

*These authors contributed equally.

**This PDF file includes:**

Materials and Methods

Supplementary Text

Figs. S1 to S19

Tables S1

Captions for Movies S1 to S2

**Other Supplementary Materials for this manuscript include the following:**

Movies S1 to S2

Materials and Methods

Plasmid construction

Plasmid constructs used in this study are listed in **Table S1**. All constructs used in this paper were assembled using standard restriction enzyme-based cloning, in-fusion cloning, and/or Gibson isothermal assembly. The maps, sequences, and construction details of all plasmids are available upon request. All constructs were sequenced to confirm mutation. Complete details of all cloning procedures are available upon request.

Flag-human Notch1 (N^FL^)-Gal4 and pGF1-UAS-H2B-mCherry were gifts from S. Blacklow (Harvard University). Flag-human N^FL^-Gal4 was provided in a Tet-ON Flp-IN vector (pcDNA5). SNAP-N^FL^-mCherry and SNAP-N^FL^-Gal4 were constructed as previously reported (*9*). All Notch1 variants with partial or full extracellular domain truncation were constructed by linearizing and amplifying SNAP-hN1-mCherry vector via inverse PCR while omitting the sequence corresponding the ECD truncation. Notch ectodomain sequences of amino acid 23-981, 23-1426, 23-1709 were deleted for SNAP-ΔEGF_1-25_-mCherry. SNAP-ΔEGF-mCherry. and SNAP-NEXT-mCherry, respectively. Note that similar Notch variants with partial ECD truncation were reported previously (*47*) where the structural integrity and function of Notch negative regulatory region (NRR) domain were preserved. Ecad-GFP was purchased from Addgene (Addgene plasmid # 28009; <http://n2t.net/addgene:28009>). SNAP-Ecad-GFP and Halo-Ecad-GFP were constructed first by linearizing and amplifying Ecad-EGFP vector via inverse PCR. SNAP- and Halo-tags were then inserted in frame with E-cadherin, downstream of the E-cadherin pro-peptide (amino acid #155) and upstream of the extracellular domain sequence, using Gibson assembly (NEB). pCMV6-Flotillin-1-Halo (Flot1-Halo) was constructed by replacing the myc-tag within the pCMV6-Flotillin1-myc (purchased from Origene (MR206823)) with the Halo-Tag sequence in frame with Flot1 using In-Fusion cloning (Clontech). Human amyloid precursor protein1-EGFP (APP-EGFP) was created by cloning human APP695 (a gift from C. Miller of King’s College London) into a pEGFP-N1 plasmid (Clontech). To facilitate membrane distribution mapping, we used APP constructs used in confocal imaging lack YENPTY motifs (**Fig. 5, F and G**), and compared the result with full-length APP (**fig. S18**). APP^ΔYENPTY^-EGFP was constructed by first linearizing the vector via inverse PCR, and deleting the sequence of final 15 amino acids upstream of C-terminus of APP (amino acid 681-695), which includes YENPTY motif (amino acid 682-687) using Gibson assembly.

### Tissue culture

Human U2OS cells were cultured in McCoy’s 5A media supplemented with 10% heat-inactivated FBS (Life Technologies) and 1% Pen-Strep (Life Technologies) and passaged with 0.05% trypsin-EDTA. MDCK cells were grown in Eagle's Minimum Essential Medium (UCSF cell culture facility) supplemented with 10% heat-inactivated FBS and 1% Pen-Strep. MDCK cells were lifted by treating for 10 minutes with PBS Ca^2+^ and Mg^2+^ free with 0.05% EDTA followed by trypsinization. HUVECs were purchased from ATCC (ATCC® CRL-1730™) and grown in Endothelial Cell Growth Medium-2 (EGM-2, Lonza CC-3162) for up to 4 passages (0.05% trypsin-EDTA) after thawing. Human immortalized keratinocyte cell line, HaCaT, were purchased from ATCC (ATCC® PCS-200-011) and grown in DMEM (Gibson) supplemented with 10% FBS and 1% Pen-Strep. All cell lines were maintained at 37°C in a humidified incubator with 5% CO_2_ and passaged every 2-3 days, depending on confluency, using 0.05% Trypsin-EDTA (UCSF cell culture facility). For polarized MDCK culture, cells were for use on a transwell filter (0.4 µm, collagen coated) at high density.

Transfection and cell line generation

All cell lines expressing recombinant proteins used in this study are listed in **Table S1**. U2OS stable cell lines were constructed from parental U2OS T-rex cell lines (Flp-IN, Tet-ON engineered cell line, gift from S. Blacklow). Constructs were inserted into the engineered Flp-IN site by co-transfection with a plasmid containing the Flp-recombinase (pOG44) via electroporation with the Neon Transfection System (ThermoFisher) according to manufacturer’s protocol (Shock Conditions: 1230V, 10ms, 4 pulses, number of cells 5 x 10^6^). The amount of total DNA used was 10 µg/well: 1 µg of DNA containing the desired construct and 9 µg pOG44. Cells transfected with desired plasmids were incubated in a selection medium containing 400 µg/mL hygromycin (Invitrogen) for at least 10 days. All cells with Notch truncation and reporter were further sorted for inducible expression of Notch variants via fluorescence-activated cell sorting (FACS) on a FacsAria2 (BD) by staining for the appropriate tag (SNAP or Halo) with fluorescently tagged antibody. For single-cell monoclonal population establishment, fluorescently-positive bulk-sorted populations were plated into 96 well plates at 0.2 cells/well by serial dilution and grown in selection medium. Each clonal cell population was tested and selected based on the levels of Notch reporter activity or Notch membrane expression. U2OS cells expressing recombinant proteins transiently were generated by transfecting plasmids encoding desired proteins using Neon-based electroporation. Cells were allowed to settle in a 6-well cell culture dish post electroporation for 6-8 hr. To remove dead cells, cells were lifted and re-plated on a fibronectin-coated glass bottom dish with 1 x 10^5^ cells per well density. MDCK cells were plated at 70% density then transfected with N^FL^-mCherry utilizing Lipofectamine 3000 (ThermoFisher) or Neon electroporation according to the manufacturer’s protocol (Shock Conditions: 1,650V, 20ms, 1 pulse, number of cells 5 x 10^6^). HUVEC cells were transfected via electroporation with SNAP-N^FL^ via the BioRad Gene Pulser system **(**250 V, 20 ms square wave, 1x10^6^ cells/mL Gene Pulser Electroporation Buffer, 5 µg/mL SNAP-N^FL^-mC). All cells transiently expressing recombinant proteins were incubated for 24-48 hr from the transfection, and then used for further analyses.

Live cell mechanogenetics experiment

Mechanogenetics experiments were performed as previously described in Kwak et al., 2019 with some modifications (*9*, *20*, *21*). Monovalent magnetofluorescent nanoparticles (MFNs) were synthesized as previously described (*20*).

*Micro-magnetic tweezers (μMT) set up*. The μMT was set up and aligned on the inverted microscope with point-scanning confocal imaging capabilities (Nikon) as previously described (*9*, *20*, *21*). The needle probe – NdFeB magnet assembly was attached to the z-translation stage (Sutter Instrument, MP-325) and its location was carefully aligned with the microscopic objective lens while observing the dummy substrate filled with DPBS. The µMT tip was positioned at the center of the objective oculus with bright-field illumination using the X-Y translation stage linked to PIMikroMove (Physik Instrumente) and µManager (UCSF). Using the z translation stage, the µMT was carefully lowered to set the height of the tip to 10 µm above the focal plane while recording the X-Y coordinates and the z-position of the needle probe.

*Preparation of cells expressing recombinant Flotilin-1 for mechanogenetics experiments*. U2OS cells were co-transfected with SNAP-Ecad-GFP (5 µg) and Flot1-Halo (5 µg) plasmids using Neon electroporation. 24 hr later, cells were re-plated on a #1.5 glass-bottomed dish (MatTek, d = 10 mm) coated with collagen at a density of 1 x 10^5^ cells per dish. To fluorescently label Flot1-Halo, cells were treated in a complete McCoy’s 5A medium containing 3.5 μM cell membrane permeable Halo-ligand 660 dye (Promega) for 30 minutes at 37°C. Cells were washed three times with DPBS, incubated with a phenol red-free complete medium, and then mechanogenetically stimulated (see below). For the cholesterol depletion experiment, we also treated cells 10 mM of methyl-β-cyclodextrine (MβCD) (Sigma-Aldrich) in serum-free McCoy’s 5A medium for 30 min at 37°C and washed with complete medium three times. To label SNAP-Ecad-GFP with MFNs, cells were first treated with 5 µM of an oligonucleotide bearing benzylguanine (BG-T_60_ACTG_10_) for 45 minutes at 37°C, washed two times with 10 ml of serum free medium, and then incubated with serum-free medium containing 10 nM monovalent MFNs bearing complementary sequence (T_60_CAGT_10_) and 0.5% alkali casein for 10 min at 37°C, 5% CO_2_. Cells were washed with 10 ml of complete medium two times, and then incubated with phenol red-free medium for mechanogenetic experiments on a confocal or wide-field epifluorescence microscope.

*Preparation of cells expressing human Notch1 receptor for the mechanogenetic experiment*. Inducible U2OS cells stably integrated with SNAP-N^FL^-mCherry were transfected with Halo-Ecad-GFP (10 µg) using Neon electroporation. 24 h later, cells were re-plated on a collagen (or fibronectin)-coated glass-bottomed dish. To induce surface expression of SNAP-N^FL^-mCherry. cells were incubated with complete medium containing doxycycline (Sigma, 2 µg/mL) for 18 h. To inhibit γ-secretase activity, cells were treated with DAPT (5 µM) and further incubated for 6 h. Cells were treated with 5 µM of an oligonucleotide bearing chloroalkane (Cl-T_60_ACTG_10_) for 45 minutes at 37°C and labeled with MFNs via the procedure described above.

*Mechanogenetic regulation of artificial E-cadherin junctions*. To induce MFN and hence cadherin clustering, the μMT was carefully directed towards a targeted subcellular location until the tip-to-membrane distance (*d*) reached 10 µm. As the tip approached the target membrane, the formation of an artificial E-cadherin junctions (cadAJs) was monitored every 5 minutes. After 30 min of mechanogenetic stimulation, the spatial distribution of MFNs and artificial cadAJs was monitored using time-lapse confocal fluorescence imaging. To investigate γ-secretase processing of full-length Notch, the spatial distribution of membrane mCherry (S-N^FL^-mC) or nuclear mCherry signals (S-N^FL^-Gal4) were monitored using time-lapse live cell confocal imaging. To observe localization of membrane microdomains, the spatial distribution of Flot1 fluorescence signal was monitored using live cell confocal imaging. Time-lapse live cell confocal imaging was performed using a 60x Plan-Apo oil objective (NA 1.4) on a Nikon A1 laser scanning confocal microscope equipped with an environmental chamber maintaining cells at 37°C, 5% CO_2_. Cells were immediately fixed with 4% paraformaldehyde (Life Technologies) in DPBS for 15 minutes and washed with DPBS 3 times for 5 minutes before immunostaining.

### Fluorescence labeling and Immunostaining

*Fluorescence labeling of cells expressing SNAP- or Halo-tag proteins.* Cells expressing SNAP- and/or Halo-tagged fusion proteins were labeled with BG- and/or chloroalkane functionalized fluorescence dyes, respectively. Dox-inducible cell lines grown on a collagen I-coated substrate were treated with doxycycline (2 µg/ml) 24 hr before labeling. Cells with transient receptor expression were labeled with dyes 48 hr post-transfection. Dye-labeling was performed by treating the cells with 5 μM fluorescence dye in serum containing media for 30 min. Cells were then washed 3 times with complete media. For live cell imaging, cells were incubated with phenol-red free complete media. For imaging of fixed cells, cells were washed with PBS, fixed with 4% PFA in PBS for 10 min, and then washed thoroughly with PBS.

*Immunofluorescence staining.* Fixed cells were permeabilized with 0.5% tween-20 diluted in PBS for 15 minutes, then blocked by incubation with blocking buffer (5% normal goat serum and 1% BSA in 1xPBS) for 1 hr in room temperature. For immunostaining of surface ADAM10 and ADAM17 expression, the sample was directly blocked without a permeabilization step. Cells were incubated overnight at 4°C or 2 hr at 25°C with the primary antibodies: mouse monoclonal anti-ADAM10 (1:100; Santa Cruz Biotechnology), mouse monoclonal anti-ADAM17 (1:200, R&D Systems), rabbit polyclonal anti-PS1 (*19*) (1:50), rabbit monoclonal anti-Nicastrin (1:200, Santa Cruz Biotechnology), mouse polyclonal anti-Paxillin (1:500; BD Bioscience), rabbit polyclonal anti-myc tag (1:200; abcam), mouse monoclonal anti VE-cadherin (1:400, BD Biosciences), and Alexa Fluor 488 Phalloidin (1:400 dilution of 200 units/mL stock ; life technologies). All antibodies were diluted in the 0.5x blocking buffer (2.5% normal goat serum and 0.5% BSA in 1xPBS). Following primary antibody incubation, cells were washed in PBS for 5 min four times and incubated with Alexa Fluor 405-conjugated goat anti-mouse (1:400, PS-1), Alexa Fluor 594-conjugated goat anti-mouse (1:400; Paxillin), Alexa Fluor 488-conjugated goat anti-mouse (1:400; VE-cadherin), and Alexa Fluor 647-conjugated goat anti-mouse (1:500, ADAM10), as appropriate. Nucleus staining was performed using Hoechst 33342 (ThermoFisher) diluted in 1xPBS (1 µg/mL) for 15 min and then washed once. Cells were incubated at room temperature in the dark for 1 hr, washed with PBS for 5 min three times, and imaged by epifluorescence or confocal microscopy. Epifluorescence microscopy was performed with 60x Apo, 1.40 NA or 100x Apo, 1.49 NA oil objectives (Nikon) on a Nikon Ti Eclipse microscope equipped with OBIS solid-state lasers (488, 552, and 647 nm, Coherent Inc.), a 300W Xenon lamp (Sutter Instrument, Lambda LS), a motorized stage (ASI, MS-2000), and a temperature- and CO_2_-controlled stage top incubator (Okolab, Bold Line). Unless otherwise noted, confocal microscopy was performed using Plan-Apo 60x, 1.4 NA or Plan-Apo 100x, 1.4 NA oil objectives (Nikon) on a Nikon A1R laser scanning confocal microscope. Images were acquired using Galvano scanning mode and confocal zoom of 3-4x magnification.

Monitoring the dynamic spatial localization of Notch intermediates

To activate Notch, we plated cells expressing SNAP-N^FL^-mCherry on a substrate coated with Dll4 fused with a Fc fragment (Dll4-Fc) (*51*). Briefly, a glass bottom dish (Lab-Tek II Chambered Coverglass, ThermoFisher, or 7-mm glass-bottomed dish, MatTek) was coated with fibronectin (Hamster, 5 µg/ml) and Dll4-Fc (2.5 µg/ml) for 1 hr at 37°C, and washed thoroughly with 10 ml of PBS. A negative control dish was also prepared by coating it with fibronectin only. U2OS cells co-expressing SNAP-N^FL^-mCherry and Ecad-GFP were plated and incubated with doxycycline (2 μg/mL), TAPI2 (100 μM) (*52*), and/or DAPT (5 μM). Different combinations of inhibitors were used to capture the respective intermediates (See **Fig. 2B**). After 48 hr, cells were labeled with SNAP-647 (NEB, 5 µM) and then fixed as detailed above. Inhibitor concentrations were maintained during wash and fixation steps. For DAPT washout experiments cells were plated and activated via Dll4-Fc ligand as described above. DAPT inhibitor was removed by washing in media at each time point (0, 0.5, 1.5, 3, 6, and 12 hrs). At each time point, cells were washed three times with large volumes of PBS and fixed 4% with PFA. The spatial distribution of Notch intermediates and cadAJs was monitored using spinning disk confocal fluorescence microscopy (Zeiss Cell Observer Z1), equipped with Yokagawa spinning disk and Evolve 512 EMCCD Camera (Photometrics). Images were obtained with Plan-Apo 63x, 1.4 NA or Plan-Apo 100x 1.46 NA oil objectives (Zeiss) with solid-state lasers of 405, 488, 561 nm, and 647 nm. The microscope was controlled with Zeiss Zen software (Zeiss).

Plasma membrane staining using DiI dyes

U2OS cells expressing Ecad-GFP were plated on a fibronectin-coated glass bottom dish (MatTek, D = 7.0 mm) at a density of 1 x 10^4^ cells per dish. After 48 hr, the dish was filled with a complete McCoy’s 5A medium containing 5 µM of DiI plasma membrane labeling dyes (Invitrogen) and incubated for 10 min at 37°C, 5% CO_2_. Cells were then washed 3 times with complete medium. The spatial distribution of cadAJs and DiI membrane staining and Ecad-GFP was monitored using spinning disk confocal microscopy.

Single-cell cleavage kinetics of SNAP-NΔEGF-mC

A 6-channel µ-slide flow chamber (Ibidi, VI 0.4) was coated with fibronectin (2.5 µg/mL) for 1 hr at 37°C and washed with PBS four times. U2OS cells co-expressing SNAP-N∆EGF-mCherry and Ecad-GFP were plated on the µ-slide flow chamber by applying 60 µL of single cell suspension at a density of 3 x 10^5^ cells/mL. After 3 hr, the channel was filled with a complete McCoy’s 5A medium containing doxycycline (2 µg/mL), TAPI2 (100 µM), and DAPT (5 µM). Cells were grown for 48 hr in normal growth medium to reach 70-80% confluency and form cadherin adherens junctions. Cells were labeled with BG-Alexa Fluor 647 (NEB) for 30 min to stain cell surface N∆EGF. Multiple cells with stable cadAJs were identified using large-area epi-fluorescence scanning (500 µm x 500 µm), and the spatial distribution of SNAP-NΔEGF-mCherry at cadAJs under TAPI2 and DAPT inhibition was imaged by confocal z-stack (step size = 0.2 µm, total range of z stacks = 10 µm) scanning from basal to apical membranes. Then, DAPT containing media was removed and replaced by flowing complete medium containing doxycycline and TAPI2 at a flow rate of 50 µl/min for 10 minutes using a syringe pump. Localization of extracellular (NECD) and intracellular (NICD) domain at the cadAJs before and during DAPT washout was monitored every 30 minutes in multiple color channels (NICD, mCherry; NECD, AF647; cadAJ, GFP) by time-lapse confocal z-stack microscopy for 12 hr. Time-lapse live cell confocal imaging was performed using a 60x Plan-Apo oil objective (NA 1.4) on a Nikon A1 laser scanning confocal microscope equipped with an environmental chamber maintaining cells at 37°C, 5% CO_2_.

Western Blot analysis

U2OS cells co-expressing Notch variants and Ecad-GFP (or Halo-Ecad-GFP) were incubated with culture media containing doxycycline (2 µg/mL) and TAPI2 (100 µM) in a 6-well plate at a density of 1x10^6^ cells per well. After 24 hr, cells were washed with ice-cold DPBS twice and lysed in RIPA (Invitrogen) or 1% NP-40 (Invitrogen) supplemented with complete protease and phosphatase inhibitor cocktail (100x; Cell Signaling Technology) at 4°C while gently shaking for 30 minutes. Insoluble fractions were removed by centrifugation of the cell lysates at 13,000 r.p.m. for 10 minutes. Total protein concentrations in lysates were determined by a BCA assay (Bio-Rad). 20 µg of whole cell lysates were then mixed with 4x Laemmli sample buffer (Bio-Rad) with 10% β-mercaptoethanol (BME) and heated to 95°C for 5 minutes. For western blot analysis of DNA-crosslinked heterodimers, the cell lysates were mixed with 4x Laemmli sample buffer (Bio-Rad) without BME before boiling to denature. Samples were then loaded into a 4-15% Mini-Protein TGX precast gel (Bio-Rad) and were run at 70 V for 30 minutes and then 120 V for 45 minutes. Separated proteins were transferred to a PVDF membrane using Mini Trans-Blot Cell (100 V constant, 1 hr) or the Trans Turbo Blot system (Bio-Rad). Membranes were blocked for 1hr at room temperature in blocking solution (5% w/v nonfat dry milk in 1x TBST). The membranes were probed with anti-V1744 NICD antibody (1:1000; Cell Signaling Technology #4147), anti-SNAP (1:1000; NEB), anti-Notch1 (1:1000, Cell Signaling Technology #3447 or #4380), anti-mCherry (1:500, Abcam #167453), anti-Ecadherin (1:100, Santa Cruz Biotechnology, sc-8426), and anti-β-actin (1:5000; Cell Signaling Technology #4970) antibodies overnight at 4°C with gentle rocking. The membranes were washed in TBST three times for 5 minutes and incubated with an anti-rabbit (Cell Signaling Technology, # at 1:2000 for NICD, mCherry, SNAP detection and at 1:10000 for β-actin detection) or anti-mouse (Cell Signaling Technology, # at 1:2000 for Ecadherin detection) HRP conjugated antibody. The target proteins were visualized by chemiluminescence using an ECL detection kit and a ChemiDoc MP imaging system (Bio-Rad). Quantification of band intensities by densitometry was carried out using the Image Lab software (Bio-Rad).

Spatial mutation of SNAP-NΔEGF-mCherry via DNA crosslinking

*DNA-mediated crosslinking* of *SNAP-NΔEGF-mCherry with Halo-Ecad-GFP*. DNA crosslinkers including benzylguanine (BG)- and chloroalkane (Cl) modified oligonucleotides were synthesized as previously described(*20*, *53*). To prepare 10x crosslinking DNA stock solution, complementary BG- and Cl-modified oligonucleotides were hybridized in situ. BG-T_10_(ACTG)_5_ and Cl-T_10_(CAGT)_5_ were mixed at equimolar concentration (20 µM) in PBS, incubated at 95°C on a dry heat block for 5 min, and slowly cooled down to room temperature for 2 hr. U2OS cells co-expressing SNAP-NΔEGF-mCherry and Halo-Ecad-GFP were cultured in a 6-well plate for western blot analysis at a density of 1 x 10^6^ cells per mL or in a channel of an Ibidi µ-slide for confocal imaging analysis at a density of 3 x 10^5^ cells per mL. Cells were grown to 70-80% confluency for typically 24 hr, followed by overnight incubation with complete medium containing doxycycline (2 µg/mL), TAPI2 (100 µM) and DAPT (5 µM). Cells were then serum starved with 2 ml of serum-free medium with doxycycline, DAPT, and TAPI2 for 6 hrs. Before adding DNA crosslinkers, cells were washed and placed in 450 µl of serum-free media. 50 µl of prewarmed 10x DNA crosslinker stock solution were added to each well and incubated at 37°C. Western blot analysis to validate receptor crosslinking were performed after 30-minute incubation of the DNA crosslinkers as detailed above.

*Live cell confocal time-lapse imaging.* After overnight incubation with the DNA crosslinkers, imaging was performed on an inverted laser scanning confocal microscope (Nikon A1) equipped with an environmental chamber at 37°C and 5% CO_2_. Images were obtained with a Plan-Apochromat 60x, 1.4 NA oil objective (Nikon) with solid-state lasers of 405, 488, 561 nm, and 647 nm. Additionally, the microscope was equipped with Ti-E Perfect Focus System (Nikon). To examine the effect of DNA-mediated crosslinking on spatial distribution of SNAP-NΔEGF-mCherry at cadAJs, multiple cadAJs were imaged in entirety from basal to apical sides for Halo-Ecad-GFP and SNAP-NΔEGF-mCherry using a 488 nm and 561 nm laser respectively, for a 12 µm range at a z-step size of 0.25 µm. To monitor dissipation of SNAP-NΔEGF-mCherry at cadAJs upon removal of DAPT inhibition, fresh phenol red-free McCoy’s 5A medium containing doxycycline and TAPI2 was introduced into the channel using a syringe pump for 10 min, and confocal *z*-stack images of the previously selected cadAJs were acquired every 30 minutes for 6 hr. Images were acquired using NIS-element software (Nikon), and image post-processing and analyses were done using Fiji/ImageJ and custom-built scripts.

Spatial mutation of SNAP-NEXT-mCherry via molecular pendant addition

*Synthesis of BG-modified polyethylene glycol (PEG).* Amine-functionalized PEGs with different molecular weights and structures were purchased from Creative PEGWorks (NH_2_-PEG3.4k), Sigma (NH_2_-*b*PEG20k), and NanoCS (NH_2_-$\mathcal{l}$PEG20k) and used without further purification. BG-functionalization of PEGs was performed by amine-NHS (N-hydroxysuccinimide; NEB) coupling reaction. Briefly, NH_2_-PEG (0.5 µmol), BG-GLH-NHS (2.4 mg, 5 µmol), and N,N-dimethylaminopyridine (0.73 mg, 6 µmol; Sigma) were dissolved in anhydrous dimethylsulfoxide (DMSO). The mixture allowed to react overnight with constant shaking. Crude products were recovered by evaporating DMSO using a Speed-Vac concentrator (Vacufuge, Eppendorf), reconstituted in 500 µl deionized water, and insoluble precipitates were removed by centrifugation at 14,000 r.p.m. for 10 minutes. The BG-modified PEG was then dissolved in 200 µl in deionized water and purified by reverse-phase high performance liquid chromatography with an Agilent Eclipse XDB C-18, 5 µm, 4.6 x 250 mm^2^ column using an elution gradient of 5-75% acetonitrile in 0.02% trifluoroacetic acid.

*Synthesis of BG-modified DNA-streptavidin conjugates.* DNA oligonucleotides bearing biotin- and BG-functional groups were synthesized by reacting biotin-(ACTG)_5_-NH_2_ (IDT DNA) with BG-GLH-NHS as described above (*53*). Equimolar amounts of streptavidin (10 nmol) and BG-DNA-biotin (10 nmol) were dissolved in PBS (0.5 ml) for 2 hr, forming streptavidin-BG complex. The solution was concentrated to approximately 50 µl using an Amicon centrifugal filter (MWCO: 30k) and then diluted again with 0.45 ml of PBS. This concentration and reconstitution step was repeated three times to remove unconjugated DNA.

*Synthesis of BG-modified human IgG*. hIgG (10 mg) and BG-GLA-NHS (0.82 mg) were dissolved in 850 µl of PBS and 150 µl of anhydrous DMSO, respectively. Two solutions were mixed and reacted for 2 hr at room temperature with gentle shaking. The solution was desalted with NAP-10 and then with NAP25 pre-equilibrated with PBS. The proteins were further concentrated until the final volume is 300-500 µl using Amicon centrifugal filter (MWCO: 30k). The IgG concentration was determined by measuring the absorbance at 280 nm.

*Spatial mutation of SNAP-NEXT-mCherry using the BG-modified macromolecules.* U2OS cells co-expressing SNAP-NEXT-mCherry and Ecad-GFP were incubated in complete McCoy’s 5A medium containing doxycycline (2 µg/ml), TAPI2 (100 µM), DAPT (5 µM), and respective BG-modified macromolecules (10 µM). After 24 h, cells were fixed and imaged by confocal microscopy to determine the enrichment factor of SNAP-NEXT-mCherry at cadAJs. Images were taken with a 100x objective and 3x confocal zoom. 20 stage positions per each treatment were manually selected and their coordinates were stored in the computer. In each position, confocal z stacks of DAPI, Ecad-GFP and Notch-mCherry were acquired for a 12 µm range at a Z step-size of 0.25 µm to monitor the cadAJs in their entirety from basal to apical sides. To assess the levels of Notch activation, a set of identical experiment but without DAPT was performed. After 24 h, cells were fixed, stained with DAPI, and imaged by confocal microscopy to determine nuclear mCherry signal. Images were taken with a 60x objective and 1x confocal zoom. 5 stage positions per each condition were selected manually. For each position, a confocal large-area scan of DAPI, Ecad-GFP, and SNAP-NEXT-mCherry was acquired for a 1 mm x 1 mm area.

Plate-bound Dll4 Notch activation in high-density grouped versus solitary cells

To activate Notch, we plated SNAP-N^FL^-Gal4 reporter cells on a substrate coated with Dll4-Fc as detailed above. Two different cell seeding densities were used: We plated cells with a density of 1 x 10^3^ cells per 10 mm glass-bottomed dish (MatTek, No. 1.5 glass), predominantly yielding solitary cells. We also plated cells with a density of 1 x 10^4^ cells per dish, predominantly yielding high-density grouped cells.

*Plate-bound E-cadherin Notch activation experiment*. Glass-bottomed dishes (MatTek, #1.5, D = 10 mm) were coated with recombinant human E-cadherin-Fc (50 µg/ml, R&D systems), recombinant human Dll4-Fc (2.5 µg/ml, Sino Biological), and fibronectin (5 µg/ml, Sino Biological) diluted in PBS for 1 hr at 37°C, and rinsed with 10 ml PBS with calcium and magnesium (UCSF cell culture facility). The U2OS SNAP-N^FL^-Gal4 reporter cells were transfected with Ecad-GFP (10 µg) via electroporation, incubated overnight, and re-plated onto a fibronectin, E-cadherin-Fc, and Dll4-Fc coated glass-bottomed dish at a density of 0.3 x 10^5^ cells/ml, same as the solitary cell assay. Negative control experiment was performed with the cells plated on E-cadherin-Fc and fibronectin coated glass-bottomed dishes without Dll4-Fc coating.

*Time-lapse epifluorescence imaging.* All cells were treated with 2 µg/ml doxycycline (sigma-aldrich) at the time of plating. 2 hr post-plating, cells were imaged using time-lapse microscopy. For a high-density cell seeding assay, several groups of cells having cell-cell contacts were manually identified and their coordinates were stored. For a solitary cell assay, a number of solitary cells without any prior cell-cell contact were manually identified and their coordinates were stored. While maintaining live cells on a microscope stage with a top stage incubator, time-lapse fluorescence images were acquired in GFP and mCherry channels. In each position, the microscope (Nikon) first found focuses using the Perfect Focus System (Nikon), took a DIC image, and two fluorescent images (GFP, mCherry). To image multiple solitary cells and grouped cells in one large image, cells were first plated at a high-density (2x10^5^/ml) at the center, and after 15 min, cells were seeded at a low-density (2x10^3^/ml) over the entire substrate area. After 24 h, cells were fixed and stained for membrane and nucleus. Epifluorescence images were obtained with an inverted microscope (Nikon, Ti Eclipse) equipped with 300W Xenon lamp (Sutter Instrument, Lambda LS), a motorized stage (ASI, MS-2000), and a temperature- and CO_2_-controlled stage top incubator (Okolab, Bold Line). Images were taken with 40x (CFI Plan fluor, N.A. 1.3, Nikon) objective lens. The microscopy setup was controlled using µ-manager software.

CRISPR editing to generate E-cadherin and N-cadherin knockout mutants

CRISPR/Cas9 was used to knock out E-cadherin and N-cadherin expression from U2OS SNAP-N^FL^-Gal4 reporter cells. The genes coding for the E-cadherin (CDH1) and N-cadherin (CDH2) protein from homo sapiens (gene ID: ENSG00000039068 and ENSG00000170558) were truncated by a CRISPR/Cas9 paired sgRNAs excision strategy (*54*, *55*). For the fragment deletion of genomic DNA, we used a pair of gRNAs against the target locus of CDH1 (Exon 1 & 2 (940bp deletion) or 13 & 14 (~4712bp deletion) (**fig. S12A**) and CDH2 (Exon 1 & 2 (~29,255bp deletion) (**fig. S13A**) genes.

*sgRNA design and expression vector cloning.* Cas9 and sgRNAs were expressed using the CMV promoter-driven Cas9-2A-mRFP-2A-Puro plasmid (hereafter, Cas9-puro vector) and the hU6 promoter-driven sgRNA plasmid (Toolgen), respectively. To design sgRNAs for fragmental deletion of target loci of genes, all candidate sgRNA target sites with a protospacer adjacent motif (PAM; 5’-NGG-3’) within the coding sequence of CDH1 and CDH2 were initially identified. For efficient deletion, selected sgRNAs for the candidate target sites were evaluated with DeepSpCas9 sgRNA prediction tool (<http://deepcrispr.info/DeepSpCas9/>) (*56*). sgRNAs with high DeepSpCas9 score were selected and sgRNA oligonucleotides annealed and cloned into the vector as previously described (*57*). Sequences of the vectors and sgRNAs listed here are available upon request.

*Generation of single-cell derived knock-out clones.* For CDH1 knockout, SNAP-N^FL^-Gal4 reporter cells were transfected with plasmid mixtures containing Cas9-puro, U6-sgRNA encoding individual sgRNAs at a weight ratio of 1:2 using the Neon system. For CDH1/2 knockout, cells were transfected with plasmid mixtures containing Cas9-puro, U6-sgRNA targeting CDH1 loci, and U6-sgRNA targeting CDH2 loci at a weight ratio of 1:1:1 using the Neon system. One day after transfection, puromycin was added to the culture media at a final concentration of 2.5 µg ml^−1^. Three days after transfection, the pooled cells were analyzed for the indel efficiency of sgRNA pairs using T7E1 assay. To obtain single cell-derived clones containing the fragment deletion, we plated the cells after puromycin selection into 96-well plates at an average density of 0.25 cells/well. 14 days after plating, individual clones were isolated and analyzed using PCR and gel electrophoresis of genomic DNA to check the deletion and wild-type alleles as previously described (*58*). We next sequenced the genomic DNA of the clones containing targeted deletions to check if the two cleavage sites were joined by the generation of indels. Sequencing of genomic regions including the target sequence was performed as previously described (*59*). Briefly, PCR amplicons that included the junction regions of the deleted lncRNA target sites were cloned into the T-Blunt vector (Promega) and sequenced using universal M13FP or RP primers.

*T7E1 assay.* The T7E1 assay was performed as previously described (*60*). Briefly, genomic DNA was isolated using the Wizard Genomic DNA purification Kit (Promega) according to the manufacturer’s instructions. The region including the target site was nested PCR-amplified using appropriate primers. The amplicons were denatured by heating and annealed to allow the formation of heteroduplex DNA, which was treated with 5 units of T7 endonuclease 1 (NEB) for 20 min at 37°C followed by analysis using 2% agarose gel electrophoresis. Mutation frequencies were calculated as previously described based on the band intensities using ImageJ software and the following equation (*60*): mutation frequency (%) = 100 × (1 − (1 − fraction cleaved)^1^*^/^*^2^), where the fraction cleaved is the total relative density of the cleavage bands divided by the sum of the relative density of the cleavage bands and uncut bands.

*RT-PCR.* Total RNA was extracted from wild-type SNAP-N^FL^-Gal4 reporter (WT) cells and knockout clonal cells using TRIzol (Ambion) or an RNeasy Kit (QIAGEN), after which complementary DNA (cDNA) synthesis was performed using a DiaStarTM RT Kit (SolGent Co., Ltd.). The synthesized cDNA was subjected to quantitative PCR (qPCR) in triplicate using an Applied Biosystems StepOnePlus Real Time PCR System with PowerSYBR Green PCR Master Mix (Applied Biosystems). Gene expression was normalized to that of the CDH1 gene in WT cells. Error bars represent the standard deviation (s.d.) of the mean of triplicate reactions. Primer sequences for qPCR are available upon request.

*Notch activation assay.* WT cells, CRISPR *CDH1* knock-out cells (CDH1^-/-^), *CDH1^-/-^* transfected with E-cadherin-GFP (CDH1^-/-^ + E-cad), and *CDH1^-/-^* transfected with N-cadherin (CDH1^-/-^ + N-cad) were plated on 8 well Nunc Lab-Tek II chambered coverglass pre-coated with recombinant human Dll4-Fc (2.5 µg/ml) and fibronectin (5.0 µg/ml) as previously described. All cells were plated at a density of 30,000 cells per well. After 24 hr incubation with doxycycline (2 µg/ml), cell cytoplasm and nucleus were stained with CellTracker CMFDA dye (Invitrogen) and Hoechst 33342 (ThermoFisher), respectively. The cells were then fixed with 4% PFA for 15 minutes at room temperature and proceeded to epifluorescence and confocal imaging.

Aβ40, Aβ42, and sAPPα ELISA

1x10^6^ of wild-type or CDH1/CDH2 knockout cells were transfected with APP-mCherry (10 µg) and plated on 6-well tissue culture plate. After 48 hr incubation, conditioned media supplemented with 1x protease/phosphatase inhibitor cocktail (ThermoFisher) were centrifuged at 3,000xg 10 minutes at 4°C to remove cell debris and the supernatant were transferred to a new tube and stored at -80C. The conditioned media were analyzed for Aβ40, Aβ42, and sAPPα contents using Aβ (Invitrogen) and sAPPα (Kusa Biosci.) ELISA kits. All ELISAs were performed according to the manufacturer’s protocols. Briefly, 50 µl/well of samples, followed by 50 µl/well of detection antibodies, were applied to Aβ40, Aβ42, sAPPα coated 96-well plate, and incubated for 3 hr at room temp with shaking at 300 rpm. After washing step, 100 µl of HRP-IgG solution was applied and incubated for 30 min at room temp with shaking. After washing step, 100 µl of stabilized substrate solution was applied and incubated for 30 min at room temp with shaking. Finally, 100 µl of stop solution was applied. Absorbance at 450 nm was read and analyzed using a plate reader (Biotek Synergy 2). Wells were washed with wash buffer 4 times between each incubation step. Standard curves were generated using recombinant Aβ40, Aβ42, sAPPα provided by the manufacturer (Invitrogen).

*In vivo experiments*

***Animals.*** We used CD-1 embryonic day 13.5 (E13.5) and postnatal day three (P3) newly born mice for *in vivo* experiments. P3 pups were obtained by purchased of an untimed pregnant female mouse (E13-15) from Charles River Laboratories (Wilmington, MA) and waited for birth. All mice were housed under specific pathogen-free conditions under a 12h light-dark cycle, and all animal handling and use were in accordance with institutional guidelines approved by the University of California San Francisco Institutional Animal Care and Use Committee (IACUC).

***Retrovirus injection.*** To generate retrovirus, we used pWZL-GFP control vector (pWZL-Blast-GFP: addgene plasmid # 12269) and pWZL-dominant negative E-cadherin (pWZL-Blast-DN-E-cadherin addgene plasmid # 18800 and gift from Dr. Kenji Shimamura). GFP sequence was inserted in frame with dominant negative E-cadherin, downstream of C-terminus using In-Fusion cloning. For retrovirus production, we transfected retroviral vectors into Phoenix-Ampho cells using Calcium Phosphate transfection kit (Sigma, CAPHOS) with 50 µM chloroquine (Sigma, C6628), and collected supernatant from transfected cells after 48 h. Collected supernatant containing viral solutions was ultracentrifuged yielding concentrated solution of viral particles (25,000 rpm for 2 hours at 4°C). Approximately 10^7^ transducing units per milliliter (TU/ml) viral solution was injected into the lateral ventricular of neonatal mouse pups (P3). After hypothermic anesthesia, viral solutions (5 µl) were slowly injected using IM-9B Narishige microinjector with 2 µl/min speed. After recovery on the warming pad, the pups were placed back to the cage. After additional 2 hours, mice were subject to intracardiac perfusion fixation using 4% paraformaldehyde in PBS.

***DAPT injection.*** 10 µM DAPT was injected into neonatal mouse pups (P3) into a lateral ventricle (10 µl in each hemisphere). DMSO was injected into control mice. After 7 hours, mice were subject to fixation procedure using intracardiac perfusion of 4% paraformaldehyde in PBS.

***Immunohistochemistry.*** Mice were perfused with 4% paraformaldehyde in PBS (pH7.4) and the brains were subject to postfixation in the same fixative for 24 h. Brains were then cryoprotected in 30% sucrose in PBS, sectioned serially (20 μm) onto Superfrost plus glass slides (Fisher Scientific; Pittsburgh, PA). The brain slices were permeabilized and blocked with PBS solution containing 3% goat serum albumin and 0.3% Triton-X100, and then treated with anti-Ncadherin (1:200, Thermofisher), anti-Notch (1:200, Thermofisher), anti-PS1 (1:50) (*19*), and anti-beta III tubulin (1:500, Abcam) overnight at 4°C. The brain slices were washed three times with PBS and treated with secondary antibodies (1:1000, Thermofisher) for 30 min. Subsequently, the slices were washed with PBS, mounted and observed with a confocal microscope (Olympus, Fluoview 3000).

Image Processing and Analysis

*Cadherin junction colocalization analysis***.** Colocalization analysis was carried out in ImageJ, using thresholding to identify cadAJs and then applying the JACOP plugin to quantify colocalization using Pearson coefficient, Manders’ overlap coefficients, and cross-correlation analysis.

*Confocal 3D z-stack image processing****.*** Custom python code was used for automatic segmentation and junction intensity ratio analysis for Notch activation and truncation studies. Code is available at **(**[**https://github.com/kmsouthard/JunctionAnalysis**](https://github.com/kmsouthard/JunctionAnalysis)) In brief, resliced z-stacks of cell-cell interfaces were thresholded to identify the cadAJs and membrane Notch signal. To minimize the domination of high Notch intensity, we identified the membrane expressing Notch using a minimal threshold of membrane intensity just above background. An unbiased signal analysis window along each side the junction was selected, and the Notch membrane intensity was measured for each cell by averaging along the respective windows, while junctional intensity was measured within segments determined by cadherin junctional intensity. The ratio of junctional intensity was calculated as ratio = I_junc_ /(I_cell1_+ I_cell2_) as deviations from the expected intensity at the junction is a function of the sum of each cell’s expression level.

*Intracellular mCherry nuclear translocation analysis****.*** mCherry nuclear translocation analysis was carried out in ImageJ. GFP images were used for automated identification of cell edges and segmentation of single cells. DAPI images were used for automated identification of nucleus by implementing the Otsu thresholding method. Nuclear mCherry fluorescence data was extracted from nuclear segments by calculating the integrated fluorescence within the nucleus and subtracting a background scattering signal. In Fig. 4H, nuclear mCherry fluorescence intensities for NEXT cells treated with the macromolecular pendants were rescaled to make the intensity of N^FL^ and NEXT to 0.002 and 1.0, respectively, which are identical to the normalized band intensities of N^FL^ and NEXT measured by western blot.

*Quantification of single-cell fluorescence*. Single-cell tracking and nuclear mCherry fluorescence signal analysis of UAS-Gal4 reporter cells were performed with ImageJ, as previously described (*9*, *34*, *61*).

*Western blot quantification.* Quantification of band intensities by densitometry was carried out using the Image Lab software (Bio-Rad). Band intensities of NICD in each lane were normalized by band intensities of loading control, β-actin in the corresponding lane.

### Estimation of Protein heights

Protein heights including extended Notch height was estimated by measuring the structural size of each domain (i.e. EGF, NRR, SNAP) in Pymol (The PyMOL Molecular Graphics System, Version 2.0 Schrödinger, LLC.) and then creating an additive estimate based on the number of domains in the full-length Notch construct and each Notch truncation.

Molecular dynamics simulations

All MD simulations were conducted using the GROMACS package (*62*). Polarized MARTINI 2.2 parameters were used for the simulations (*63*). The system was composed of 54 PPCS, 54 DPPC, 108 DPPS, 288 DIPC, 216 CHOL, and 10108 water molecules with a molar composition of lipid was CHOL:PPCS:DPPC:DIPC = 3.0 : 1.5 : 1.5 : 4.0 in upper layer and CHOL:DPPS:DIPC = 3.0 : 3.0 : 4.0 in lower layer. To create immobilized lipids, we increased the mass of the phosphorus atom within DPPS by a factor of 1,000, keeping everything else the same in the parameter file. The pressure was set at 1.0 bar with a semi-isotropic parrinello-rahman coupling with compressibility 4.5 x 10^-5^ bar^-1^and the temperature was set to 295 K using nose-hoover coupling. Each system was neutralized and brought to a concentration of 0.15 M with randomly placed sodium and chloride ions. We employed the LINCS algorithm to constrain to bond lengths (*64*). A time step of 20 fs was used with an update of the neighbor list every 10 steps, which are typical values employed in MARTINI simulations. Each simulation was run afterwards for 12 µs, the last 3 µs of which was used for analysis. The MD simulations were analyzed using the in-built GROMACS tools. MDAnalysis libraries (*65*, *66*) were used for calculating diffusion constant of lipid component and *g_energy* was used for calculating interlayer interaction.

Statistical analysis.

Statistical analysis was performed in GraphPad Prism 8.0 (GraphPad) or Microsoft Excel. Figure legends indicate all statistical tests used in the figure. Unless otherwise noted in the figure legends, statistical differences were determined using Student’s t-test (two-tailed unpaired or paired t-test, depending on the experiment) when only two groups were compared or by ordinary one-way ANOVA followed by Tukey posthoc test when multiple groups were analyzed. The number of samples (‘n’) used for each experimental analysis is indicated in the figure legends. Sample sizes of sufficient power were chosen on the basis of general standards accepted by the field. In all cases, statistical significance was assumed for probability (P) < 0.05. *P<0.05; **P<0.01; ***P<0.001; ****P<0.0001; ns indicates when no significant difference was detected.

Supplementary Text

Interrogation of the mechanism underlying γ-secretase recruitment to cadAJs

We investigated how cadAJs recruit γ-secretase. Several reports have suggested possible engagement of cadAJs with spatially discrete and ordered membrane microdomains (*67*-*70*). Similarly, γ-secretase proteolytic activity is closely linked to detergent-resistant membranes (*71*-*78*). Both of these membrane features preferentially associate with membrane proteins such as Flotillin-1 (Flot1) (*70*-*73*, *79*). We therefore investigated localization of Flot1 across the cell membrane. Consistent with the artificial cadAJ experiment (**Fig. 1F and fig. S5, A and B**), we observed strong Flot1 and γ-secretase fluorescence signals at cell-cell cadAJs (**fig. S5, A to E**). These observations support the notion that both cadAJs and γ-secretase are associated with common and long-lived lipid microdomains enriched with Flot1, otherwise known to be short-lived and transient when alone (*70*). Clustering of E-cadherin triggers rapid F-actin polymerization at the cytoplasmic leaflet of the plasma membrane. Given the established interaction between F-actin and membrane constituents like phosphatidylserine that nucleate and stabilize Flot1-containing membrane microdomains, we reasoned that F-actin at the cadAJs may anchor phosphatidylserine leading to formation of the membrane microdomains (*80*). To test this notion, we performed a coarse-grained molecular dynamic (MD) simulation of a lipid membrane comprising of 1,2-dilinoleoyl-sn-glycero-3-phosphocholine (DIPC; outer leaflet), 1,2-dipalmitoyl-sn-glycero-3-phosphocholine (DPPC; outer leaflet), N-palmitoyl-O-phosphocholineserine (PPCS; outer leaflet), 1,2-dipalmitoyl-sn-glycero-3-phosphoserine (DPPS; inner leaflet), and cholesterol (Chol; inner and outer leaflet). We immobilized a portion (30%, red colored in **fig. S7A**) of DPPS in the inner leaflet to reflect its interaction with F-actin, and compared the results to another simulation set without DPPS immobilization. While both simulations showed lipid segregation within both the inner and outer leaflets, DPPS immobilization resulted in a microdomain having a strong transbilayer coupling (**fig. S7A**) (*80*). We also observed a significant decrease in lipid diffusion, indicating the microdomain stabilization (**fig. S7, B and C**).

To further confirm the role of lipid microdomains in recruiting γ-secretase to cadAJs, we tested whether cholesterol – a key component of the lipid microdomain - depletion disrupts γ-secretase localization within cadAJs. Because cholesterol depletion also destabilizes native cadAJs (*71*), we instead generated an artificial cadAJ by clustering E-cadherin while depleting cholesterol in the cell membrane by treating the cells with methyl-β-cyclodextrin (MβCD) (**fig. S7D**, See Methods). Vivid Flot1 and presenilin-1 signals were seen at the artificial cadAJ without MβCD treatment (**Fig. 1F and fig. S4, A and B**), but no presenilin-1 signal was detected at the cadAJ with MβCD (**fig. S7D**). From these observations, we confirmed that cadAJs recruit γ-secretase through its molecular association with a common lipid microdomain, rather than a direct cadherin–γ-secretase interaction.

_­­­­_
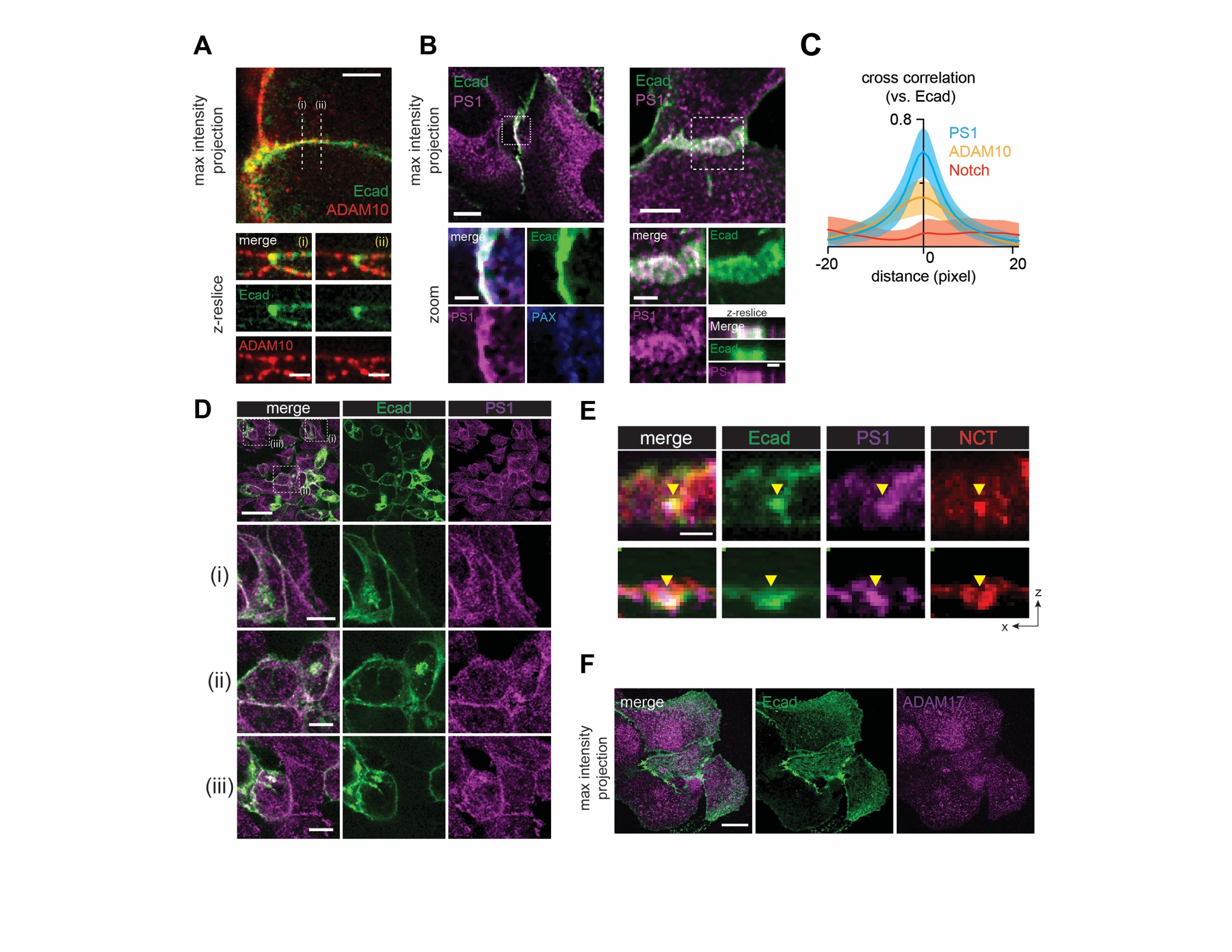

**Fig. S1. Distribution of ADAM10 and γ-secretase at the cell-cell interface. (A)** Representative confocal fluorescence images showing ADAM10 distribution relative to E-cadherin-based adherens junction (cadAJ). U2OS cells were transfected with Ecad-GFP (green), and stained with ADAM10 antibody (red). Z-resliced images along the white dashed lines. Scale bars, 5 µm (maximum projection) and 3 µm (z-resliced images). **(B)** Confocal immunofluorescence images showing endogenous presenilin-1 (PS1, magenta) distribution relative to cadAJs (green). Paxillin was also imaged as a negative control (blue) Scale bars, 10 µm, 3 µm, and 3 µm for maximum intensity projection, zoomed-in, z-resliced images, respectively. **(C)** Cross-correlation analysis of PS1, ADAM10, and Notch distributions over E-cadherin. PS1 fluorescence intensities exhibited strong positive correlation with cadAJs, while Notch showed nearly zero correlation. ADAM10 showed weak positive correlation over cadAJs. The cadAJs were set as x = 0 in the analyses. The shading along the curves represents error bars (s.e.m.; n ≥ 9). **(D)** Representative wide-field confocal immunofluorescence images showing PS1 (magenta) enrichment within cadAJs (green). Scale bar, 50 µm (low-magnification), 10 µm (zoom-in). **(E)** Confocal z-reslices of immunofluorescence images showing PS1 (magenta) and Nicastrin (NCT, red) distribution relative to cadAJs (green). Scale bar, 2 µm. **(F)** Representative confocal immunofluorescence images showing ADAM17 (magenta) distribution relative to cadAJs. ADAM17 exhibited no preferential localization relative to cadAJs. Scale bar, 20 µm.

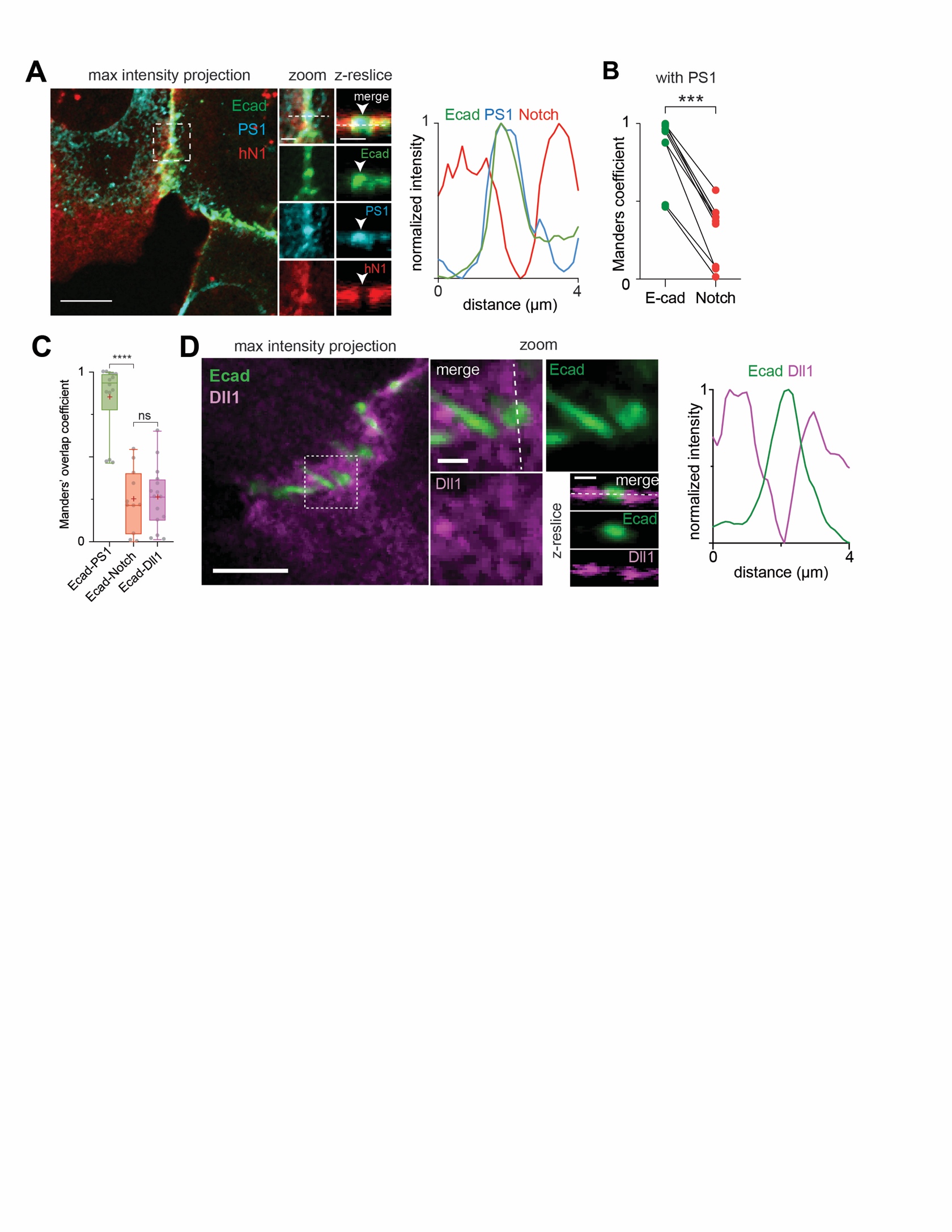

**Fig. S2. Quantitative analysis of PS1, Notch, Dll1 distribution relative to cadAJs and additional confocal images representing their spatial distribution.**

**(A)** Confocal images of U2OS cells co-expressing Ecad-GFP (green) and SNAP-N^FL^-mCherry (red), and immunostained with PS1 antibody (cyan). Line profiles show fluorescence signal intensity along the white lines. Scale bars, 10 µm, 2 µm, and 2 µm for maximum intensity projection, zoomed-in, and z-resliced images, respectively. **(B)** Paired analysis of Manders’ overlap coefficients of E-cadherin and Notch signals over PS1 in multiple cells (n = 9). ***P < 0.001. **(C)** Box-whisker plots showing Manders’ overlap coefficients (MOCs) of PS1 (cyan), Notch (red), and Dll1 (purple) relative to cadAJs. Each dot represents the MOC of a selected cadAJ. Boxes and whiskers denote the inner-quartile and full ranges. Colored lines and (+) marks indicate median and mean, respectively (****P < 0.0001; ns, not significant; n ≥ 11; ordinary one-way ANOVA with Tukey’s test). **(D)** Left: Confocal fluorescence images of U2OS cells co-expressing Ecad-GFP (green) and Halo-Dll1 (magenta). Right: Line profiles showing Ecad and Dll1 fluorescence signals along the white dashed line of the zoomed-in maximum projection image. Scale bars, 5 µm, 1 µm, and 1 µm for maximum intensity projection, zoomed-in, and z-resliced images, respectively.

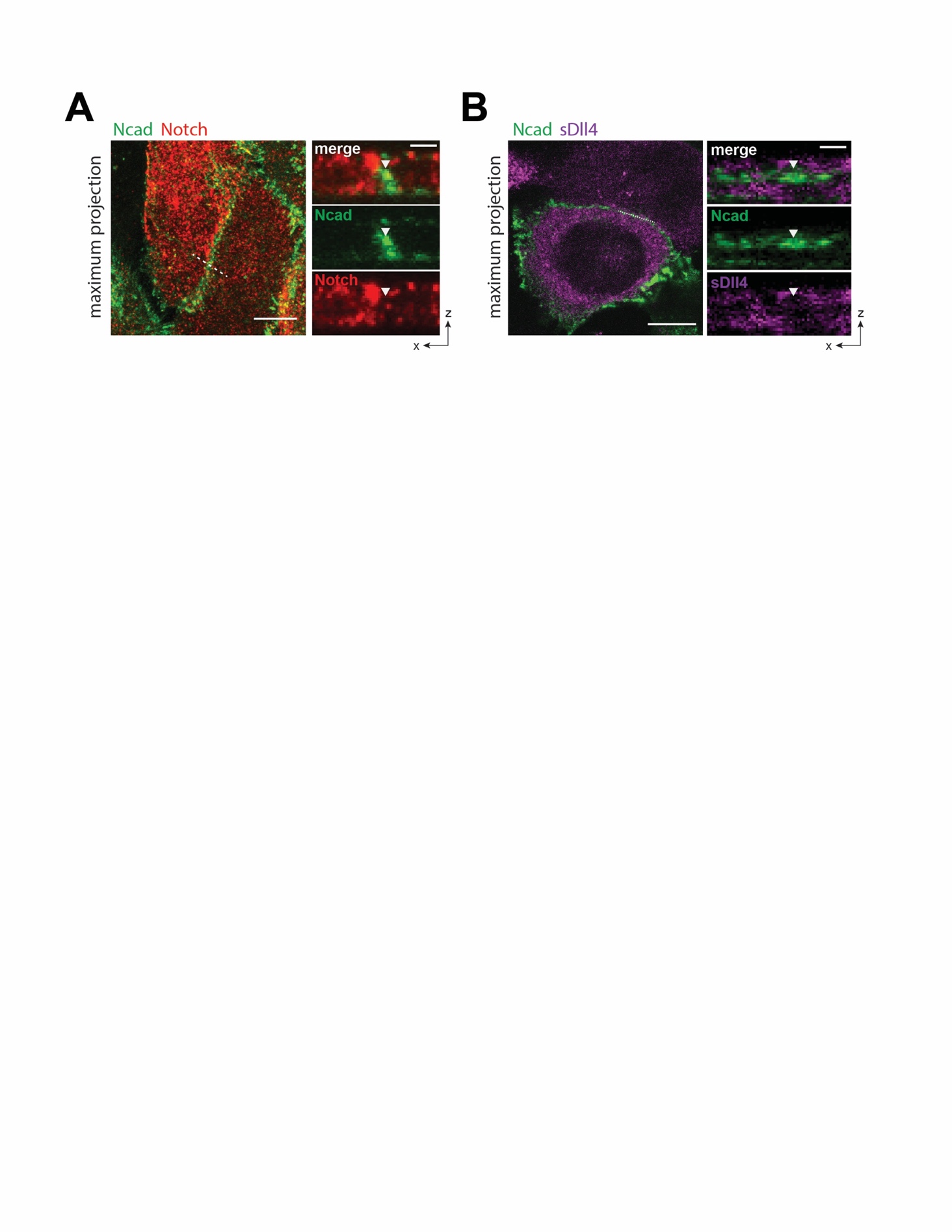

**Fig. S3. Spatial exclusion of endogenous Notch1 from cadAJs in HaCaT cells.** (**A**) Representative confocal fluorescence images of HaCaT cells immunostained with anti-Notch1 (red) and anti-N-cadherin (green). (**B**) HaCaT cells labeled with fluorescently tagged soluble Dll4 (sDll4, magenta). Left: Maximum projection image. Scale bar, 10 µm. Right: Confocal z-resliced image along the white dashed line. Scale bar, 2 µm. Both Notch1 and sDll4 were excluded from cadAJs (white arrowheads).

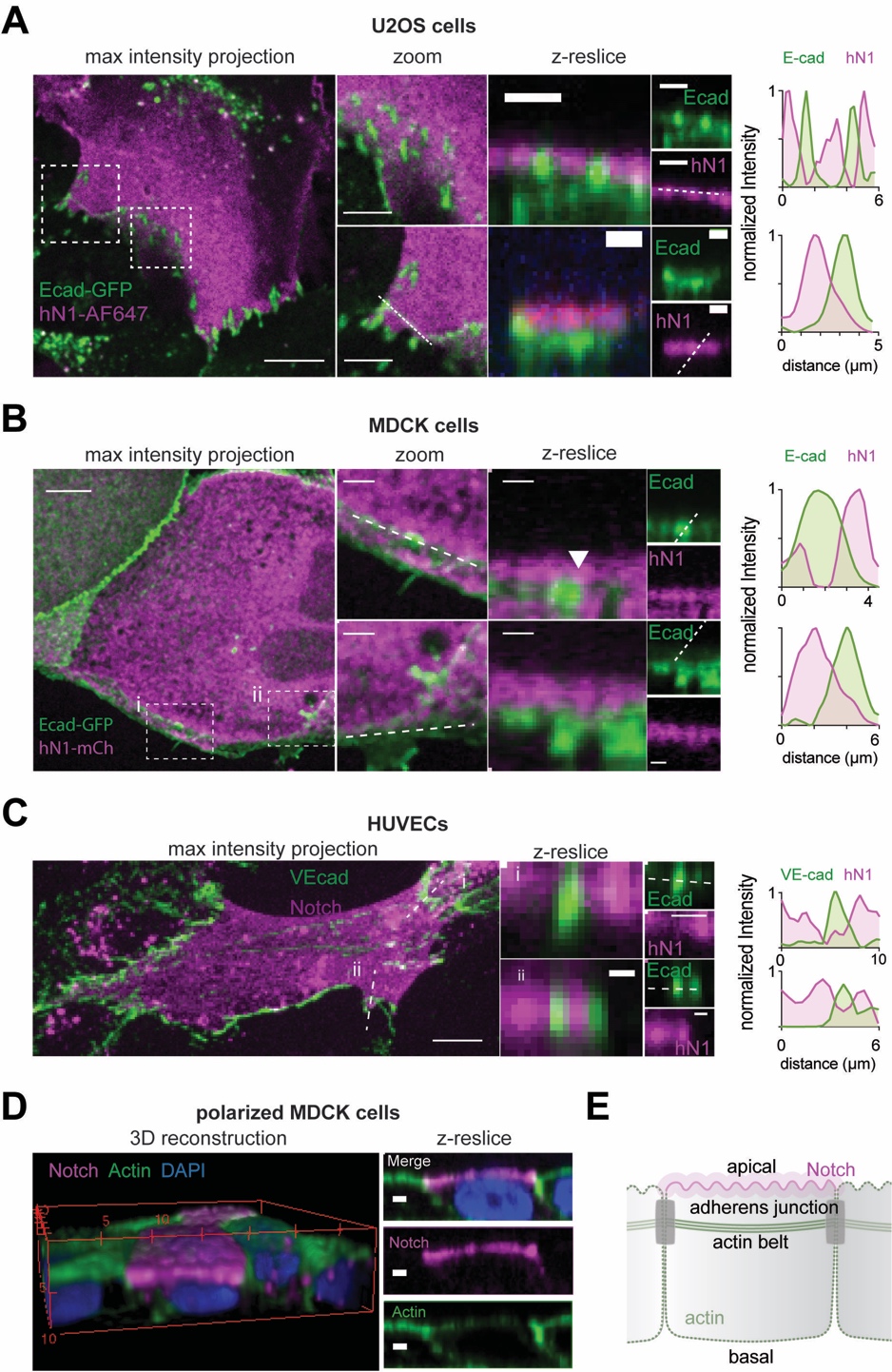

**Fig. S4. Notch exclusion from cadAJs is consistent for several cadherin family proteins, host cell types, and cell polarization states.**

A series of cell lines including U2OS, MDCK, and HUVEC were imaged. **(A and B),** U2OS (**A**) and MDCK (**B**) cells were co-transfected with plasmids encoding Ecad-GFP (green) as well as SNAP-N^FL^-mCherry. Notch receptors were labeled with BG-Alexafluor647 (magenta). Line profiles show fluorescence signal intensities along the white lines. Scale bars 10 µm, 5 µm, and 2 µm for maximum intensity projection, zoomed-in, and z-resliced images, respectively. **(C)** Confocal images of HUVECs expressing SNAP-N^F^-mCherry and immunostained with vascular endothelial cadherin (VE-cad) antibody. Line profiles show fluorescence signal intensities along the white lines. Scale bars 5 µm and 2 µm for maximum intensity projection and z-resliced images, respectively. **(D)** Polarized MDCK cells grown on a transwell filter. Notch, actin, and nucleus were immunostained with BG-AF647, phalloidin-488, and DAPI, respectively. Scale marked every 5 µm for 3D construction. Scale bar, 2 µm. **(E)** A schematic representation of Notch localization in a polarized epithelial cell.

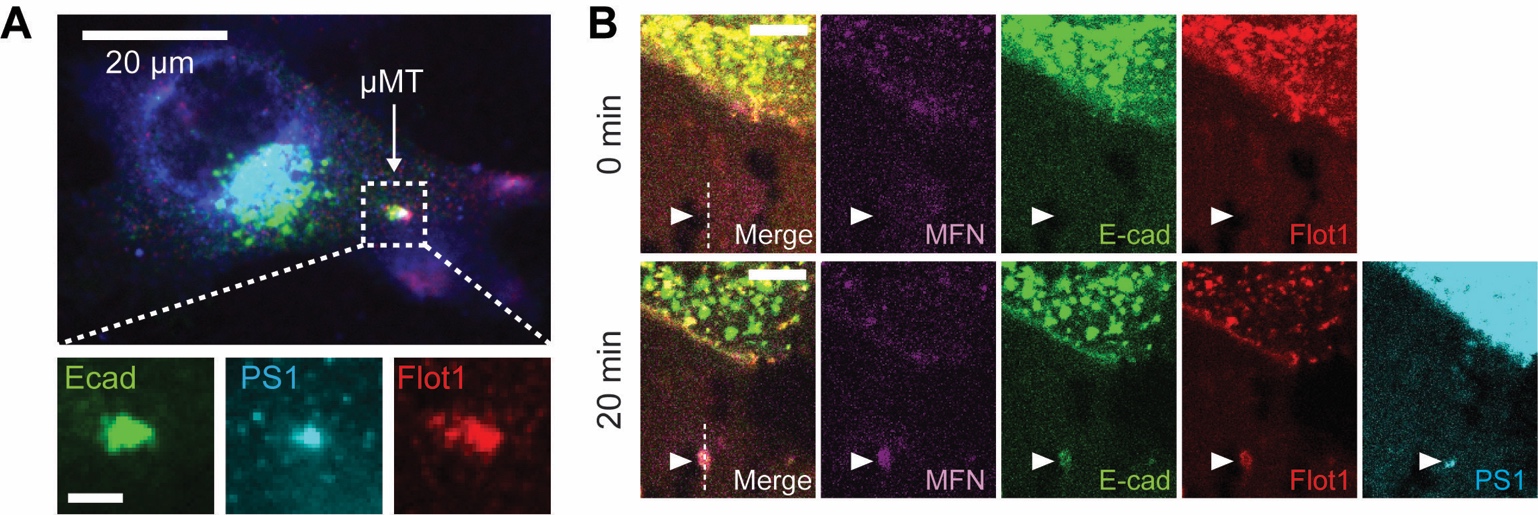

Fig. S5. Interrogation of the mechanism underlying γ-secretase recruitment into cadAJs through mechanogenetics. (A and B) Confocal fluorescence images showing PS1 and Flot1 localization at artificial cadAJs by mechanogenetics. E-cadherin and Flot1 were labeled with fluorescent tags. Endogenous PS1 was immunostained after fixation. Scale bar, 20 µm.

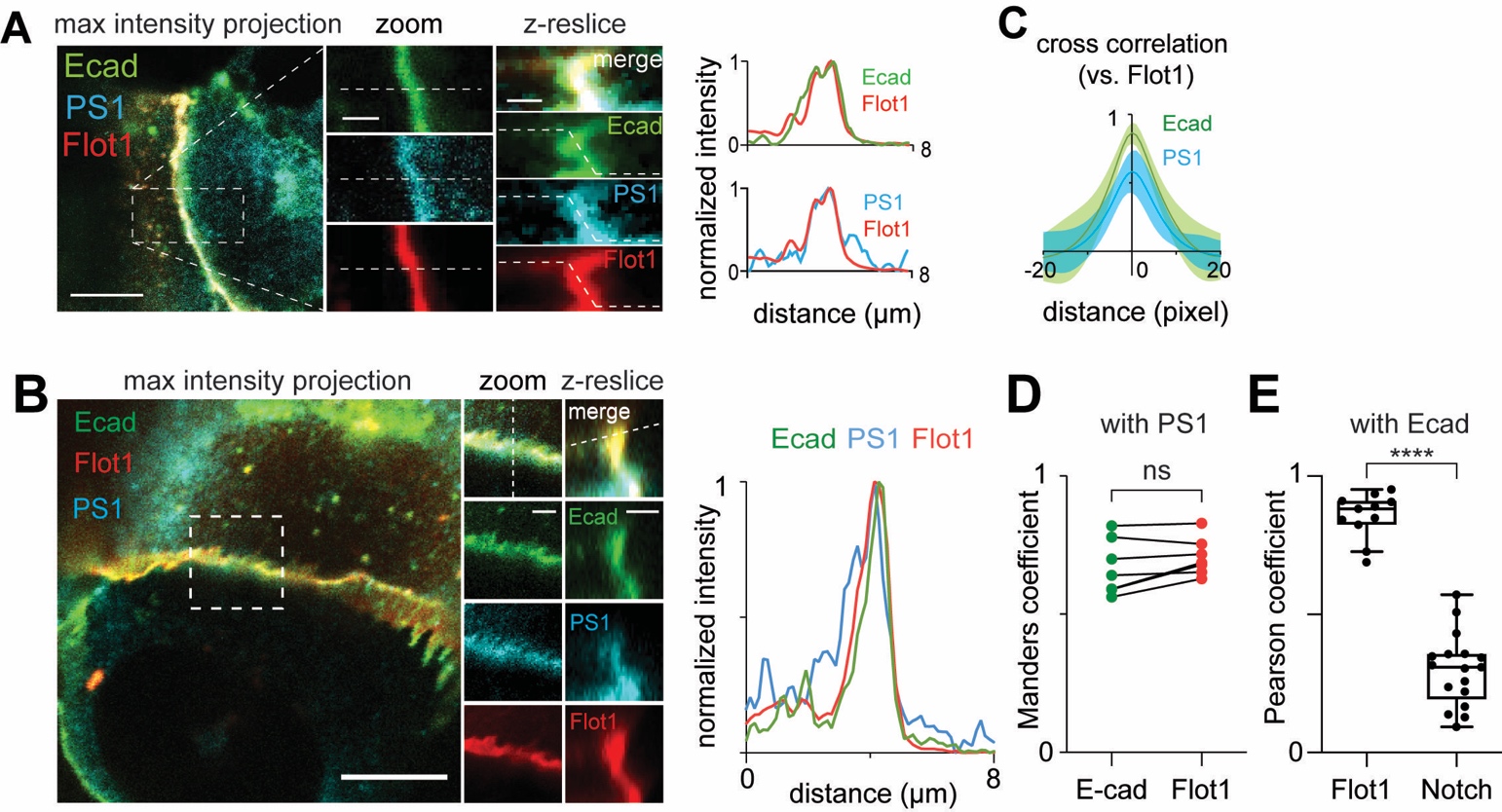

Fig. S6. Interrogation of the mechanism underlying γ-secretase recruitment into cadAJs through native cell-cell cadAJ spatial mapping. (A and B) Representative confocal fluorescence images showing the PS1 and Flot1 distribution relative to native cell-cell cadAJs. (left) A maximum projection image of merged channels. Scale bar, 10 µm. (center) Magnified images of the boxed region. Scale bar, 2 µm. (right) Z-resliced images showing the sections of the cadAJs. Scale bar, 2 µm. Line profiles of fluorescence signal from E-cadherin, PS1, and Flot1 along the white dashed lines. (C) Cross-correlation analysis of E-cadherin and PS1 over Flot1. Both Flot1 and PS1 fluorescence intensities exhibited strong positive correlation with the cadAJ. The shading along the curves represents error bars (s.e.m.; n = 7). (D) Manders’ overlap coefficients of E-cadherin and Flot-1 signals over PS1 (ns, not-significance; Two-tailed paired Student’s t test; n = 6 single cells). (E) Quantitative assessment of Flot1 and Notch colocalization with E-cadherin. Overlaid are box and whisker plots. **** P<0.0001. Two-tailed unpaired Student’s t test.

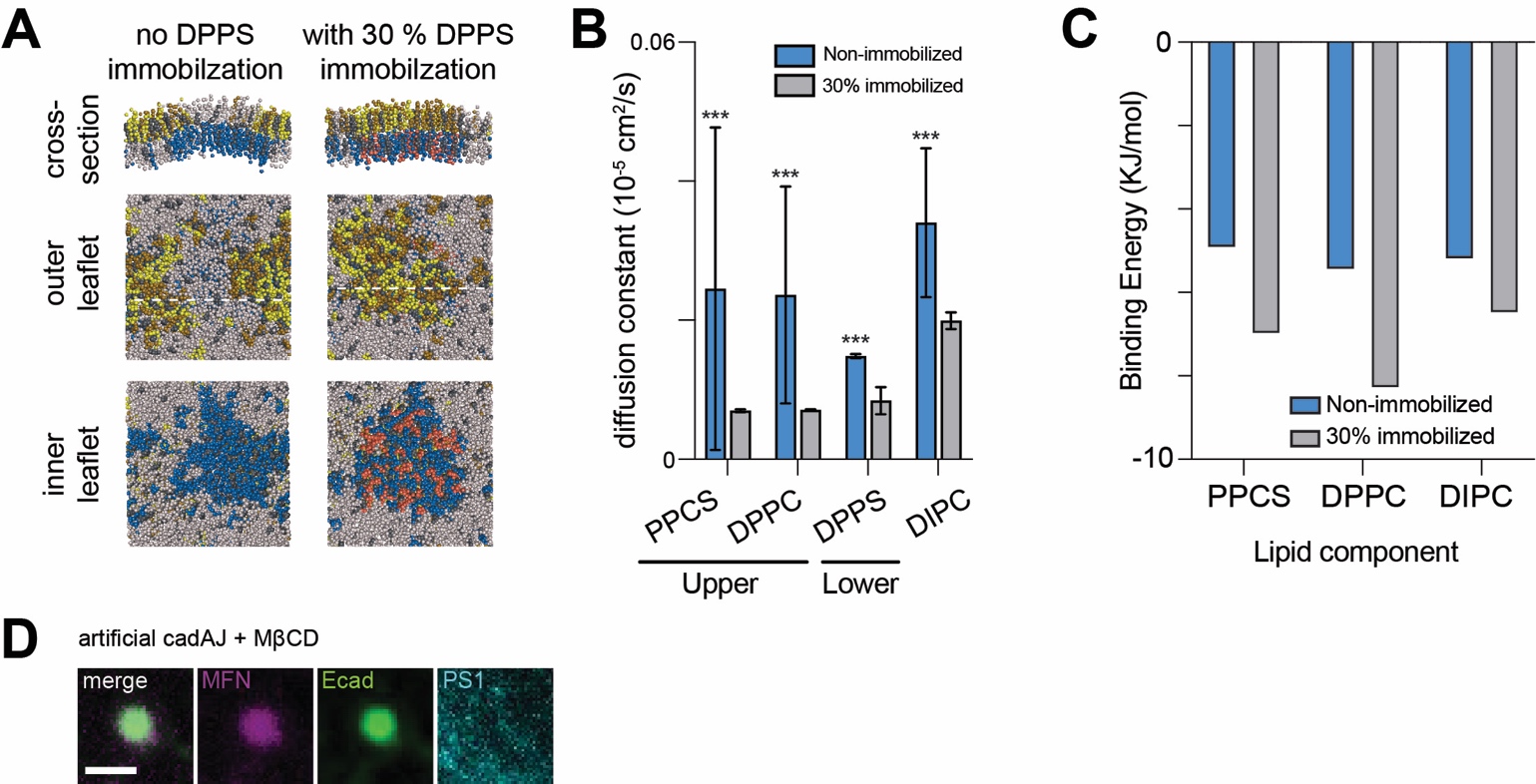

Fig. S7. Interrogation of the mechanism underlying γ-secretase recruitment into cadAJs through molecular dynamics simulation, and cholesterol depletion experiments.

(A) Snapshot images of coarse-grained MD simulation of a lipid bilayer comprising of 1,2-dilinoleoyl-sn-glycero-3-phosphocholine (DIPC; gray), 1,2-dipalmitoyl-sn-glycero-3-phosphocholine (DPPC; yellow), N-palmitoyl-O-phosphocholineserine (PPCS; green), 1,2-dipalmitoyl-sn-glycero-3-phosphoserine (DPPS; blue), immobilized DPPS (pink), and cholesterol (CHOL; grey). Left and right panels represent the simulation results without and with a partial (30%) DPPS immobilization, respectively. (B) and (C) Diffusion coefficients (B) and binding energy (C) of individual lipid components in a lipid bilayer with or without a partial (30%) DPPS immobilization. Both decreases in lipid diffusion and increased binding energy indicate lipid microdomain stabilization due to DPPS immobilization. (D) Confocal fluorescence images showing PS1 distribution at an artificial cadAJ after cholesterol depletion with MβCD treatment. No PS1 recruitment was seen at the cadAJ, suggesting that γ-secretase recruitment to the cadAJ requires lipid microdomain formation at the cadAJ. Scale bar, 2 µm.

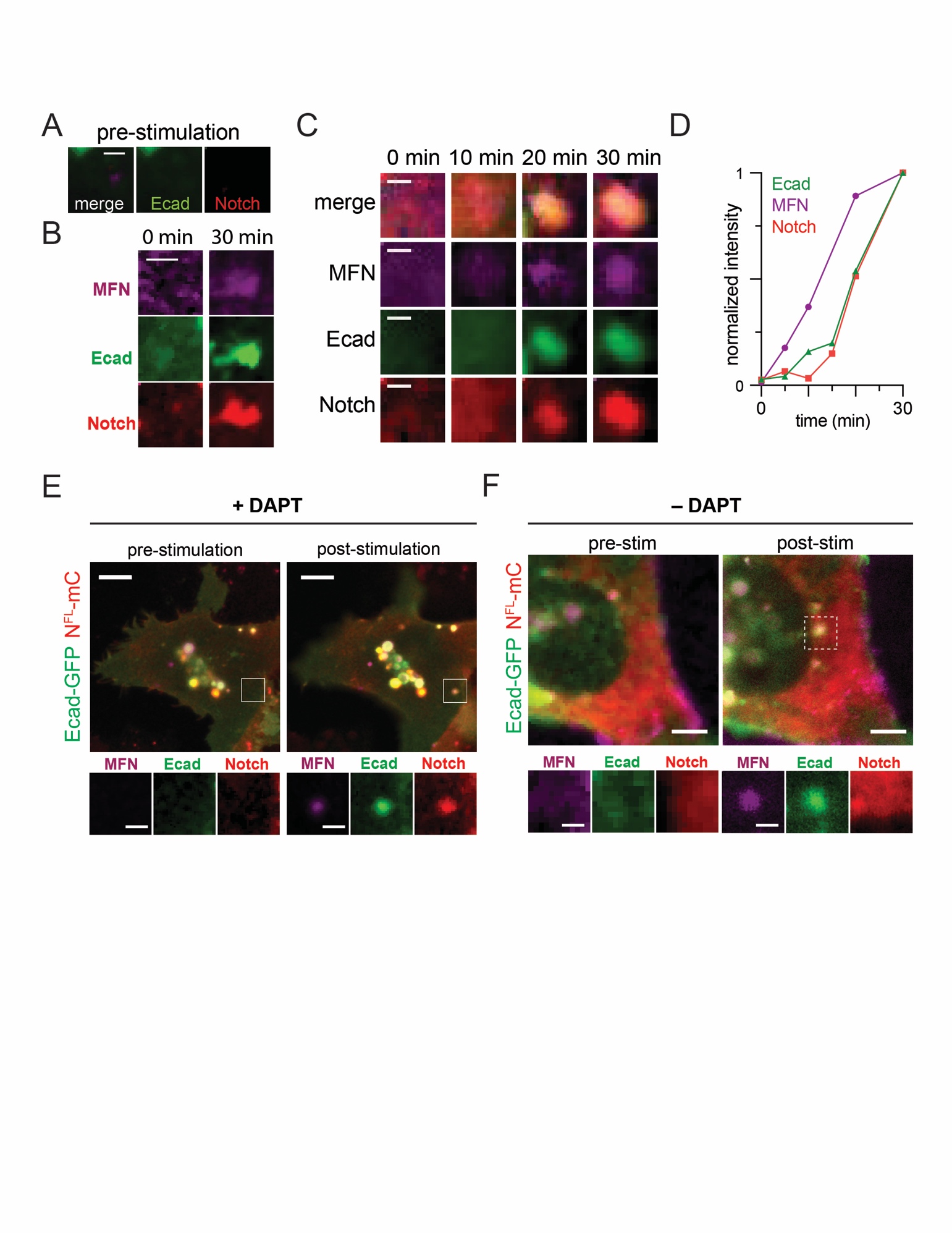

Fig. S8. Recruitment and processing of full-length Notch receptors to the artificial cadAJs generated via mechanogenetics.

(A) Pre-stimulation confocal images of a subcellular locus of the cell shown in Fig. 1G. Initial dim MFN (magenta), E-cadherin (green), and Notch (red) signals were evenly distributed. Scale bar, 2 µm. (B and C) Additional artificial cadAJs showing Notch recruitment. Time-lapse epifluorescence images (C) were acquired before micromagnetic tweezer (µMT) stimulation and then at 10, 20, 30 minutes of the µMT application. Gradual MFN and E-cadherin clustering were clearly seen, followed by Notch accumulation to the cadAJ. Scale bar, 2 µm. (D) Kinetics of signal enrichments at the artificial cadAJ shown in the panel (C). (E and F) Representative images of artificial cadAJs formed in live cells. Cells treated with both TAPI2 and DAPT (E) or with only TAPI2 but no DAPT (F). Magnified images were shown in lower panels. An intense mCherry signal was observed at the artificial cadAJ with TAPI2 and DAPT treatment, while no enrichment of Notch1 signal was seen from cells without DAPT. Scale bar, 5 µm (low-magnification), 2 µm (zoomed-in images).

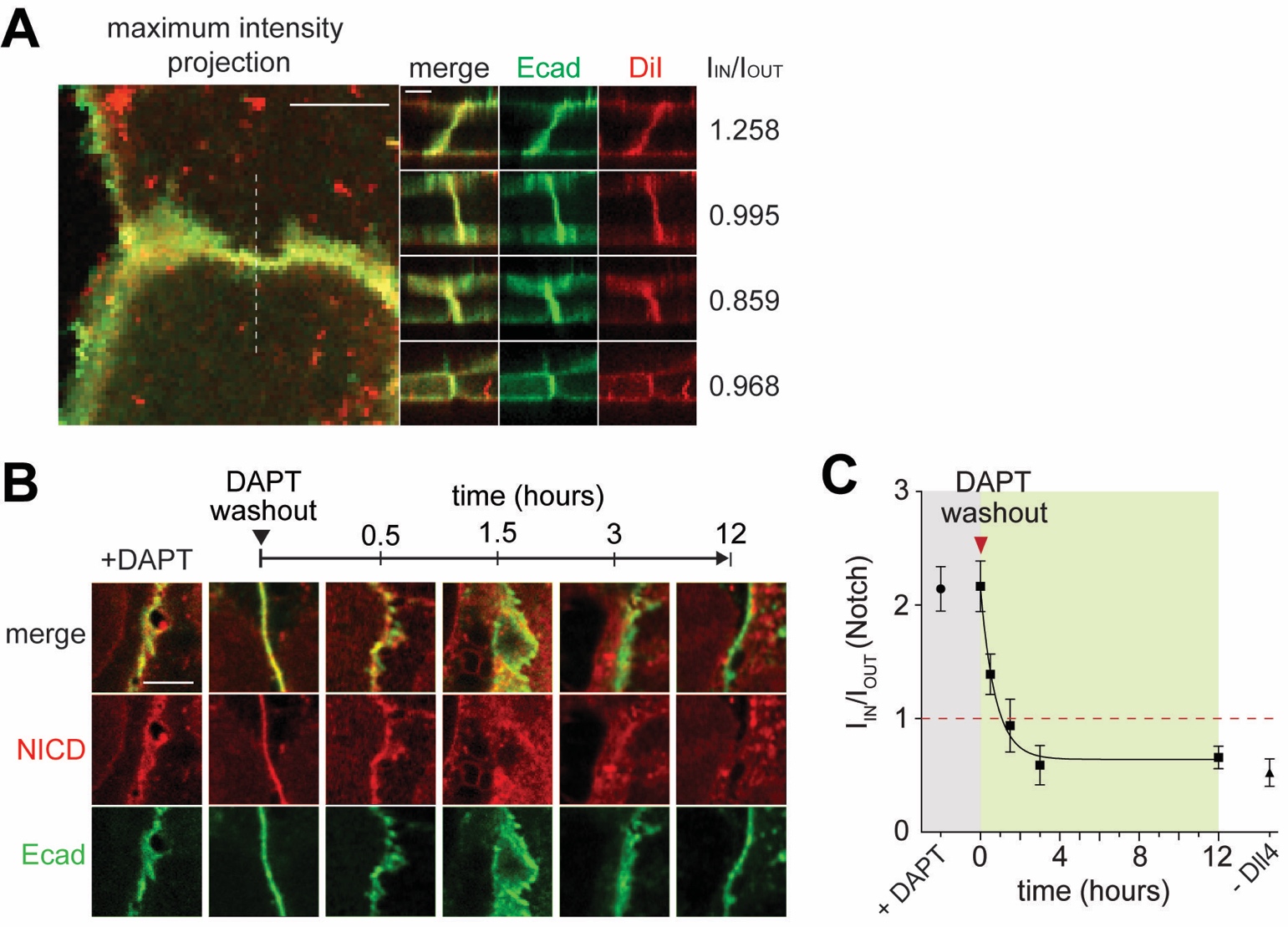

Fig. S9. Confocal images showing the gradual release of NICD from S2-cleaved Notch concentrated within cadAJs.

**(A)** Representative confocal images and enrichment factors (I_IN_/I_OUT_) of Dil membrane staining dye distribution relative to cadAJs. Scale bars, 10 µm (maximum projection), 3 µm (z resliced images). **(B)** Representative time-course confocal z-resliced images showing S2-cleaved Notch at cadAJs as a function of time after DAPT removal. The NICD signal (red) at the cadAJ gradually decreases, indicating NICD release. Images shown here are not from identical cells, but represent a general trend of NICD signal at cadAJs for each time point. Scale bar, 5 µm **(C)** Quantification I_IN_/I_OUT_ ratio as a function of time after DAPT washout.

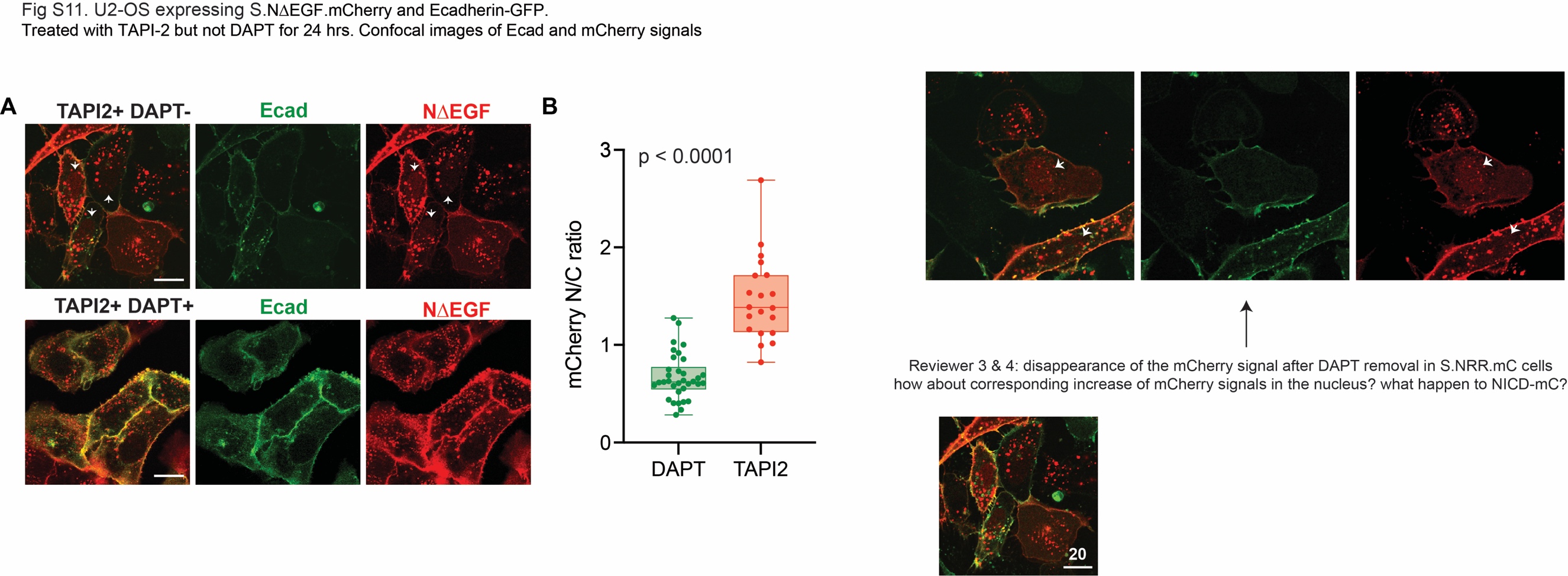

**Fig. S10. Nuclear location of NICD released from cell membrane that recombinantly expresses NΔEGF.** **(A)** Confocal fluorescence images of U2OS cells expressing SNAP-NΔEGF-mCherry and Ecad-GFP. (Upper) Cells treated with TAPI2 only. White arrowheads indicate the cells with nuclear NICD-mCherry accumulation. (Lower) Cells treated with both TAPI2 and DAPT. Scale bar, 20 µm. **(B)** Quantification of the ratio of nucleus-to-cytosolic mCherry signals in cells with DAPT (n = 39) and those without (n = 21) DAPT. The ratio increased from cells without DAPT, suggesting nuclear localization of NICD. Each dot represents the ratio in a single cell. A box and a whisker indicate the interquartile and the full range, respectively. Colored lines indicate median. Two tailed unpaired Student’s t-test.

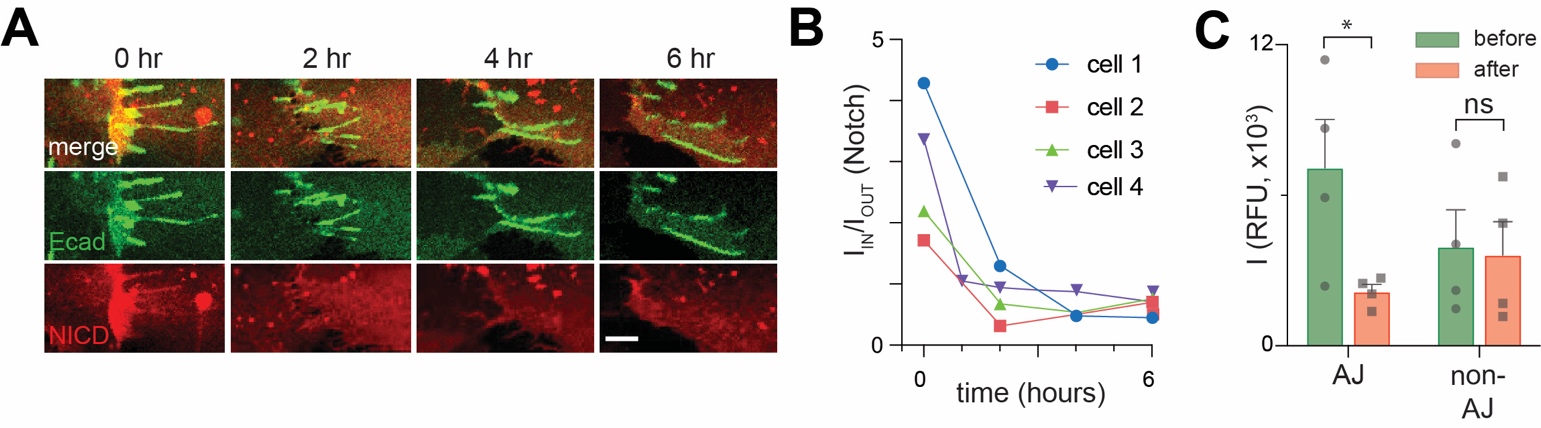

Fig. S11. Confocal images showing the gradual release of NICD from NΔEGF within cadAJs.

**(A-C)** Time traces of NICD release from U2OS cells expressing NΔEGF (red) and Ecad-GFP (green). **(A)** Cells showing strong NΔEGF enrichment at cadAJs under DAPT treatment were chosen. DAPT was removed while keeping TAPI2 concentration constant and time series confocal images were acquired at 0, 2, 4, and 6 hrs from DAPT removal. Scale bar, 5 µm. **(B)** Single-cell traces (n = 4) of enrichment factor. **(C)** Quantification of changes in NICD signal from these four cells at the cadAJs and non-cadAJ membrane, at t = 0 (green, before washout) and t = 6 hr (red, after DAPT washout). CadAJs and non-cadAJ membrane were detected based on thresholding and automatic segmentation using the custom-built script. Average mCherry fluorescence signal from cadAJs and non-cadAJ membranes was also measured using the same script. Intracellular-mCherry signal significantly decreased at the cadAJs, but not at non-cadAJ membranes (*P < 0.05, ns: non-significant, paired two-tailed Student’s t-test, n = 4).

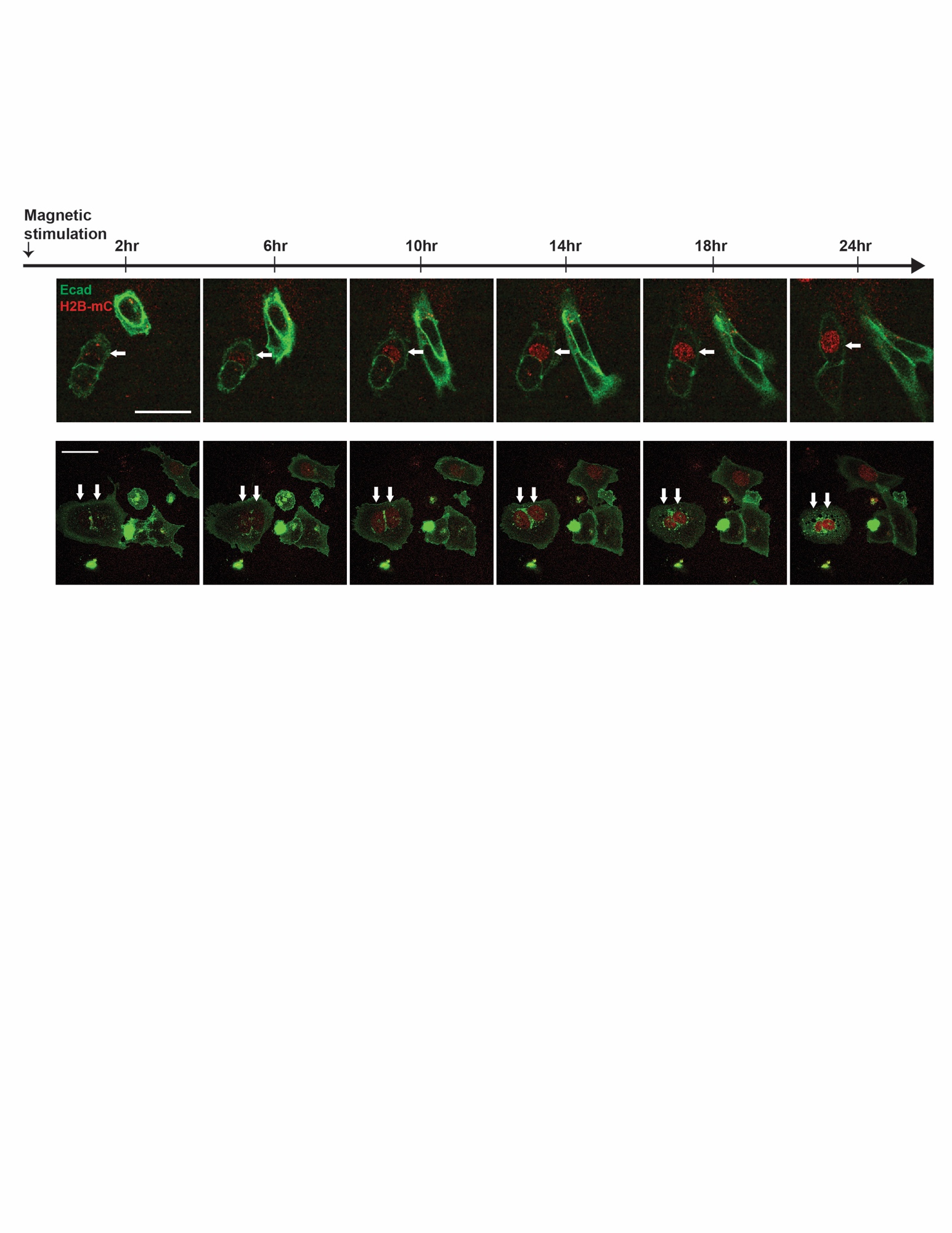

Fig. S12. Colocalization of Notch with artificial cadAJs can stimulate Notch activation in a ligand-independent manner. Representative time-lapse images showing Notch signal activation in UAS-Gal4 reporter cells with artificial cadAJs (white arrows). Cells were cultured in the presence of TAPI2 and no source of S2 cleavage. Neighboring cells without magnetic stimulation were used as internal negative controls. Images were acquired using epifluorescence imaging through FITC and RFP channels (488 nm and 561 nm excitation) every 2 hr for 24 hr. Scale bar, 50 µm.

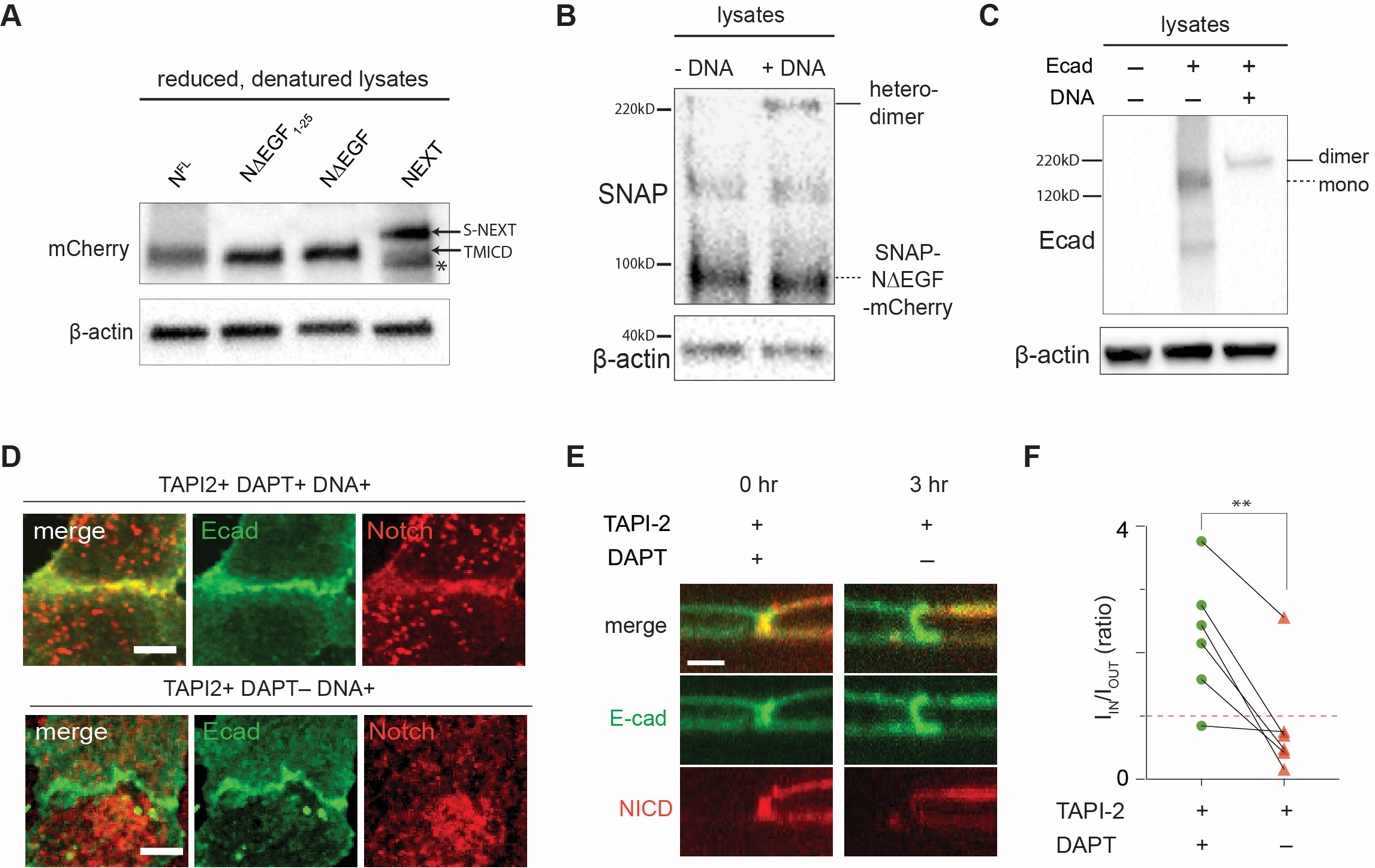

Fig. S13. Spatial mutation of NΔEGF using DNA-mediated crosslinking.

(A) Western blot of cell lysates stably expressing Notch1 truncation variants. The blot was labelled with mCherry antibodies. Cells were treated with TAPI2 and DAPT to inhibit any S2 or S3 cleavage. N^FL^, NΔEGF1-25, and NΔEGF all contain the SDS/DTT-sensitive link (disulfide bridge) that produces the protein band of 78kD corresponding to the polypeptide consisting of small fraction of Notch ECD after S1 site, transmembrane domain (TMD), and intracellular domain (ICD) with mCherry protein (TMICD, black arrow). NΔECD (NEXT) does not have the link, and produces the band corresponding to the intact protein containing SNAP-tag, TMD, and ICD with mCherry protein (S-NEXT, black arrow). Asterisk indicates an unidentified band from S-NEXT. (B and C) A representative western blot of lysate from cells expressing NΔEGF and Halo-Ecad-GFP after 2 hr incubation with or without DNA crosslinkers. The blot was labelled with anti-SNAP (B) and anti-Ecadherin (C) antibodies. The expected mass of NΔEGF, E-cadherin monomer, and the complex with the Notch construct and E-cadherin forming a heterodimer are 90 kD, 158 kD, and 230 kD, respectively. β-actin detection was used to assess protein loading. In both blots, predicted bands representing Notch-E-cadherin heterodimers (solid black lines) and SNAP-NΔEGF-mCherry or Halo-Ecad-GFP monomers (dashed black lines) are indicated. (D) Representative confocal maximum intensity projection images showing the distribution of NΔEGF relative to the cadAJs after crosslinking. Cells were treated with or without DAPT. Scale bar, 10 µm. (E) Single-cell confocal z-resliced images showing intracellular mCherry signal at the cadAJ under DNA and DAPT treatment (left) and after washing out DAPT (right). Removing DAPT elicited a significant reduction in mCherry signal intensity from the cadAJ. Scale bar, 5 µm. (F) Paired analysis of multiple cells expressing NΔEGF (n = 6) in enrichment factor (I_IN_/I_OUT_) after DAPT washout. Each dot represents I_IN_/I_OUT_ value before and after DAPT washout from a single cell. Each line corresponds to the I_IN_/I_OUT_ changes before and after DAPT washout in a same single cell (**P<0.01; paired two-tailed Student’s t test; n = 6).

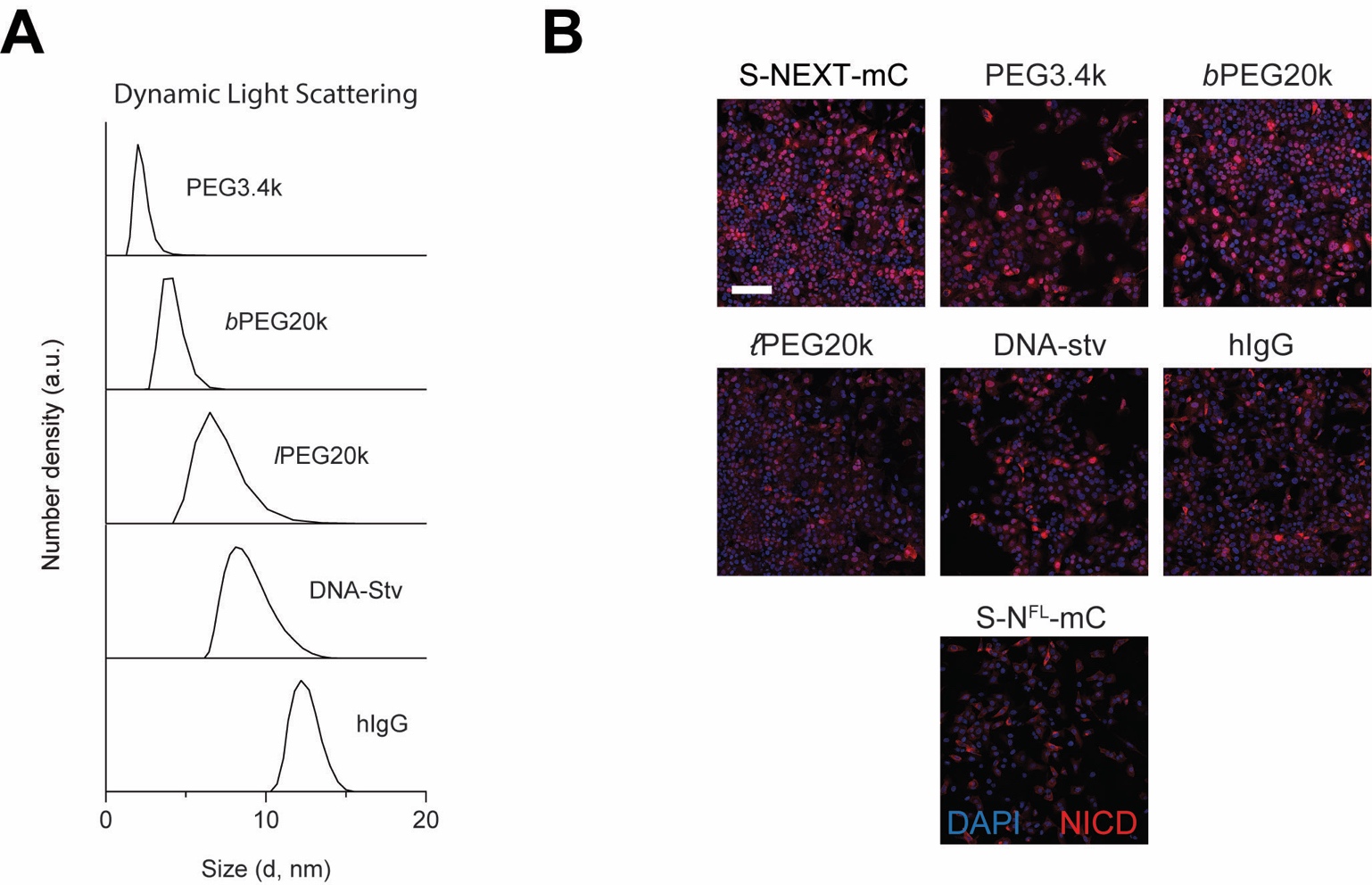

**Fig. S14. Spatial mutation of NEXT using chemical ligation of macromolecular pendants.**

**(A)** Dynamic light scattering spectra of BG-modified macromolecules used in the experiment to induce spatial mutation of NEXT in **Fig. 4, D to G**. **(B)** Larger area (1 x 1 mm^2^) confocal fluorescence images shown in **Fig. 4F**. Scale bar, 200 µm.

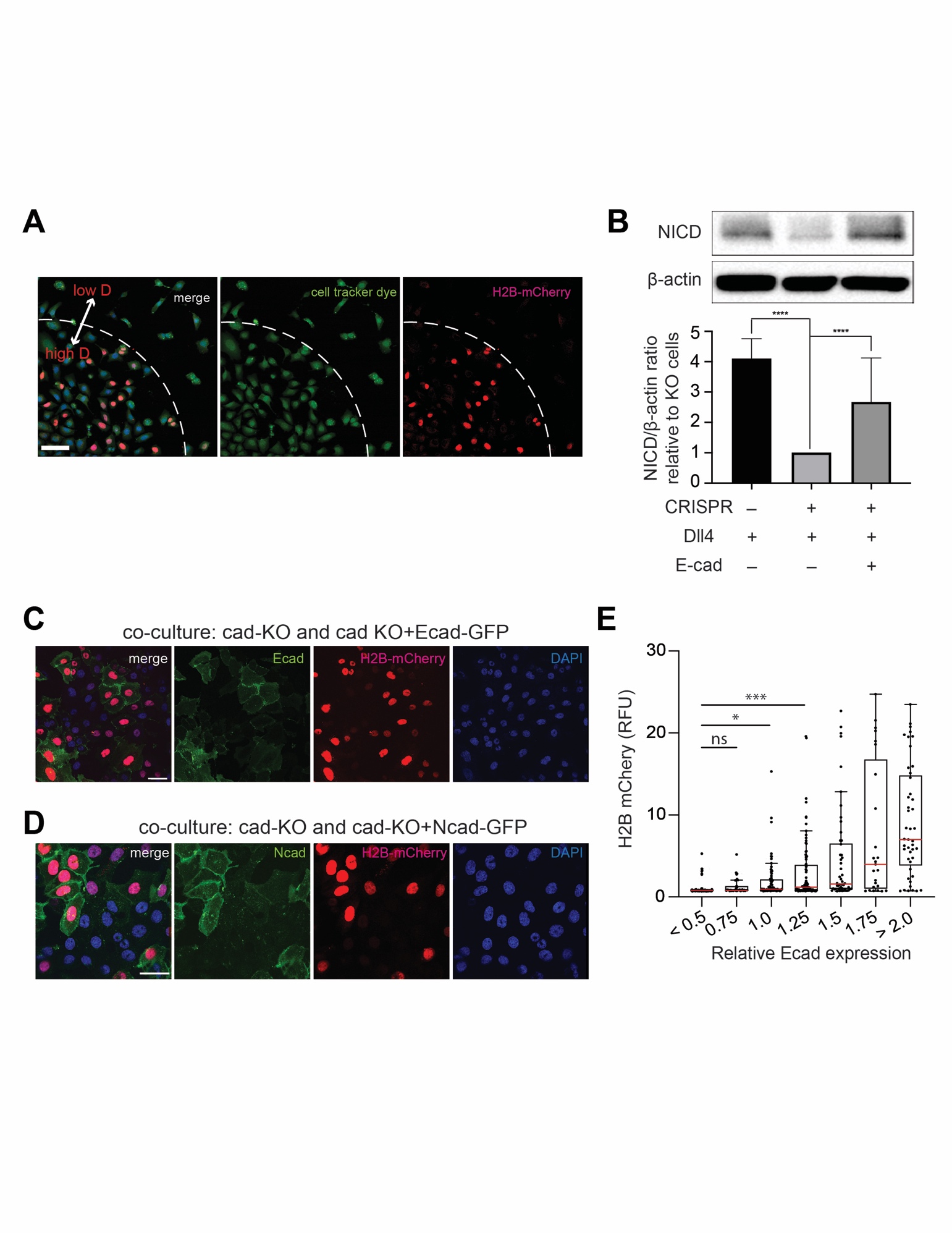

**Fig. S15**. **Reintroduction of E- and N-cadherin into E-cadherin knockout cells rescued Notch activation.** **(A)** Representative low magnification epi-fluorescence image showing both grouped cells and multiple solitary cells. Scale bar, 100 µm. **(B)** Western blot analyses of cleaved NICD levels in the wild-type SNAP-N^FL^-Gal4 cells, CDH1 knock-out (Ecad-KO) cells, and Ecad-KO cells with recombinant E-cadherin transfection. (top) A representative image of immunoblotting. (bottom) Quantification of cleaved NICD levels. The average intensity of NICD bands relative to β-actin bands was quantified and then normalized to that of Ecad-KO cells (mean ± s.d.; ***P < 0.001; n = 5 biological replicates; one-way ANOVA followed by Tukey’s multiple comparison test). **(C and D)** Representative epi-fluorescence images showing Notch activation in co-culture of Ecad-KO cells with Ecad-KO+Ecad cells **(C)** or with Ecad-KO+Ncad cells **(D)**. Ecad-KO cells shows no GFP signal (green) while Ecad-KO+Ecad or Ecad-KO+Ncad cells show robust GFP signal indicative of reintroduction of E- or N-cadherin. Scale bar, 50 µm. **(E)** Comparison of Notch signal activation, readout by mean nuclear H2B-mCherry fluorescence, as a function of E-cadherin expression, readout by membrane GFP fluorescence signal. Each dot represents H2B-mCherry signal of a single cell, and cells are grouped into bins based on their levels of Ecad expression. * p<0.05, *** p<0.001, ns, non-significant. One-way ANOVA followed by Tukey’s test.

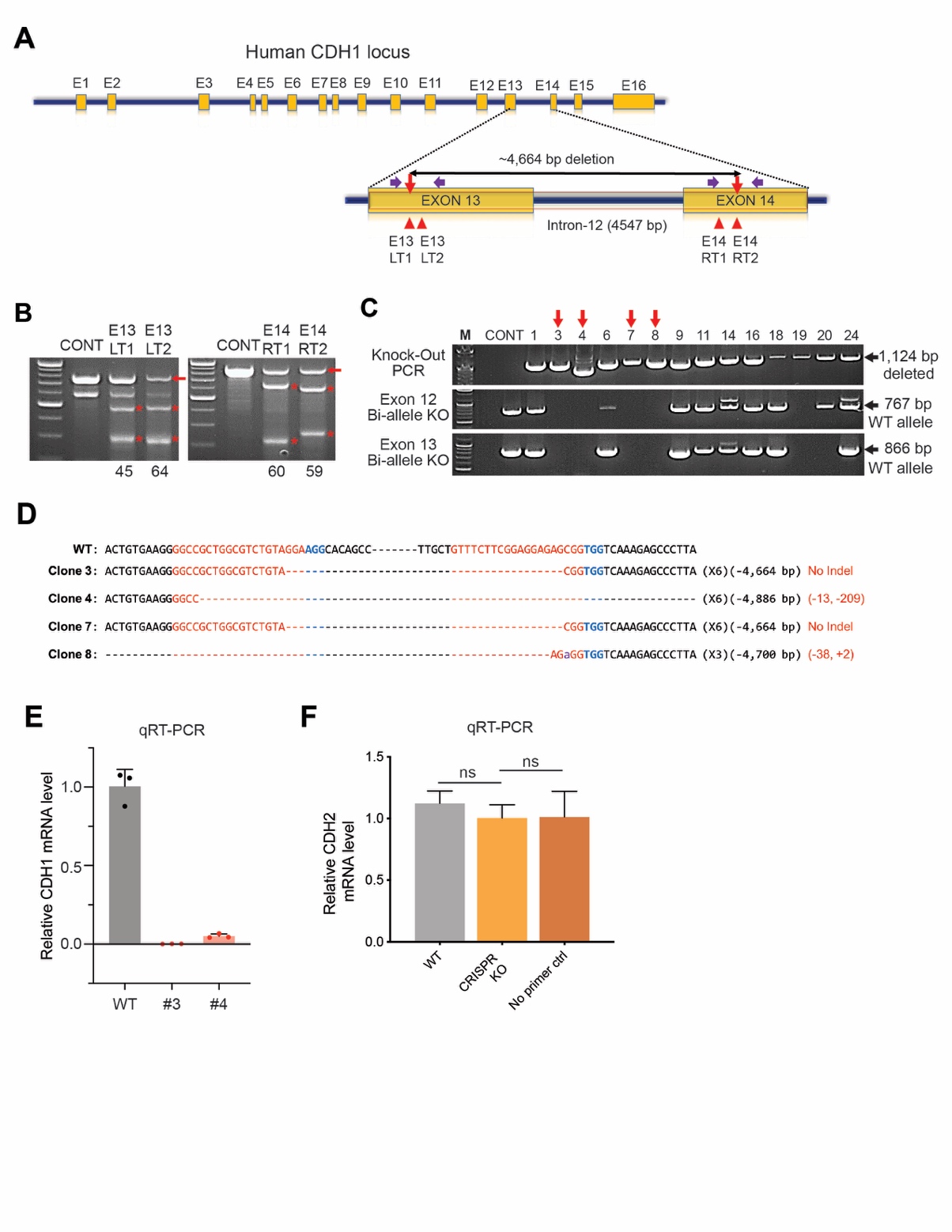

**Fig. S16. Generation of U2OS SNAP-N^FL^-Gal4 fluorescence reporter cell lines lacking E-cadherins via CRISPR/Cas9.**

**(A)** Schematic representation of human CDH1 gene structure and targeted segmental deletion sites. The sixteen exons are shown in orange boxes (E1-E16). Red arrowheads indicate the sgRNA-binding sites (E13LT1, E13LT2, E14RT1 and E14RT2). The targeted segmental deletion of 4.6kb is shown with a black line with red arrow tips, respectively. Purple arrows represent PCR primers used for the T7E1 assay and detection of alleles with targeted deletions, respectively. **(B)** T7E1 assays to evaluate the efficiency of each sgRNAs. U2OS SNAP-N^FL^-Gal4 reporter cells were analyzed after transfection with plasmids encoding Cas9 and each sgRNA for CDH1. Mutation frequencies (indel [%]) were calculated from the band intensities. Untransfected cells were used as controls (CONT). The red arrows and asterisk indicate the uncleaved and expected cleaved position of DNA bands generated by cleavage by T7E1, respectively. M: Marker lane. **(C)** Identification of clones containing the entire segmental deletion in both target alleles for selected clones. The wild-type and deleted alleles were detected by PCR using different primer pairs at exon12 and 13 target regions, respectively. The black arrows indicate the expected positions of the amplicons with wild type allele or allele with deletions. The red arrows indicate the candidate clones containing the entire segmental deletion in both target alleles. **(D)** Sanger sequencing of the CDH1 wild-type (wt) allele and alleles containing deletions of the segmental CDH1 gene. Cas9 recognition sites are shown in red and the protospacer adjacent motif (PAM) sequences are shown in blue bold characters. Dashes indicate deleted bases. The number of occurrences is shown in the first set of parentheses (e.g., X6, X3 and X6 indicate how many times each sequence was observed). The cleavage sites are indicated by red arrows. The sequence length of the consequent large deletion is shown in the second set of parentheses. **(E)** CDH1 mRNA expression levels in CDH1 KO clone #3 and #4 were determined using qRT-PCR. CDH1 expression levels in the selected clones containing a segmental deletion were quantified relative to CDH1 mRNA levels of the wild-type U2OS SNAP-N^FL^-Gal4 cells. Clone #3 was used for subsequent experiments. **(F)** qRT-PCR analysis of CDH2 mRNA expression levels in U2OS SNAP-N^FL^-Gal4 reporter cells (WT), CDH1 KO clone #3 (Ecad-KO), and a negative control sample (no primer pair added). CDH2 mRNA levels in both WT and Ecad-KO cells were quantified relative to the negative control sample. Both WT and Ecad-KO cells showed negligible CDH2 mRNA levels, indicating that Ecad-KO cells have minimal mRNA expression of both CDH1 and CDH2. Mean ± SEM; n = 3; ns, non-significant; one-way ordinary ANOVA test.

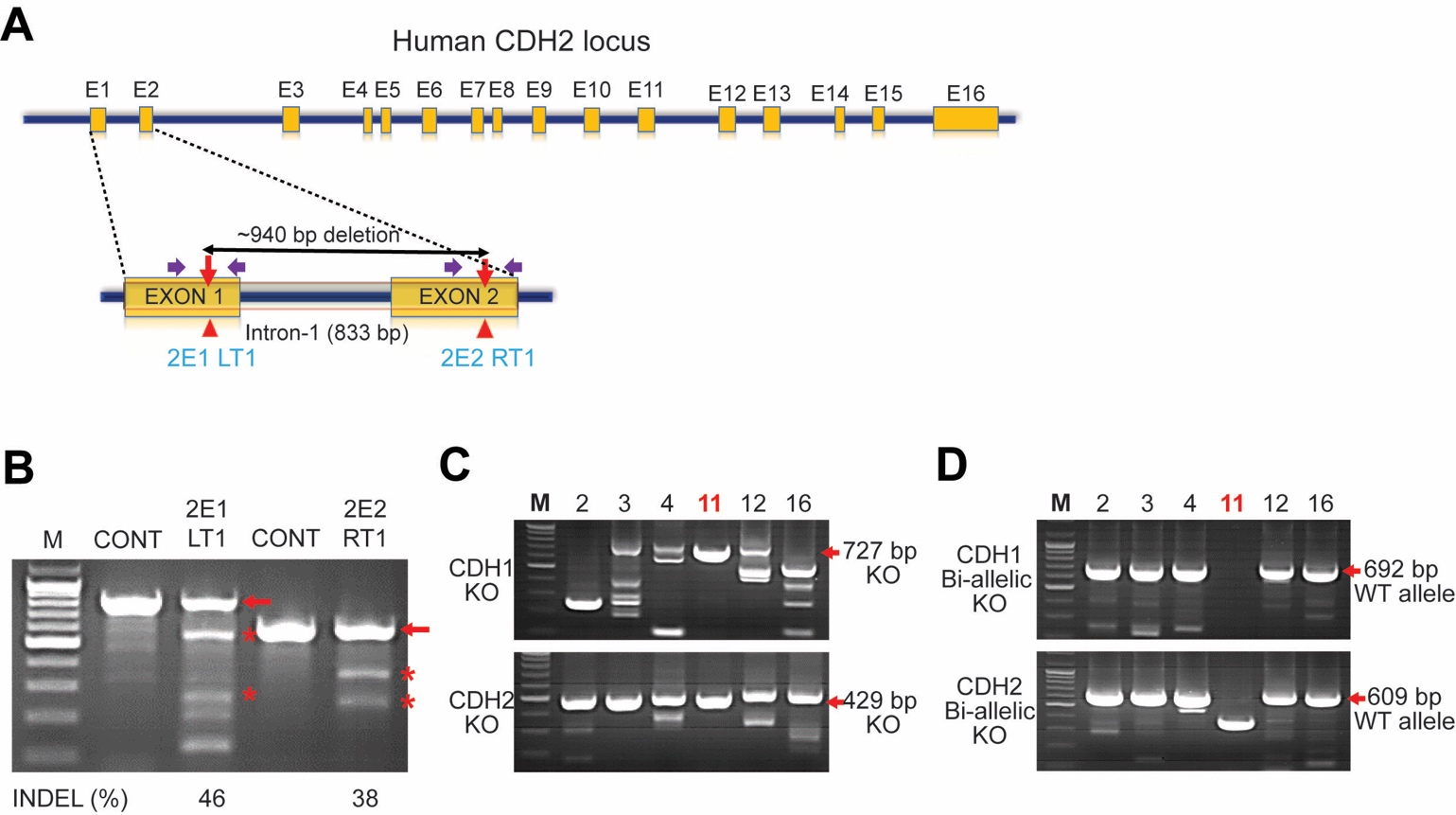

**Fig. S17. Generation of U2OS SNAP-N^FL^-Gal4 fluorescence reporter cell lines lacking E- and N-cadherins via CRISPR/Cas9.**

While CDH2 expression levels in U2OS cells are negligible, we also generated CDH1/2-KO cell lines to eliminate any potential contributions of minimal N-cadherin expression to Aβ generation. **(A)** Schematic representation of human CDH2 gene structure and targeted segmental deletion sites. The sixteen exons are shown in orange boxes (E1-E16). Red arrowheads indicate the sgRNA-binding sites (2E1LT1 and 2E2RT1). The targeted segmental deletion of 940 bp is shown with a black line with red arrow tips, respectively. Purple arrows represent PCR primers used for the T7E1 assay and detection of alleles with targeted deletions, respectively. **(B)** T7E1 assays to evaluate the efficiency of each sgRNAs. U2OS SNAP-N^FL^-Gal4 reporter cells were analyzed after transfection with plasmids encoding Cas9 and each sgRNA for CDH2. Mutation frequencies (indel [%]) were calculated from the band intensities. Untransfected cells were used as controls (CONT). The red arrows and asterisk indicate the uncleaved and expected cleaved position of DNA bands generated by cleavage by T7E1, respectively. M: Marker lane. **(C)** PCR-based evaluation of targeted segmental deletions in double KO (CDH1 and CDH2) single-cell derived clones. U2OS SNAP-N^FL^-Gal4 cells were transfected with plasmids encoding Cas9 and two sets of sgRNA pairs, each targeting CDH1 and CDH2, respectively. The red arrow indicates the expected position of the amplicon from the region containing the segmental targeted deletion M: Marker lane**. (D)** To obtain the bi-allelic KO clones, we validated bi-allelic deletion in both CDH1 and CDH2 genes from the indicated clones. The red arrows indicate the expected positions of the amplicons with wild type allele or allele with deletions. Clone #11 (red color), showing bi-allelic deletion of both CDH1 and CDH2, was used for further subsequent experiments. M: Marker lane**.**

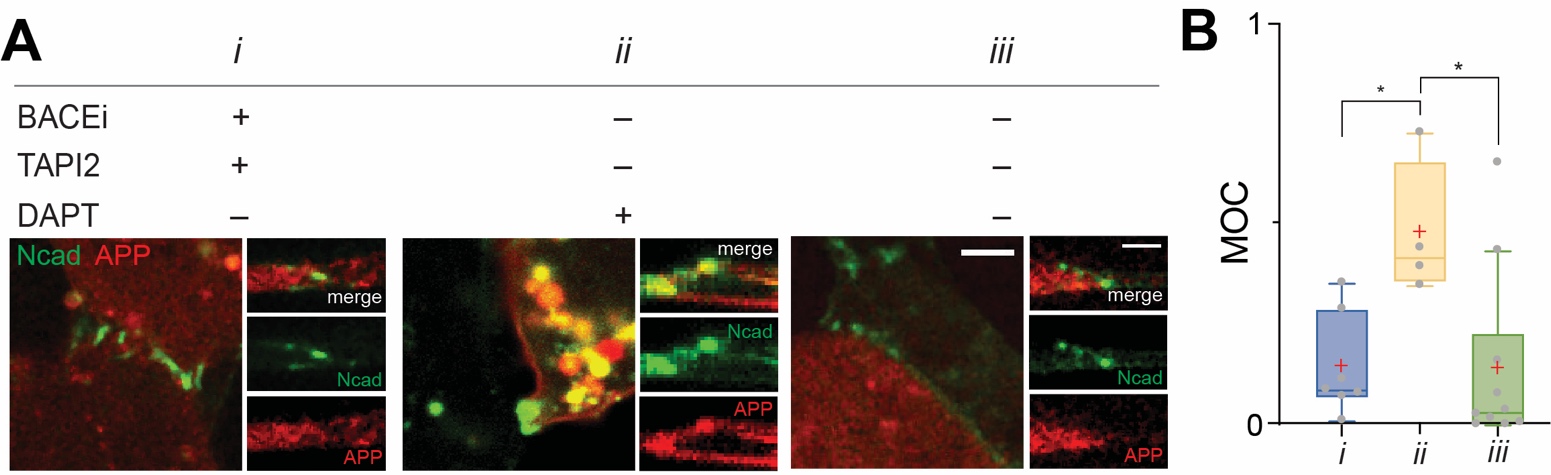

**Fig. S18. Amyloid precursor proteins (APPs) with intact YENPTY motif show size-dependent spatial segregation and membrane proteolysis, consistent with APP lacking the YENPTY motif. (A)** Representative confocal maximum projection (right) and z-resliced (left) images of U2OS cells co-expressing N-cadherin (green) and full-length APP (red). To capture the spatial distribution of the APP intermediates, cells were cultured with a combination of α-, β-, and γ-secretase inhibitors. Scale bar, 3 µm (max. projection) and 2 µm (z-resliced). **(B)** The spatial redistribution of APP relative to the NcadAJs was quantified using Manders’ overlap coefficient (MOC). Data are presented as boxes and whiskers, representing interquartile and min-to-max ranges, respectively; n ≥ 4 NcadAJs, each detected from a single cell.

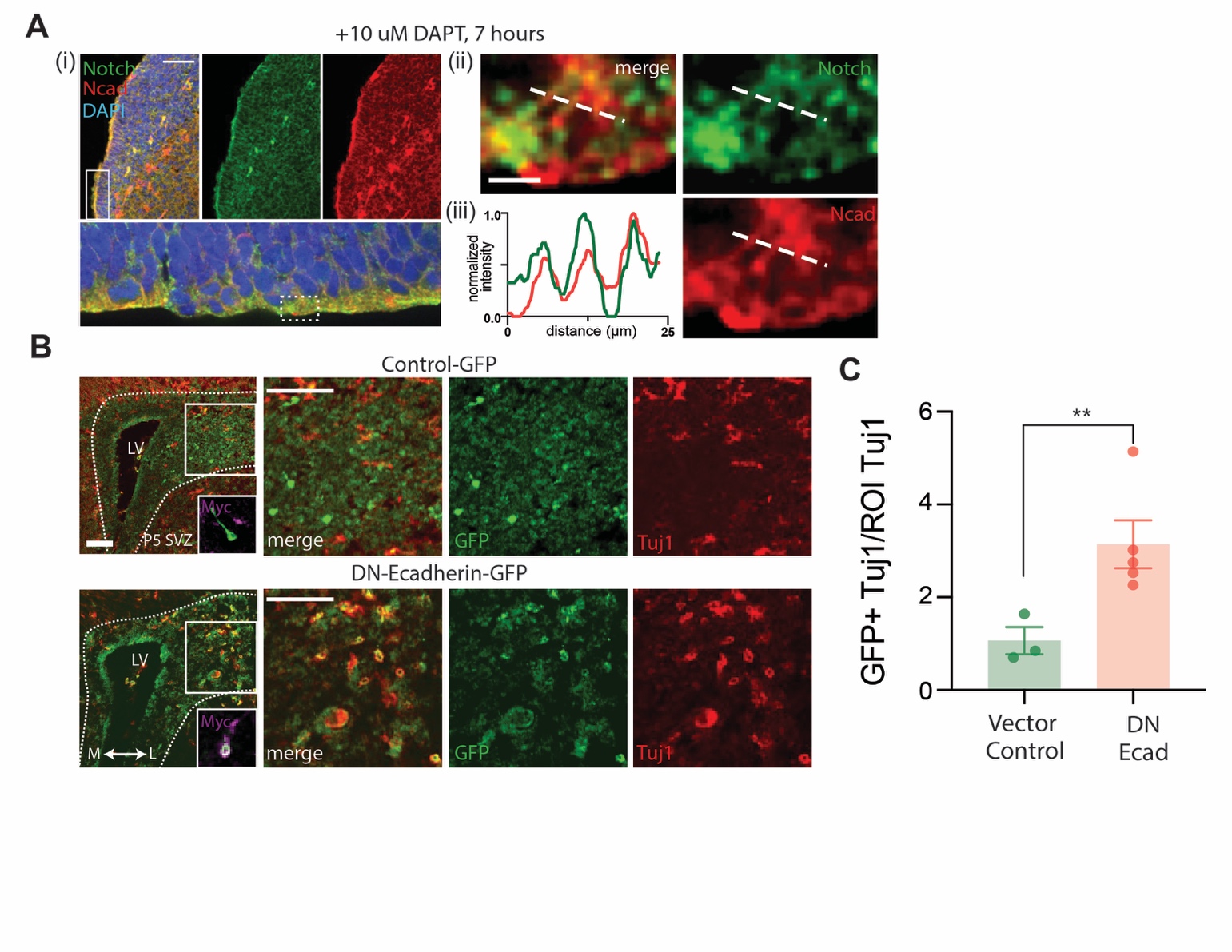

**Fig. S19. Additional immunofluorescence images shown in Figure 6. (A)** Immunostaining of the subventricular zone (SVZ) in the lateral ventricle (LV) of the E13.5 DAPT-treated mouse brain. Notch was colocalized at NcadAJ, visualized by immunostaining with anti-N-cadherin and anti-Notch1 antibodies. (i) Representative lower magnification image. The indicated area (a white box) is magnified and rotated 90º clockwise in the lower panel. Scale bar, 100 µm. (ii) Magnified view of the region indicated with a white dashed box in the (i) lower panel. Scale bar, 2.5 µm. (iii) Line profiles of N-cadherin and Notch distributions. (**B**) Additional confocal images of coronal sections of developing mouse brain retrovirally infected with dominant negative form of E-cadherin vector (DN-Ecad-EGFP). Transduced cells differentiated into post-mitotic neurons can be identified as EGFP+/Tuj1+, while those remained as NPCs with plasmid transfection are only EGFP+. (left) Low-magnification images. Insets show the magnified image of a representative single cell immunostained for myc-tag. (right) Magnified view of the boxed region. Scale bar, 50 µm. **(C)** Ratio of GFP/Tuj1-double positive cells to total Tuj1-positive post-mitotic neurons in these two conditions. Data are represented as mean ± SEM. * p<0.05, two-tailed unpaired Student’s t test.

Table S1. Cell lines and genetic constructs used in this study.

| Cell line | Stable expression*^,^** | CRISPR-KO | Transient expression 1*^,^** | Transient expression 2*^,^** | Figure |
| --- | --- | --- | --- | --- | --- |
| Plain U2OS |  |  | Halo-Dll1 | Ecad-GFP | 1(B, C)  S2 (C, D) |
|  |  |  | Ecad-GFP | Flot1-Halo | S6 (A-E)  S7 (D) |
|  |  |  | SNAP-Ecad-GFP | Flot1-Halo | 1(E, F)  S5 (A, B) |
|  |  |  | Ecad-GFP |  | S1 (A-F)  S8A |
|  |  |  | APPΔY-GFP | SNAP-Ncad | 5(F, G) |
| T-rex U2OS  (doxycycline inducible) | SNAP-N^FL^-mCherry  (N^FL^) |  | Ecad-GFP |  | 1(B, C)  2(B, C)  3(E, F)  4H  S2(A, B)  S4A  S9(B, C)  S13A |
|  |  |  | Halo-Ecad-GFP |  | 1G  4(I, J)  S8(A-F) |
|  | SNAP-NΔEGF_1-25_-mCherry  (NΔEGF_1-25_) |  | Ecad-GFP |  | 2(E, F)  3(E, F)  4H  S13A |
|  | SNAP-NΔEGF-mCherry  (NΔEGF) |  | Ecad-GFP |  | 2(E, F)  3(E, F)  4H  S10(A-C)  S11(A, B)  S13A |
|  |  |  | Halo-Ecad-GFP |  | 4(A-C)  4H  S13(B-F) |
|  | SNAP-NEXT-mCherry  (NEXT) |  | Ecad-GFP |  | 2(E, F)  3(E, F)  4(D-H)  S13A  S14B |
|  | SNAP-N^FL^-Gal4 &  UAS-H2B-mCherry |  |  |  | 4(K, L)  5(A-E)  S12  S15(A, B)  S16F |
|  |  | Ecad | Ecad-GFP |  | 5(D, E)  S15(A, C, E) |
|  |  |  | Ncad-GFP |  | 5(D, E)  S15(A, D) |
|  | SNAP-N^FL^-Gal4 &  UAS-H2B-mCherry  (uninduced) |  | APP-mCherry | Ncad-GFP | S19(A, B) |
|  |  |  | APP-mCherry |  | 5(H-J) |
|  |  | Ecad &  Ncad | APP-mCherry |  | 5(H-J) |
| HACAT |  |  |  |  | S3(A, B) |
| MDCK |  |  | SNAP-N^FL^-mC | Ecad-GFP | S4(B, D) |
| HUVEC |  |  | SNAP-N^FL^-mC |  | S4C |

*X-Y represents a fusion protein of X with Y.

**X-Y-Z represents a Y protein fused with X- and Z- tag proteins at its N- and C-termini, respectively.

S: Supplementary Figures

Abbreviations:

Notch receptors and ligands

N^FL^: full length Notch

NΔEGF_1-25_: Notch with EGF domain truncation from 1-25.

NΔEGF: Notch with complete EGF repeat domain truncation.

NEXT: Notch with extracellular domain truncation

Dll1: Delta like ligand 1

Cadherins

Ecad: epithelial (E) cadherin

Ncad: neural (N) cadherin

VEcad: vascular endothelial cadherin

Amyloid precursor proteins

APPΔY: amyloid precursor proteins with deletion of YENPTY motifs

APP: full-length amyloid precursor proteins

Self-labeling and fluorescent protein tags

SNAP: SNAP tag

Halo: Halo tag

GFP: green fluorescent protein

Others

Flot1: flotillin 1

H2B: histone 2B

UAS: upstream activation sequence

**Movie S1**. Time-lapse of Notch activation in cell groups vs. solitary cells.

Left column: Representative movies of U2OS cells expressing SNAP-N^FL^-Gal4 and UAS-H2B-mC with robust cadAJs within high-density culture. The cells in this group exhibited robust increase in nuclear mCherry fluorescence signal. Right columns: Representative movies of U2OS cells having no prior contact with other cells (n=9). The solitary cells exhibited negligible increase in nuclear mCherry fluorescence signals. Each movie is digitally cropped and centered on the specific representative cells. SNAP-N^FL^-Gal4 expression and mCherry fluorescence are pseudo-colored as green and red, respectively. Time stamps are relative to Dox addition.

Movie S2. Time-lapse Notch activation in solitary cells cultured on a substrate with Ecad-Fc.

Left column: Representative movies of solitary cells cultured on a substrate coated with Ecad-Fc but not with Dll4-Fc (n=9). Negligible increase of mCherry fluorescence signal was observed. Right column: Representative movies of solitary cells cultured on a substrate coated with Ecad-Fc and Dll4-Fc (n=9). Gradual increase of bright mCherry fluorescence signal was observed. SNAP-N^FL^-Gal4 expression and mCherry fluorescence are pseudo-colored as green and red, respectively. Time stamps are relative to Dox addition.
